# Quantitative imaging of species-specific lipid transport in mammalian cells

**DOI:** 10.1101/2024.05.14.594078

**Authors:** Juan M. Iglesias-Artola, Kai Schuhmann, Kristin Böhlig, H. Mathilda Lennartz, Milena Schuhmacher, Pavel Barahtjan, Cristina Jiménez López, Radek Šachl, Karina Pombo-Garcia, Annett Lohmann, Petra Riegerová, Martin Hof, Björn Drobot, Andrej Shevchenko, Alf Honigmann, André Nadler

**Author notes:** These authors contributed equally: Kai Schuhmann and Kristin Böhlig.

## Abstract

Eukaryotic cells produce over 1000 different lipid species which tune organelle membrane properties, control signalling and store energy^1,2^. How lipid species are selectively sorted between organelles to maintain specific membrane identities is largely unknown due to the difficulty to image lipid transport in cells^3^. Here, we measured transport and metabolism of individual lipid species in mammalian cells using time-resolved fluorescence imaging of bifunctional lipid probes in combination with ultra-high resolution mass spectrometry and mathematical modelling. Quantification of lipid flux between organelles revealed that directional, non-vesicular lipid transport is responsible for fast, species-selective lipid sorting compared to slow, unspecific vesicular membrane trafficking. Using genetic perturbations, we found that coupling between active lipid flipping and passive non-vesicular transport is a mechanism for directional lipid transport. Comparison of metabolic conversion and transport rates showed that non-vesicular transport dominates the organelle distribution of lipids while species-specific phospholipid metabolism controls neutral lipid accumulation. Our results provide the first quantitative map of retrograde lipid flux in cells^4^. We anticipate that our pipeline for quantitative mapping of lipid flux through physical and chemical space in cells will boost our understanding of lipids in cell biology and disease.

## Main

Eukaryotic cells produce a plethora of chemically distinct lipid species with varying side chain unsaturation, length and regiochemistry which belong to dozens of lipid classes defined by lipid headgroup and backbone^1^. Lipid species are differentially distributed across organelle membranes, which is important to establish organelle identities and functions^2,3,5,6^. How the organelle specific distribution of lipids is established and maintained is incompletely understood^3^.

Lipid biosynthesis occurs mostly in the endoplasmic reticulum (ER) and lipids are subsequently distributed via vesicular trafficking and membrane contact sites (MCS) to other organelles^2,7,8^. During anterograde lipid transport towards the plasma membrane (PM) lipids are further modified, before they are either recycled via the retrograde pathway to the ER or catabolized in lysosomes, peroxisomes and mitochondria. Understanding how the highly dynamic interplay between local metabolism and inter-organelle transport gives rise to distinct organelle identities requires quantitative measurements of intracellular lipid transport kinetics and local metabolism on the lipid species level. While anterograde lipid flux from the ER to the PM has been characterized for some lipid classes using metabolic labelling and organelle fractionation^3^, the trafficking of individual lipid species in particular in the retrograde lipid transport pathway is not well understood, with the notable exception of Sphingomyelin (SM)^9^. So far, one of the key limitations has been that distinct lipid species could not be faithfully imaged using fluorescent microscopy, hindering the analysis of spatiotemporal transport dynamics. Here we used minimally modified lipid probes, ultra-high-resolution Fourier-Transform (FT) mass spectrometry, fluorescence imaging and mathematical modelling to quantitatively map the kinetics of species-specific lipid transport and metabolism, identify the primary mechanism of lipid sorting into organelle membranes and build a publicly accessible lipid flux atlas available at: http://doi.org/21.11101/0000-0007-FCE5-B

### Fluorescence imaging of species-specific lipid transport

To quantify the kinetics of transport and metabolism of individual lipid species in mammalian cells, we leveraged photoactivatable and clickable (bifunctional) lipids ^10–15^ via a combination of pulse-chase fluorescence imaging with ultra-high resolution mass spectrometry and mathematical modelling. In contrast to other lipid probes which are optimized to either modulate lipid levels (photo-caged lipids^16–18^, photoswitchable lipids^19,20^), to visualize lipid localization (lipid-fluorophore conjugates^21,22^) or to monitor lipid metabolism (isotope-labelled lipids^23,24^, clickable lipids^25–27^), bifunctional lipids allow monitoring both lipid transport and metabolism with the same probe.

To make distinct lipid species accessible for high resolution fluorescence imaging and mass spectrometry, we relied on two minimal modifications (diazirine and alkyne) within the lipid alkyl chain (Fig. 1a). We generated a library of bifunctional lipid probes covering four different lipid classes: Phosphatidylcholine (PC), phosphatidic acid (PA), phosphatidylethanolamine (PE) and sphingomyelin (SM). Within the PC class we varied acyl chain length and unsaturation degree as well as *sn-1*/*sn-2* acyl chain positioning to generate regioisomers (Fig. 1b, see Supplementary Information for synthetic details). We confirmed that incorporation of bifunctional acyl chains did not alter lipid specific membrane properties such as phase behaviour and nanodomain formation in model membranes using FRET assays^28,29^ (Extended Data 1).

**Figure 1.**
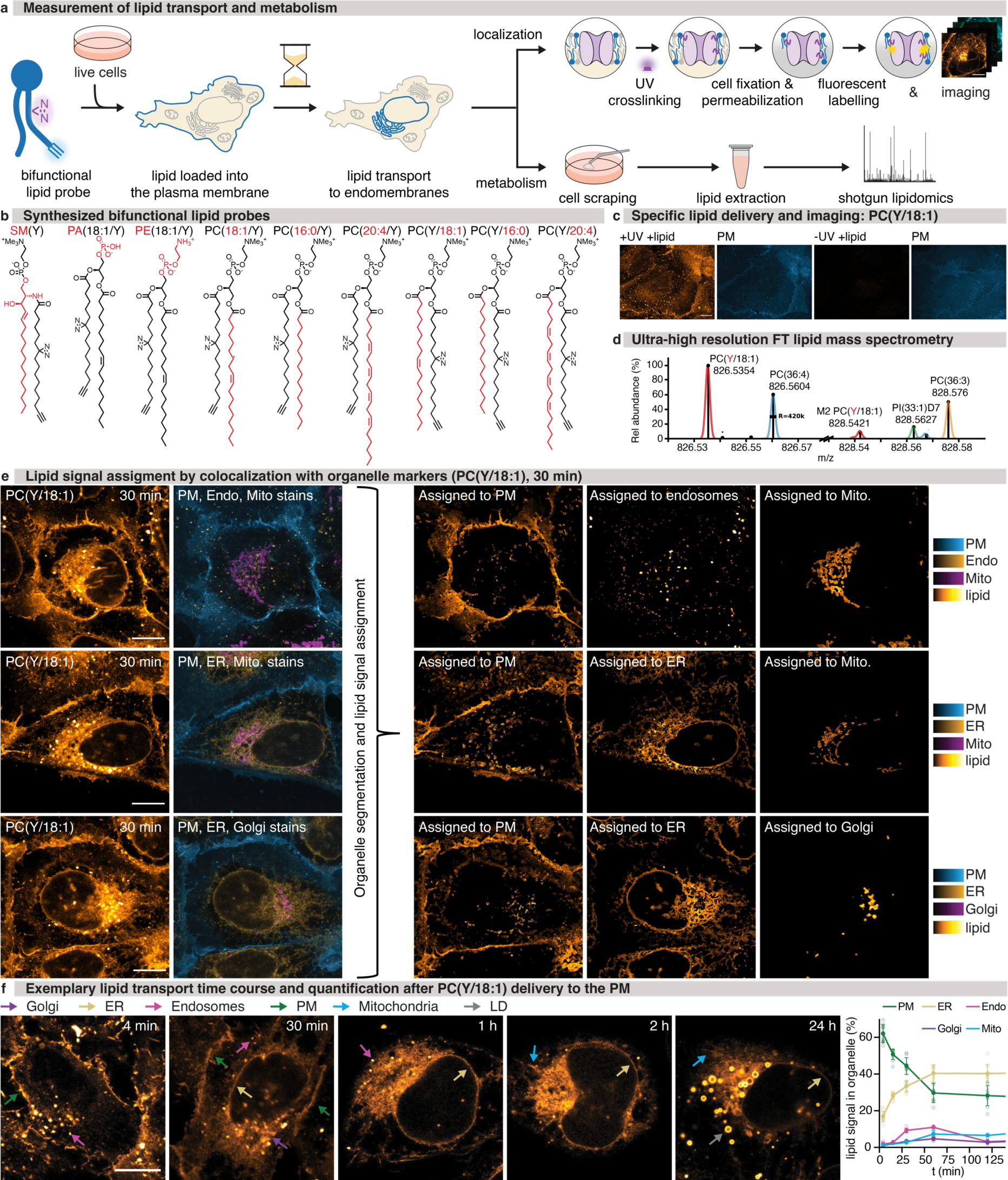
Lipid probe library, imaging and MS work-flows, and lipid transport time course experiments. **a.** Schematic description of combined analysis of lipid transport and metabolism. Lipid probes were loaded into the plasma membrane using alpha-methyl-cyclodextrin mediated exchange reactions, crosslinked and fluorescently labelled for imaging or extracted and analysed by mass spectrometry to monitor metabolism. **b.** Bifunctional lipid probes synthesized for this study. Unique structural elements are highlighted in red. **c.** Lipid delivery to the plasma membrane and selectivity of lipid labelling shown for PC(Y/18:1). Scale bar: 10 μm. **d.** Ultra-high mass resolution (Rs _m/z= 800_ = 420 000) enables base line separation of peaks spaced by a few mDa and their unequivocal assignment to molecular ions of lipids (as annotated; [M-H]^−^/[M+HCO2]^−^) in total lipid extract. M2: 2^nd^ isotopic peak; PI(33:1)D7: deuterated internal standard). **e.** Representative images (PC(Y/18:1), 30 min timepoint) showing lipid signal assignment to individual organelles via four-colour fluorescence imaging and automated image segmentation. Scale bar: 10 μm. Lipid signal images are shown at identical settings. **f.** Left panels: Representative images from time-course experiments show the temporal development of lipid signal distribution for PC(Y/18:1). Coloured arrows indicate lipid localization in different organelles (green: plasma membrane, yellow: endoplasmic reticulum, cyan: mitochondria, violet: Golgi apparatus, magenta: endosomes grey: lipid droplets). Scale bar: 10 μm. Images are brightness-contrast adjusted to allow for comparing lipid distributions at different timepoints. Right panel: Quantification of temporal development of intracellular lipid distribution for PC(Y/18:1). Error bars: SD, each datapoint is derived from 3-15 z-stack images containing 5-10 cells per stack.

Studying lipid transport from the plasma membrane to internal membranes requires a well-defined starting point to chase retrograde transport into the cell. We thus incorporated lipid probes into the outer leaflet of the plasma membrane of U-2 OS cells via a 0.5-4 min loading pulse of alpha-methyl-cyclodextrin mediated lipid exchange from donor liposomes (Figure 1a). Plasma membrane integrity was not affected by the loading process (Extended Data 2) and quantification using mass spectrometry showed that approximately 1 – 3% of the total cellular lipidome was exchanged with bifunctional lipid probes while the overall lipidome composition, including cholesterol content, remained essentially unaffected (Extended Data 1, 5, 8).

After the loading pulse, cells were kept at 37 °C for 0 min - 24 h prior to lipid photo-crosslinking, cell fixation, removal of non-crosslinked material and fluorescence labelling via click chemistry (Fig. 1a, see Supplementary Information for details). The transport of bifunctional lipids was analysed by confocal imaging of the photo-crosslinked and fluorescently labelled lipids at all time points (Fig. 1a). Control samples without lipid probes or UV irradiation showed very low unspecific background labelling (Fig. 1c, Extended Data 1,2). Lipid imaging was complemented with quantitative shotgun lipidomics by ultra-high resolution Fourier Transform (FT) Mass Spectrometry for each timepoint to quantify the metabolic conversions during the transport. To this end, we used the mass difference between the two nitrogen atoms of the diazirine functional group (28.0061 Da) and two CH_2_ (28.0313 Da) groups to distinguish bifunctional lipids from their native counterparts (Fig. 1d, ED5).

To quantify the temporal development of intracellular lipid distribution, we assigned the lipid fluorescent signal to distinct organelle membranes, by determining the colocalization of lipids with organelle markers for plasma membrane, Golgi apparatus, endoplasmic reticulum, endosomes and mitochondria (Fig. 1e,f, 2c, Extended Data 3,4, see Supplementary Information for details). Segmented probability maps were generated for every organelle marker using the pixel classifier approach of the Ilastik software package^30^. We then retrieved the organelle specific lipid signal intensity distributions from pixels that were unambiguously assigned to one organelle. Based on these distributions, lipid signal was partitioned between organelles in regions where organelle masks overlapped (Fig. 1e Extended Data 4, see Supplementary Information for details). Taken together, we developed a lipid imaging pipeline that enables quantification of the inter-organelle transport of distinct lipid species starting from the plasma membrane and correlation of lipid transport with time dependent metabolic conversion of lipids observed by mass spectrometry.

### Lipid species-specific retrograde transport occurs via non-vesicular routes

Visual inspection of the lipid localization in confocal images revealed clear differences in transport kinetics between the lipid classes, and even between individual species within the same lipid class (Fig. 2c and Extended Data 3,4). Polyunsaturated PC species were rapidly transported into the endo-membrane system after 4 minutes. In contrast, noticeable transport to the ER took up to one hour for saturated PCs. Overall, poly-unsaturated PC species, phosphatidic acid (PA) and phosphatidylethanolamine (PE) exhibited a pronounced early localization in the ER, whereas saturated PC species and SM were retained much longer in the plasma membrane (PM) and subsequently showed persistent localization in endosomes (Fig. 2c, Extended Data 3, 4). These observations indicated that the kinetics of intracellular lipid transport differ both on the level of lipid classes and individual lipid species.

**Figure 2.**
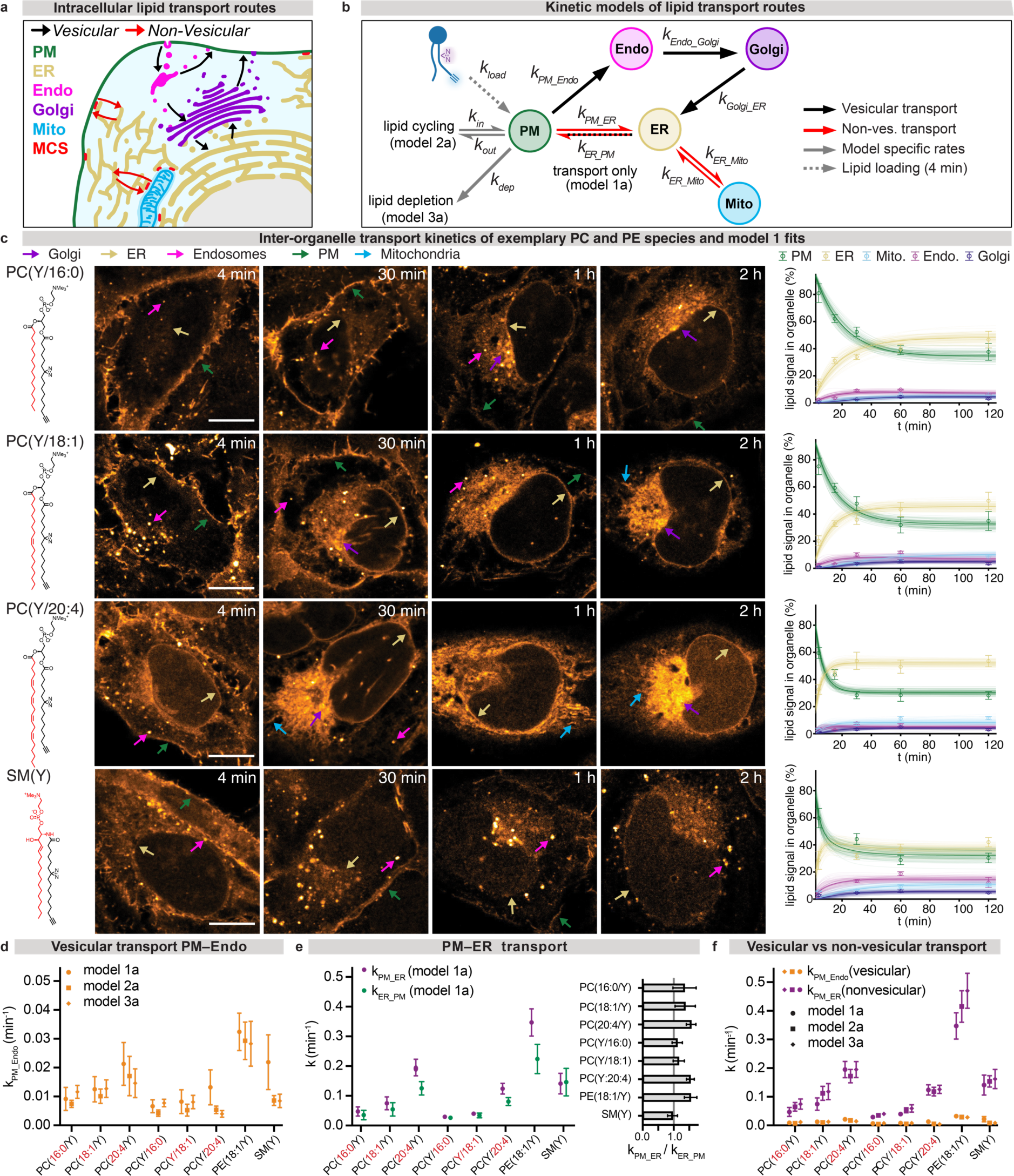
Retrograde lipid transport occurs primarily via non-vesicular routes. **a.** Schematic representation of the analysed cellular lipid transport pipelines. **b.** Kinetic models for quantifying lipid transport from fluorescence microscopy and mass spectrometry data. **c.** Kinetics of lipid transport exemplarily shown for PC(Y/16:0), PC(Y/18:1), PC(Y/20:4), SM(Y) and corresponding model 1a fits. Unique structural elements are highlighted in red. Scale bar: 10 μm. Images are brightness-contrast adjusted to allow for comparing lipid distributions at different timepoints. **d.** Comparison of rate constants describing retrograde vesicular transport from the PM to endosomes (models 1a-3a shown). **e.** Comparison of rate constants describing retrograde non-vesicular transport from the PM to the ER and total transport in the anterograde direction. **f.** Comparison of rate constants describing retrograde vesicular transport from the PM to endosomes and retrograde non-vesicular transport from the PM to the ER for all analysed lipid probes. Error bars, image quantification: SD each datapoint is derived from 3-15 z-stack images containing 5-10 cells per stack. Error bars, rate constants: SD, calculated from 100 MC model runs.

To understand whether the observed transport selectivity arises from differential sorting of lipid species during vesicular or non-vesicular transport (Fig. 2a), we fitted a kinetic model describing the main lipid transport routes to the lipid transport data (Fig. 2b). The model included vesicular transport via endocytosis from the PM, into endosomes and the Golgi apparatus to the ER and the competing, non-vesicular route from the PM to the ER as well as lipid exchange between the ER and mitochondria (Fig. 2b,c, Extended Data 6,7, see Supplementary Information for details). Kinetic models were fitted globally for each lipid species, except for PA(18:1/Y) which was transported too fast for the time resolution of the time-course experiments, to obtain inter-organelle transport rate constants (Fig. 2d). To assess the robustness of derived kinetic parameters, we compared different model versions featuring lipid transport networks of increasing complexity and accounting for quantitative bifunctional lipid content derived from mass spectrometry (Fig. 2b, Extended Data 5, 6, 7, see Supplementary Information for details). The obtained rate constants between different models were very similar, indicating robustness of the results (Fig. 2d-f, Extended Data 6, 7, Supplementary Information).

Comparison of the lipid transport rate constants for vesicular transport via endosomes with non-vesicular transport to the ER revealed that non-vesicular trafficking was up to an order of magnitude faster for all lipids compared to vesicular transport (Fig. 2f). Furthermore, the rate constants of non-vesicular trafficking showed significant variation between lipid classes and species (Fig. 2e,f). The fastest non-vesicular retrograde transport was found for PE, followed by polyunsaturated PC species and SM, while transport of saturated PC species was comparatively slow.

To determine the structural determinants of lipid species selective non-vesicular transport, we compared the rate constants obtained for six different PC species. Polyunsaturated PC species were transported up to 10-fold faster via the non-vesicular route than saturated PC species, whereas PC species bearing the bifunctional fatty acid at the *sn-2* position were transported up to two-fold faster than the corresponding regioisomers featuring the bifunctional fatty acid at the *sn-1* position (Fig. 2f). These findings imply that while both unsaturation degree and acyl chain positioning influence the rates of non-vesicular lipid transport, unsaturation degree appeared to be the primary discriminating structural feature.

In contrast to the remarkable selectivity observed during non-vesicular transport, differences between the transport rate constants of the same PC species were significantly smaller in the vesicular endosomal transport pathway and followed no obvious trends (Figure 2d, Extended Data 7). Rate constants describing individual steps of vesicular transport in the anterograde direction (from ER via Golgi and Endosomes to the PM, models 1b-3b) were less well identified. Characterizing these rates more precisely would presumably require delivering lipid probes to the ER and monitoring transport to the PM. (See Table S1 in the Supplementary Information for rate constant overview). Taken together, we find that retrograde non-vesicular lipid transport is both faster and more selective than vesicular transport.

### Lipid flippase knockdown mediates directionality of non-vesicular lipid transport

Next, we assessed the implications of the predominant non-vesicular lipid transport for the steady state lipid distributions between organelles. The highest fraction of lipid signal in the PM versus the ER at steady state was found for SM, followed by the saturated PC(16:0/Y) species, whereas polyunsaturated PC and PE localized preferentially to the ER (Figure 2c, Extended Data 6, c-e, upper panels). An analysis of quasi-equilibrium constants 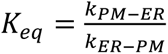 for lipid exchange between PM and ER gave a very similar result (Figure 2e, left panel). These findings are in line with the known lipid concentration gradients between organelles established by membrane fractionation^3^ and imply directional, non-vesicular lipid transport in cells. How different PC species, SM and PE can be directionally transported via membrane contact sites is not well understood. While some lipid transfer proteins can move cholesterol, PA and PS against concentration gradients by PI(4)P_2_ co-transport^8,31^, it is unclear which process provides the energy for directional transport of other lipids, in particular via bridge-like lipid transfer proteins^32,33^.

One attractive mechanism could be the coupling of passive non-vesicular transport to active trans-bilayer flipping of lipids between membrane leaflets catalysed by P4-ATPases, either directly^8^ or indirectly by the utilization of the transmembrane lipid concentration gradient by scramblases^34–36^. To test this hypothesis, we studied the role of lipid leaflet flipping on the transport of PE from the PM to the ER, which is known to be enriched in the inner leaflet of the PM^37^ (Fig. 3a). We genetically knocked down TMEM30A (Extended Data 7j,k), which is the common subunit of plasma membrane flippases that move aminophospholipids to the inner PM leaflet^38,39^ (Fig. 3a) in HCT116 cells, which were chosen for genetic manipulation as they feature a much more intact genome compared to other cancer cell lines. Quantification of PE(18:1/Y) transport between the PM and the ER revealed that PE was transported 3-fold slower in KD cells compared to WT. KD cells had a significantly lower 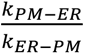 ratio (1.6±0.1 vs 2.2±0.1), indicating an altered steady state distribution, with PE being more strongly enriched in the PM when lipid flipping is perturbed. These results provide direct evidence that ATP-dependent lipid flipping and non-vesicular transport of PE from the PM to the ER are coupled.

**Figure 3.**
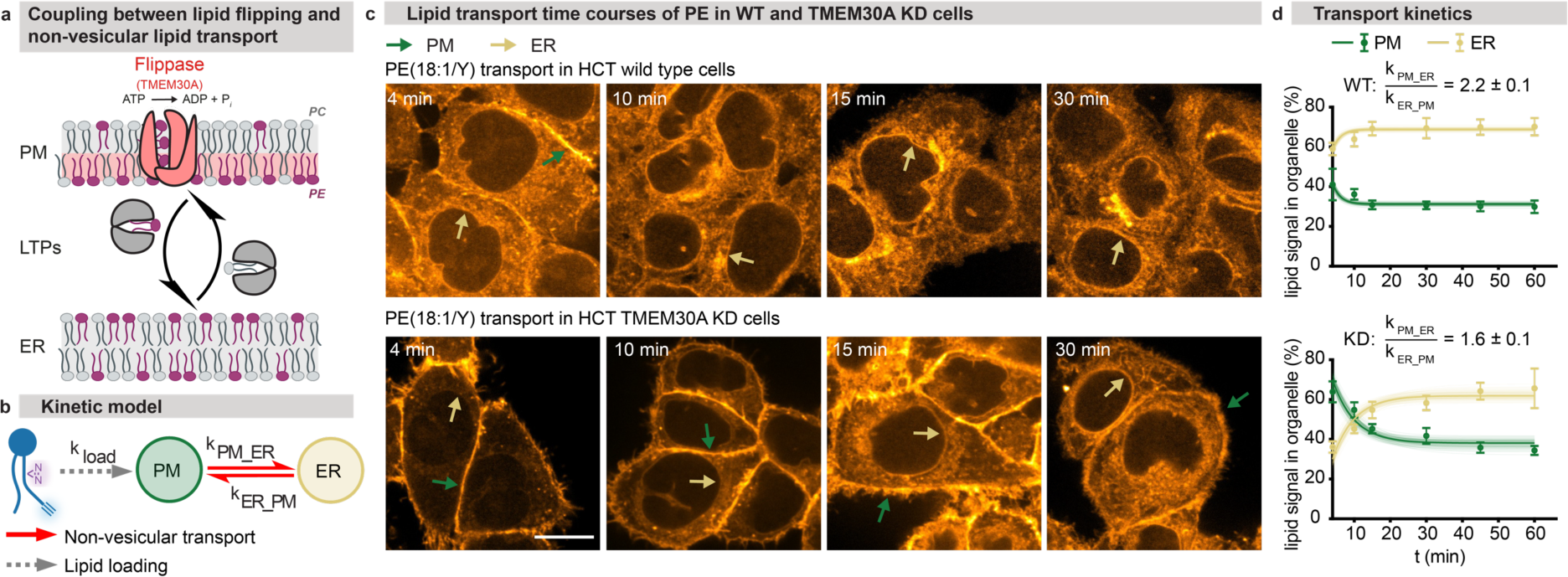
Genetic perturbation experiments confirm involvement of flippases in directional lipid transport. **a.** Schematic representation of lipid trans-bilayer movement (lipid flipping) and non-vesicular lipid transport by lipid transfer proteins. **b.** Kinetic model for exchange of lipids between the PM and the ER. **c.** Comparison of time-course experiments for PE(18:1/Y) show that lipid internalization dynamics are slower in HCT116 TMEM30A KD cells compared to HCT116 wild type cells. Coloured arrows indicate lipid localization in different membranes types (green: PM, yellow: ER). Scale bars: 10 μm. Images are brightness-contrast adjusted to facilitate comparing intracellular lipid localization. **d.** Quantification of lipid internalization kinetics and model fits. Error bars, image quantification: SD, each datapoint is derived from 3-15 z-stack images containing 5-10 cells per stack. Errors, quasi-equilibrium constants: SD, calculated from 100 MC model runs.

### Lipid transport is up to 60 times faster than metabolic lipid conversion

To assess the relative contributions of lipid metabolism and lipid transport processes to lipid sorting into organelles, we next compared transport kinetics to lipid conversion kinetics. We monitored turnover of bifunctional lipid probes by ultra-high resolution FT lipid mass spectrometry (Extended Data 8-10, see lipid metabolism annex for in depth discussion of species-specific metabolic conversion). To obtain a measure of global lipid metabolism, we determined how fast the bifunctional acyl chain of a respective lipid species is redistributed to other lipids by calculating the fraction of the initially supplied bifunctional species with respect to the total abundance of lipids carrying the bifunctional fatty acid (Fig. 4a). To derive the apparent conversion rates, we fitted a mono-exponential model to this dataset. We found an order of magnitude difference between the apparent conversion rate constants of lipid species with values ranging from k_met_ = 0.001 to k_met_ = 0.015 min^−1^ (Fig. 4a,b Extended Data 8a-c).

**Figure 4.**
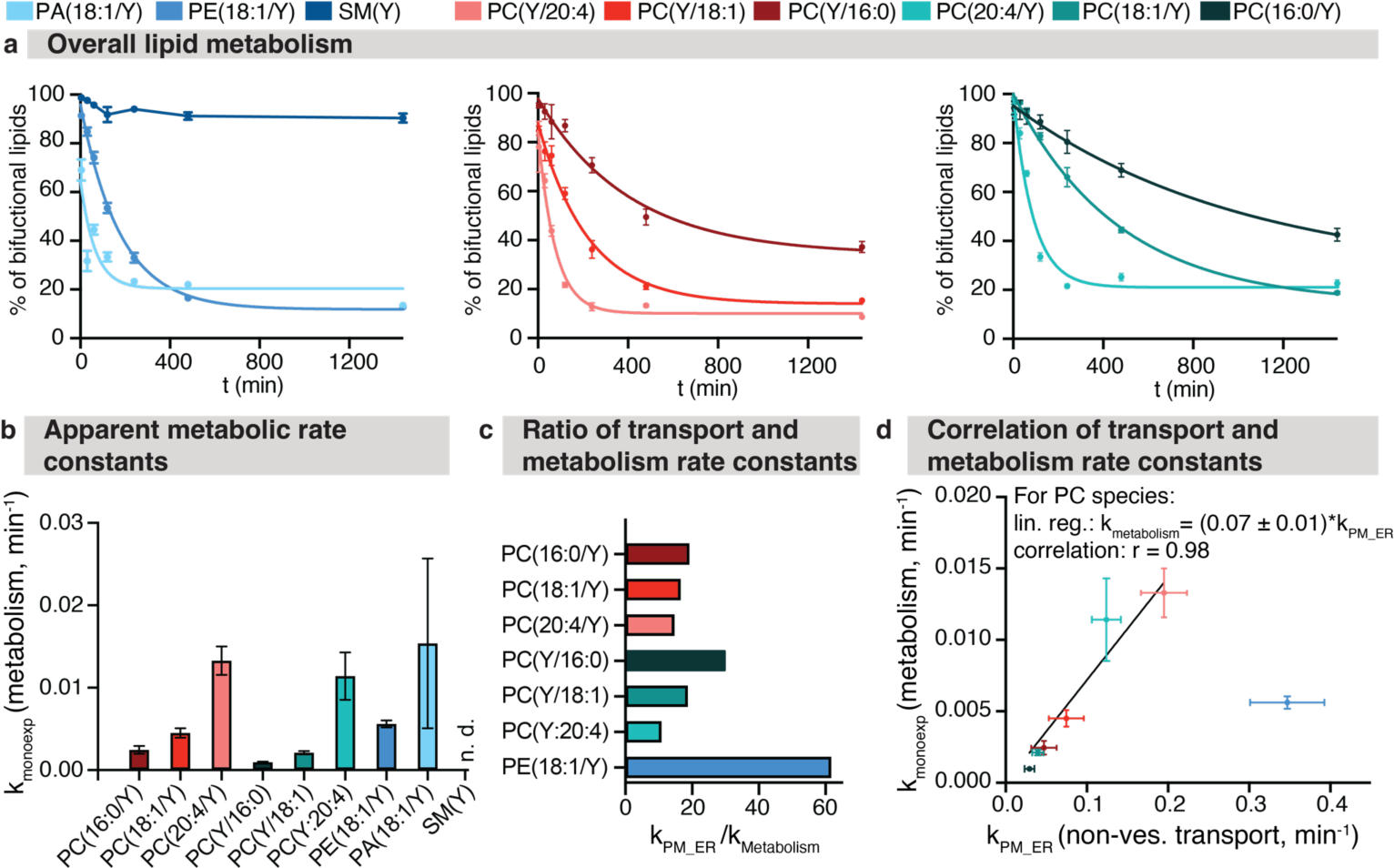
Lipid metabolism is approximately one order of magnitude slower than lipid transport. **a.** Fraction of initially supplied lipid probe as % of the bifunctional lipidome as a proxy for the speed of lipid metabolism. Solid lines indicate mono-exponential fits. SM(Y) data was not fitted as very little interconversion was observed, instead a linear interpolation is shown. Error bars: SD. **b.** Comparison of determined mono-exponential rate constants for the metabolism of individual lipid species. Error bars: SE **c.** Comparison of transport and metabolic rate constants shows that lipid transport is at least one order of magnitude faster. **d.** Lipid transport and metabolism rate constants are highly correlated for PC species despite a clear timescale separation. Error bars metabolic rate constants: SE, transport rate constants: SD, calculated from 100 MC model runs.

Polyunsaturated PC species were metabolized faster than monounsaturated and saturated PC species; PA and PE were converted faster than the corresponding PC species with the same fatty acid composition, whereas SM was largely stable. To establish the relative speed of lipid metabolism versus lipid transport, we compared the apparent rate constants of bifunctional lipid probe conversion to the non-vesicular transport rate constants from the PM to the ER (Fig. 4c). This analysis showed that metabolism is slower than transport by a factor of 10-60 for all investigated probes (Fig. 4b, c)

Interestingly, we found that transport and metabolism rate constants are highly correlated for the PC species, despite the pronounced time scale separation between transport and metabolism (Fig 4d). Since most PC lipid molecules are metabolically converted after the steady state distributions are reached, this cannot be explained by delayed access of enzymes to bulk lipids within organelle membranes. The biophysical properties of the respective lipids could be directly responsible, for instance caused by highly correlated activation energies for the transfer of lipids from the bulk membrane into binding pockets of enzymes and lipid transfer proteins, respectively. Alternatively, metabolic conversion could be directly coupled to transport, e.g. by lipid substrate handover from a lipid transfer protein to a lipid metabolizing enzyme.

Taken together, we find that lipid transport is much faster than lipid metabolism. This finding suggests that the differential steady state distribution of lipid species in the organelles of the secretory pathway mainly results from selective non-vesicular transport rather than local metabolic conversion.

### Species-specific metabolism controls accumulation of neutral lipids

On short time scales, lipid sorting was found to be dominated by non-vesicular lipid transport. To assess whether cases exist where lipid metabolism controls cellular lipid distribution, we analyzed the later time points of the time course experiments, after the steady state distributions are reached. Quantification of lipid imaging data revealed that bifunctional lipids derived from PC regioisomers bearing the same fatty acids with the bifunctional fatty acid either at the *sn-1* or *sn-2* position differently accumulated in lipid droplets (Fig. 5a,b, Extended Data 3). Supplying *sn-1*-modified PCs resulted in ≈ 2-fold higher accumulation of bifunctional lipids in lipid droplets after 24 h compared to *sn-2* modified PCs, suggesting different conversion pathways for the regioisomers (Fig. 5b). As the entire lipid droplet intensity distribution was shifted to higher intensities after supplying *sn-1*-modified PCs, this is highly likely due to differential metabolism at all lipid droplets as opposed to specialized sub-populations (Fig. 5b, Extended Data 10).

**Figure 5.**
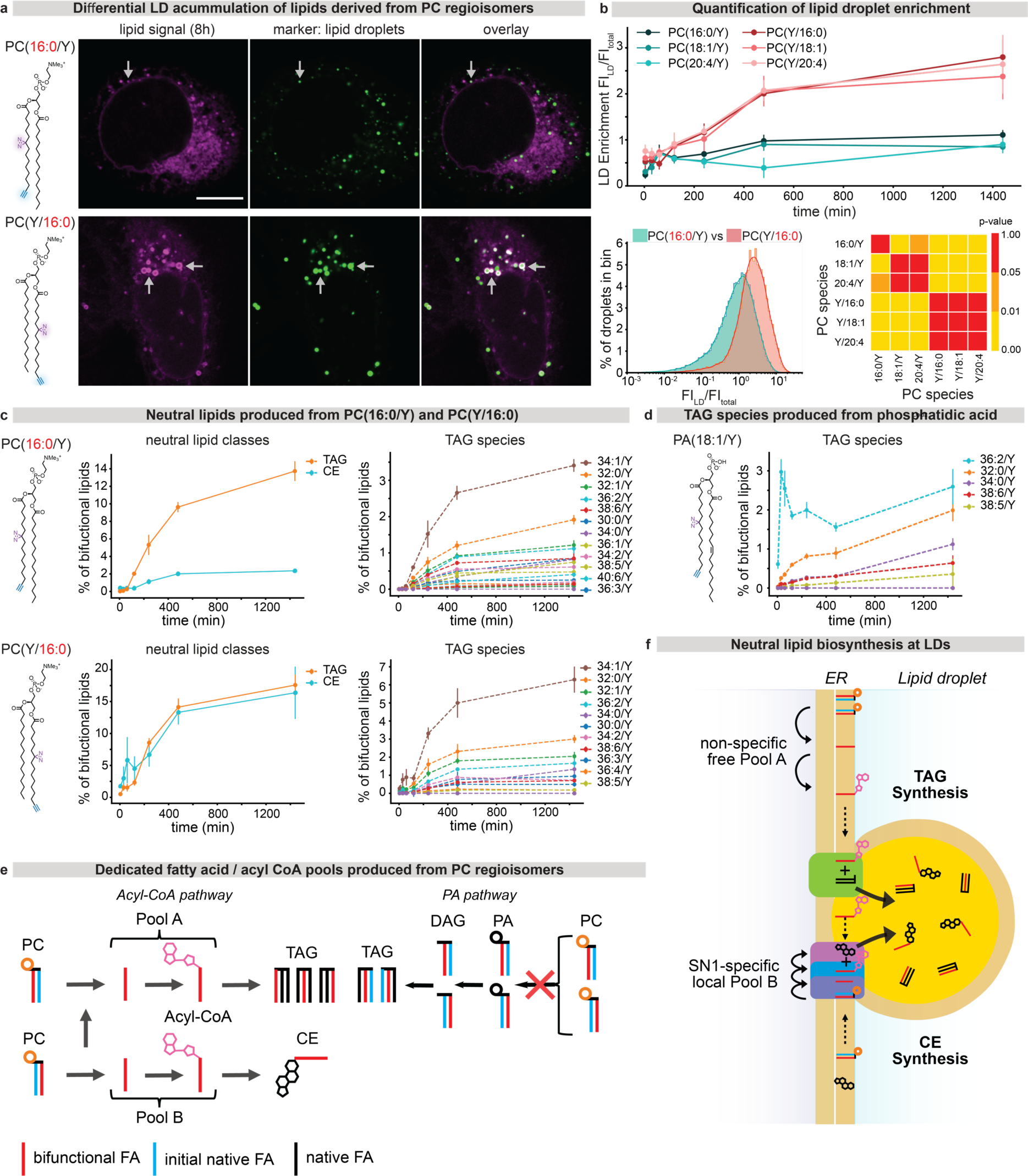
Dedicated pools of fatty acids are utilized during neutral lipid biogenesis. **a.** Lipid droplets (stained with LipidSpot 610, green) exhibit a bright lipid signal (magenta) 8h after loading PC(Y/16:0), bottom panels which is not observed after loading PC(16:0/Y), top panels. **b.** Upper panel: Quantification of fluorescence intensity of cellular lipid droplets for all PCs over time. Lower panels: Comparison of intensity distribution of individual lipid droplets for PC(16:0/Y) and PC(Y/16:0), 8 h after loading and statistical analysis of the similarity of the respective distributions for all PC species. **c.** Mass spectrometric determination of bifunctional lipid content in neutral lipids demonstrates that significantly more TAG than CE is generated after loading PC(16:0/Y) whereas similar amounts of CE and TAG are generated after loading PC(Y/16:0) (left panels). Both species yield complex TAG patterns (right panels) and all TAG species are produced with similar kinetics **d.** A single TAG species is initially produced after supplying PA (18:1/Y), whereas all other species are produced with slower kinetics **e.** Schematic overview of neutral lipid biosynthesis at lipid droplets **f.** Proposed neutral lipid biosynthetic pathway model featuring dedicated free fatty acid / Acyl-CoA pools. Error bars: 95% confidence intervals, three biological replicated containing 3 technical replicates each.

The complementary mass spectrometric data showed that the production of bifunctional cholesterol ester (CE) was up to 7-fold increased starting from the *sn-1* modified PCs compared to the respective *sn-2* regioisomers (Fig. 5c, Extended Data 10). In contrast, conversion into a wide range of triacylglycerols (TAGs) occurred with similar kinetics and abundance for all lipid probes (Fig. 5c, Extended Data 10) while native TAGs remained unchanged (Extended Data 5). Together, this suggests that the observed difference in bifunctional lipid accumulation resulted from differential CE formation rates.

The observed conversion rates into TAGs suggest a reaction sequence of an initial cleavage of the bifunctional fatty acid, generation of bifunctional Acyl-CoA and subsequent incorporation into TAGs by DGAT2 on lipid droplets (Fig. 5f). The alternative route via phosphatidic acid and DAG is incompatible with the obtained data, as supplying PA(18:1/Y) resulted in the rapid formation of a single TAG species with rapid kinetics while other species were generated much later (Fig. 5d). Since canonical CE and TAG biosynthesis routes both involve the same precursor, bifunctional Acyl-CoA, the differential PC regioisomer rates can only occur if the bifunctional fatty acid of the *sn-1* PC isomer is preferentially channeled towards CE, for example via a spatially coupled enzyme cascade comprising a *sn-1* specific phospholipase, an Acyl-CoA synthetase and a sterol O-Acyl transferase (Fig. 5e,f).

Taken together, our data indicate that cellular PC metabolism generates spatially separated pools of identical lipid metabolites for the biosynthesis of TAG and CE, respectively. These pools are preferentially accessed by specific enzymes (DGAT2 and ACAT1) and are derived from molecularly defined precursor lipid species. Thus, our data provides direct evidence for metabolic bias^40^ of specific lipid species within the lipid storage pathway and demonstrate that subcellular accumulation of neutral lipids is regulated by species-specific metabolism despite slower kinetics compared to lipid transport processes.

## Discussion

Here we introduce a pipeline to profile the transport and metabolism of individual lipid species in the organelle system of eukaryotic cells with high spatiotemporal resolution. Using this approach, we quantified the transport of 9 different lipid species through 5 organelles and their metabolic conversion over 24h and determined the underlying kinetic parameters. The complete lipid flux dataset can be interactively accessed under http://doi.org/21.11101/0000-0007-FCE5-B and the original data can be downloaded from http://doi.org/21.11101/0000-0007-FCE4-C. The kinetic comparison of transport and metabolism between different lipid species allowed us to address the fundamental question how cells maintain organelle-specific lipid compositions.

We found that small variations in lipid chemical structure strongly influenced the kinetics of non-vesicular lipid transport and metabolism, implying a high degree of selectivity on the level of individual lipid species. This is most strikingly illustrated by up to 10-fold differences in non-vesicular transport speed between individual PC species that differ in unsaturation degree and acyl chain positioning and the starkly different metabolic fate of PC regioisomers during neutral lipid biogenesis. Conversely, a similar degree of selectivity was not observed during retrograde vesicular lipid transport of lipids mediated by endocytosis. Based on our quantification, we estimate that between 85-95 % of plasma membrane lipids are transported via non-vesicular routes rather than via endocytosis in the retrograde direction (Extended Data 7), as non-vesicular lipid transport was found to be significantly faster than membrane trafficking.

Together with earlier data indicating that membrane trafficking of the bulk phospholipids PC and PE in the anterograde direction occurs mainly via non-vesicular pathways^41,42^, our results imply that organelle membrane lipid compositions are primarily maintained through a combination of fast species-specific non-vesicular lipid transport and slower, spatially controlled lipid metabolism. We found that non-vesicular lipid transport and ATP-dependent lipid flipping by P_4_ATPases are coupled, suggesting that P_4_ATPases could supply the energy required for directional transport. Furthermore, this finding provides a possible explanation for the much higher rates of lipid exchange observed between cellular membranes compared to in vitro membrane models^43^.

Our results furthermore reveal that structurally very closely related lipid species such as PC regioisomers give rise to significantly different product patterns after 24 h of metabolic turnover. Product pattern divergence is particularly pronounced during neutral lipids biosynthesis, where our results suggest a highly coupled reaction sequence of fatty acid cleavage, Acyl-CoA synthesis and incorporation into TAG/CE, respectively (Fig. 5). Being able to identify such highly localized, species-specific reaction sequences in cellular lipid processing will open the door to develop pharmaceutical interventions in lipid-related diseases with strong genetic components such as MAFLD^44^.

Taken together, our findings suggest that non-vesicular lipid transport plays a key role in the maintenance of organelle identity and reveal localized lipid metabolism on sub-organellar scales during neutral lipid biogenesis. Combining our approach with genetic interventions will shed light on the molecular mechanisms that underpin species-selective lipid transport and metabolism. We anticipate that this work will have a major impact for revealing the functions of lipids in cell biology and human diseases.

## Author contributions

KB, MS, CJL, AL and AN synthesized lipid probes. AN and AH developed the initial lipid delivery and imaging protocol. JMIA developed the final, optimized lipid imaging protocol, optimized & performed colocalization experiments with organelle markers, carried out lipid imaging time course experiments, developed the image analysis pipeline and built the website with help of the scientific computing facility. KS performed lipidomics experiments. KS, JMIA, AS and AN analyzed MS data. JMIA, AN and AH analyzed fluorescence microscopy data. RS, PR and MH performed and analyzed biophysical characterization of model membrane systems. HML, MS, KPG and PB contributed to lipid imaging protocol development. PB and HML characterized the TMEM30A HCT KD line and assessed lipid transport changes. BD performed kinetic modelling. AN prepared figures with contributions from JMIA, AH, BD, KS and PB. AN and AH wrote the manuscript with contributions from JMIA, KB and MS. JMIA, AH and AN designed the project. All authors read and commented on the manuscript.

## Acknowledgments

A.N. gratefully acknowledges financial support by the European Research Council (ERC) under the European Union’s Horizon 2020 research and innovation program (grant agreements no GA 758334 ASYMMEM and AURORA). AN, AH and AS acknowledge financial support by the Deutsche Forschungsgemeinschaft (DFG) via the TRR83 consortium. This research was supported by an Allen Distinguished Investigator Award, a Paul G. Allen Frontiers Group advised grant of the Paul G. Allen Family Foundation to AN and AH. MH and RŠ acknowledge the financial support provided by the Advanced Multiscale Materials for Key Enabling Technologies project, supported by the Ministry of Education, Youth, and Sports of the Czech Republic; project No. CZ.02.01.01/00/22_008/0004558, co-funded by the European Union. M.S. is supported by the ELISIR program of the EPFL School of Life Sciences and acknowledges funding from the Swiss National Science Foundation (SNSF grant IC00I0-227891). We thank the following services and facilities at MPI-CBG Dresden for their support: Protein Expression Facility, Mass Spectrometry Facility, Scientific Computing Facility, Genome Engineering Facility and the Light Microscopy Facility. We thank Jan Peychl, Britta Schroth-Diez, and Sebastian Bundschuh for the outstanding support and expert advice.

## Extended Data figures

**Extended Data Figure 1.**
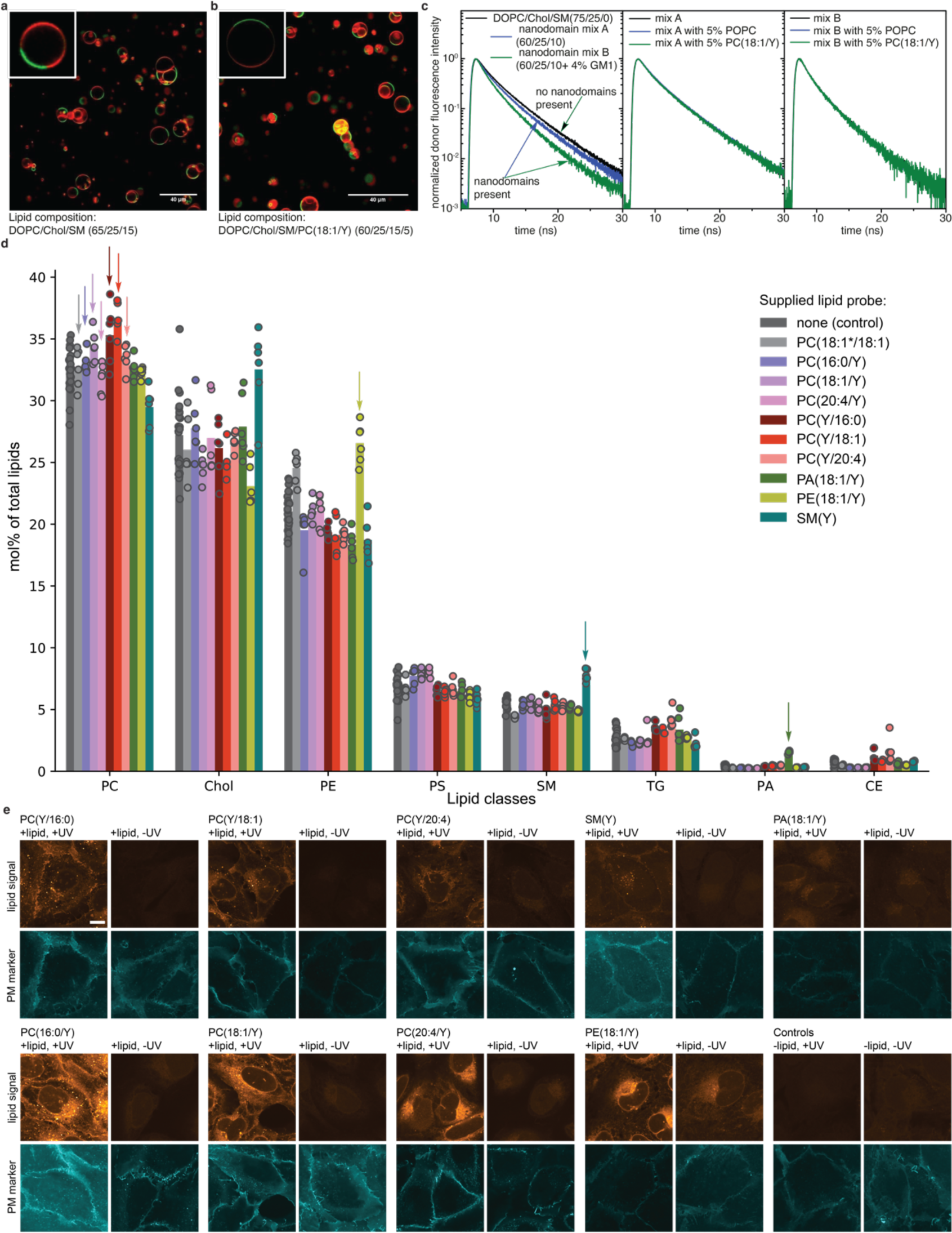
Biophysical characterization of bifunctional lipid containing model membranes, lipidome assessment after bifunctional lipid loading and lipid imaging signal comparison for all probes. **a, b.** Formation of liquid ordered L_o_ (stained by Bodipy-FL-GM_1_; green) and liquid disordered L_d_ (stained by DiD; red) microdomains is unaffected by replacing 5 % of DOPC content with PC(18:1/Y) in giant unilamellar vesicles GUVs. **c.** Formation of ganglioside nanodomains leading to faster deexcitation of Bodipy-FL-GM_1_ donors via FRET is unaffected by replacing 5 % of POPC content with PC(18:1/Y) in GUVs. **d.** Comparison of lipidome composition directly after lipid loading bifunctional lipid probes (4 min timepoint) with control lipidome. Arrows indicate supplied lipid type. **e.** +UV lipid signal vs -UV lipid signal for all probes, 30 min timepoint shown. Note: The high intensity in -UV conditions for PE(18:1/Y) is explained by the fact that PE can be chemically fixed with formaldehyde due to its primary amine group, which is not the case for the other lipids.

**Extended Data Figure 2.**
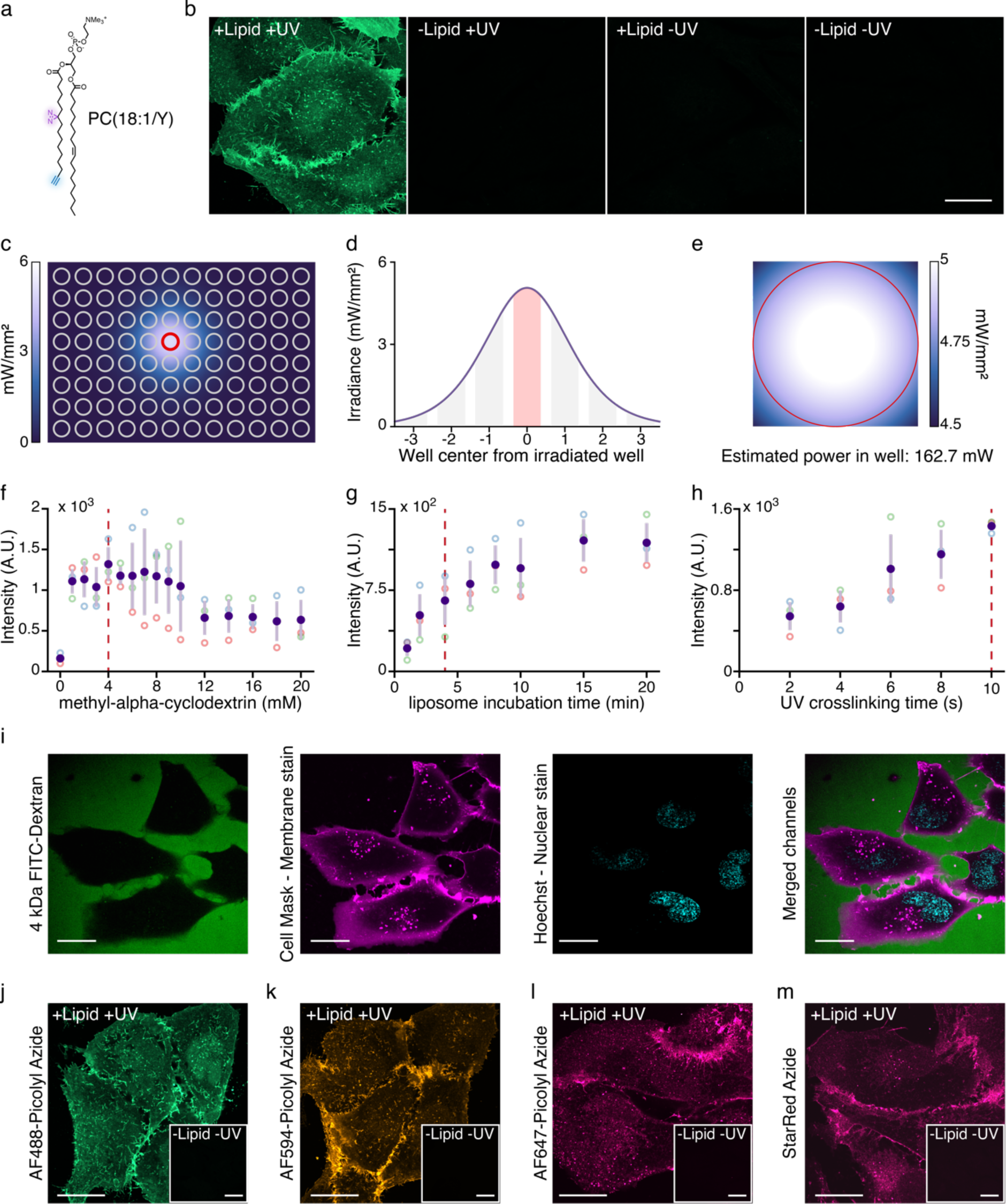
Optimization of the lipid imaging protocol. **a.** Structure of PC(18:1/Y) used for protocol optimization. **b.** Representative imaging results using optimized lipid loading, crosslinking and click chemistry conditions. **c-e.** Characterization of UV illumination in the 96-well plate format used for this study. **f-h.** Optimization of lipid loading and crosslinking conditions. Dashed red lines indicate chosen conditions. **i.** Lipid loading does not compromise cell membrane integrity as demonstrated by exclusion of 4 kDa FTIC-Dextran from cell interior. **j-m.** Lipid signal visualization using different Picolyl-Azide dyes. AF594-Picolyl-Azide was used for this study.

**Extended Data Figure 3.**
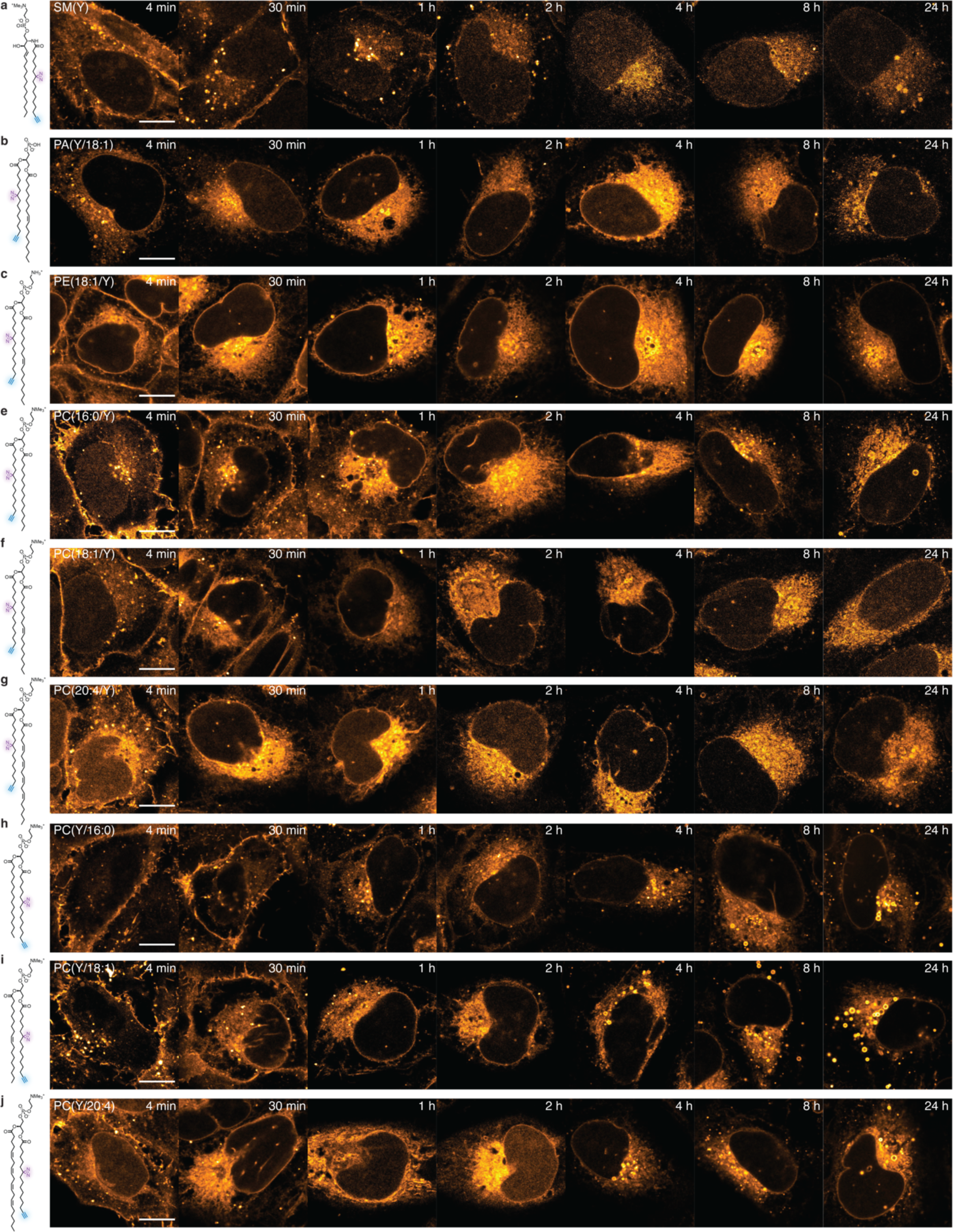
a-I Lipid transport time courses for all probes and timepoints. Representative images for lipid transport time course experiments. Scale bars: 10 μm. Images are brightness-contrast adjusted to facilitate comparing intracellular lipid localization. The full dataset 3D dataset including marker channels can be accessed on https://lipidimaging.org/.

**Extended Data Figure 4.**
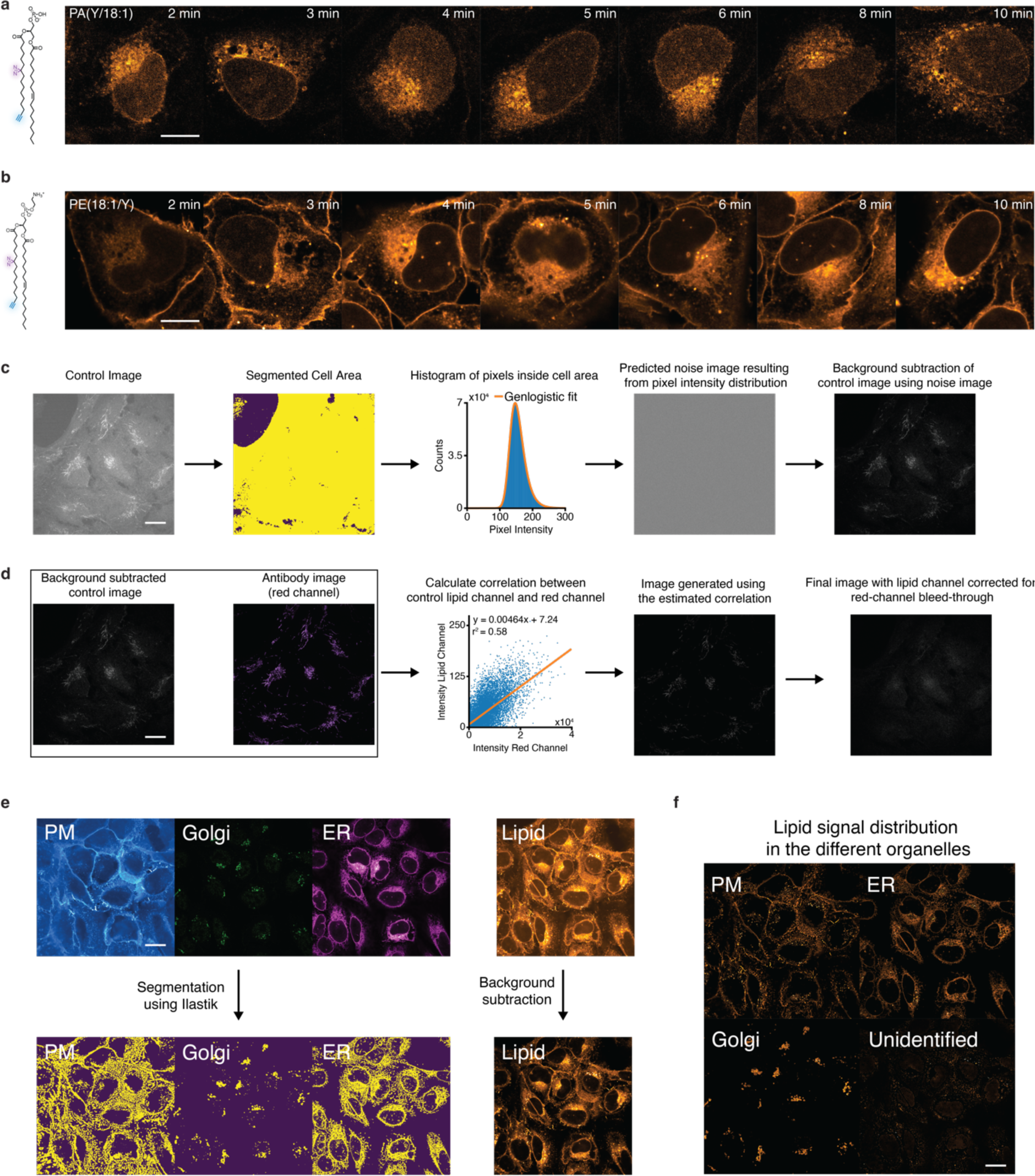
High time-resolution time courses for PE (18:1/Y) and PA(18:1/Y) and image analysis pipeline. **a, b.** Representative images for lipid transport time course experiments at higher time resolution using PA(18:1/Y) and PE(18:1/Y). Scale bars: 10 μm. Images are brightness-contrast adjusted to facilitate comparing intracellular lipid localization. **c, d.** Background subtraction strategy. For most data, background was removed using a predicted noise image derived from control images (+UV, -lipid). In cases where a AF647-Tom20 antibody was used as a mitochondrial stain, we observed a faint mitochondrial signal in the AF594 (lipid) channel in control conditions. For the corresponding +lipid images we estimated the extent of the bleedthrough signal by determining the correlation between the mitochondrial signal in the marker channel and the lipid channel, using these parameters to generate an image for the expected artefactual mitochondrial signal in +lipid images & subtracting it from the raw +lipid image. Scale bars: 20 µm. **e.** Segmentation of marker channels to generate probability masks and representative result of lipid channel background removal. Scale bar: 20 µm **f.** Lipid signal assignment to individual organelles shown in **e**. Scale bar: 20 µm

**Extended Data Figure 5.**
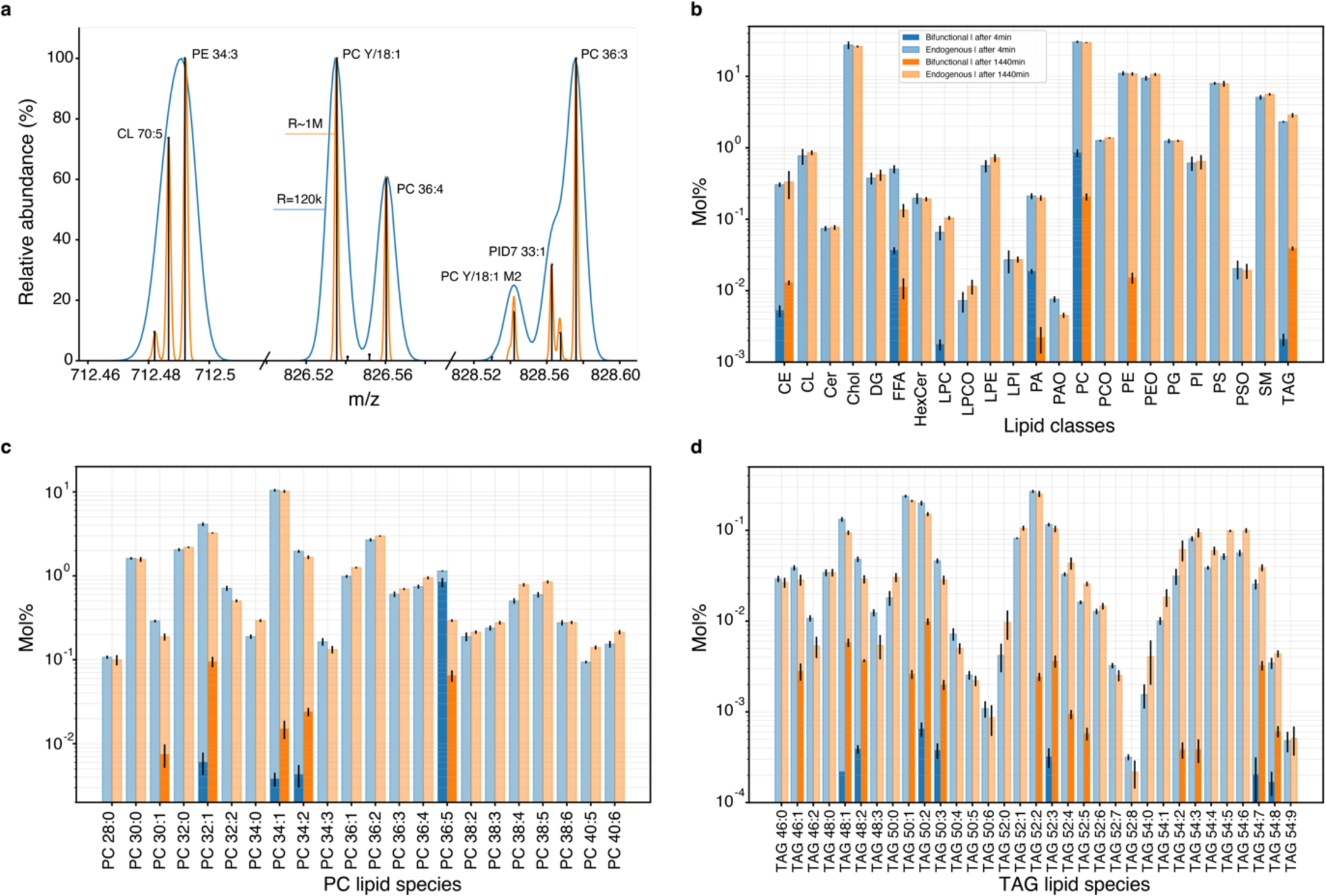
Ultra-high resolution (UHR) shotgun lipidomics of bifunctional lipid probes. **a.** Shotgun UHR mass spectrometry resolves lipid peaks spaced by a few mDA and matches bifunctional precursors and their metabolites in multiple lipid classes. Blue line: Section of the spectrum acquired at the conventional (R_s_ 120,000) resolution on Q Exactive mass spectrometer; orange line: Same spectrum section acquired at R_s_ ∼ 1M resolution using optional Booster X2 data processing system and extended (2 sec) transients. Vertical lines are peak centroids. **b.** Mol% profile acquired at two time points (see inset for color coding) of 23 lipid classes (light bars), of which 7 classes comprise lipids with bifunctional lipid moieties (dark bars) produced from PC Y/20:4. **c.** PC profile covering 22 species with 5 species containing the bifunctional fatty acid. PCs bearing a bifunctional fatty acid (16:1) are annotated as endogenous lipids having the same number of carbons and double bonds in both FA moieties, albeit having different (+28.0061 Da) masses. **d.** Bifunctional fatty acids from the source PC(Y/20:4) are, incorporated into different lipid classes e.g. the cellular TAG pool consisting of of 33 species with 14 species bearing the bifunctional fatty acid. The molar abundance of PC species containing the bifunctional fatty acid other than PC(Y/20:4) (**c**) and TAG (**d**) species increases with time, while the abundance of the starting PC(Y/20:4) decreases. Molar% profiles of native lipid classes (**b**), but also the species profile within PC and TAG classes (**c**, **d**) are not perturbed, indicating that the supplemented bifunctional lipids act as true tracer compounds and do not change the overall lipidome compositions.

**Extended Data Figure 6.**
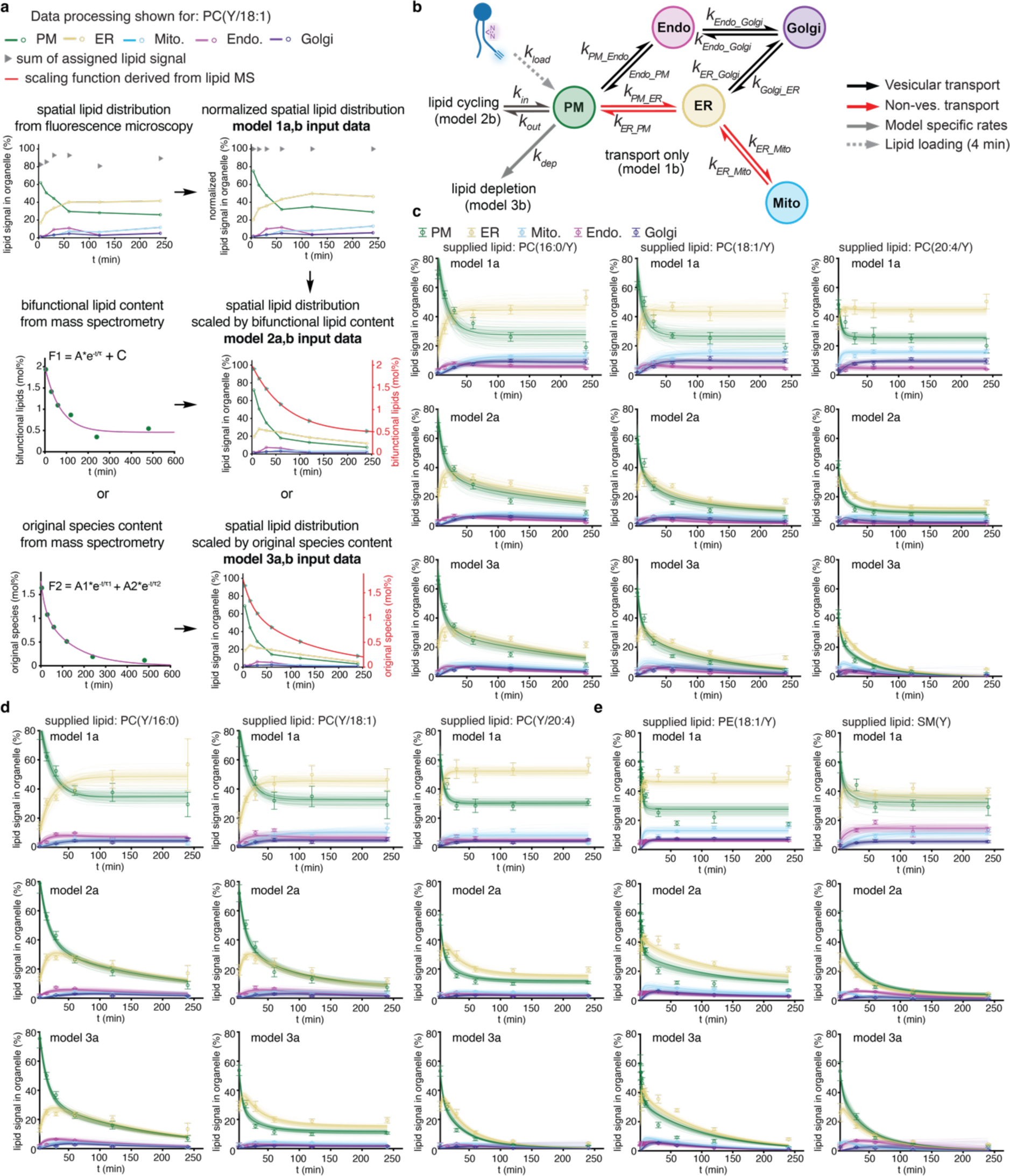
Kinetic analysis of lipid imaging and lipid MS data. **a.** Data processing steps for quantification results from lipid imaging time course experiments exemplarily shown for PC(Y/18:1) (see Supplementary Information for details). **b.** Transport scheme detailing kinetic models 4-6. **c.** Model 1-3 fits for PC(16:0/Y), PC(18:1/Y), PC(20:4/Y). **d.** Model 1-3 fits for PC(Y/16:0), PC(Y/18:1), PC(Y/20:4). **e.** Model 1-3 fits for PE(18:1/Y), SM(Y).

**Extended Data Figure 7.**
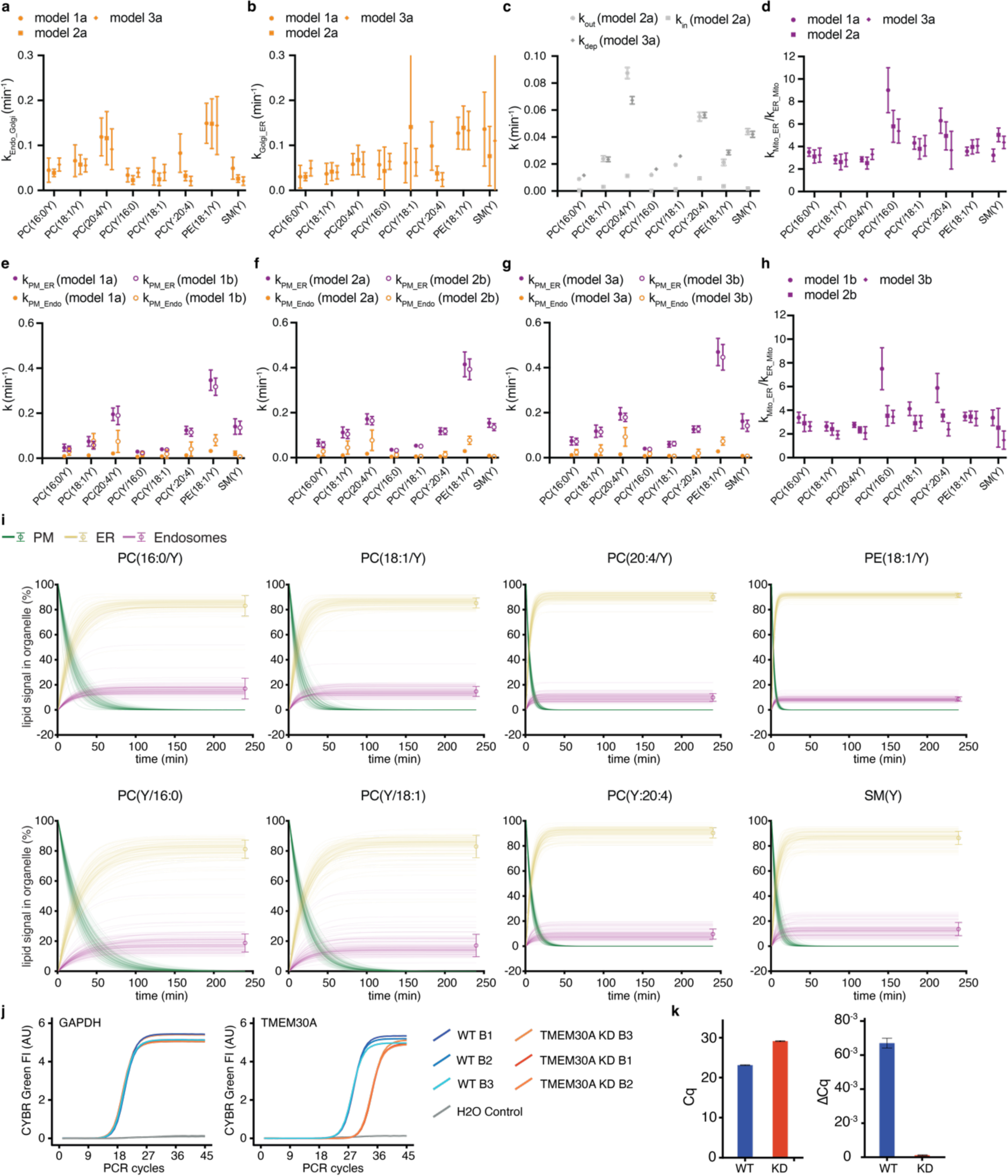
Results of kinetic analysis, estimation of lipid flow through vesicular and non-vesicular pathways and TMEM30a KD characterization. **a, b.** Rate constants for vesicular transport from endosomes to the Golgi and from the Golgi to the ER derived from models 1-3. **c.** Rate constants describing lipid cycling (lipid exchange with the extracellular space, model 2) and lipid depletion (model 3). **d.** Ratio of rate constants describing lipid exchange between the ER and mitochondria (models 1-3). Note: Individual rate constants could not be identified from the data, presumably as preceding lipid transport steps were rate-limiting. **e-g.** Comparison of rate constants describing retrograde vesicular transport from the PM to endosomes and retrograde non-vesicular transport from the PM to the ER for all analysed lipid probes, corresponding models 1a and 1b, 2a and 2b, 3a and 3b shown together. **h.** Ratio of rate constants describing lipid exchange between the ER and mitochondria (models 1b, 2b, 3b). Note: Individual rate constants could not be identified from the data, presumably as preceding lipid transport steps were rate-limiting. **i.** Fraction of bifunctional lipids transported via the non-vesicular route to the ER and the vesicular route to endosomes during retrograde transport, model 1a rate constants used for simulations. **j.** Confirmation of TMEM30A KD in HCT116 cells shown by qPCR of GAPDH and TMEM30A in WT and KD cells. **k.** Quantification cycle (Cq) and Cq normalized to GAPDH (ΔCq).

**Extended Data Figure 8.**
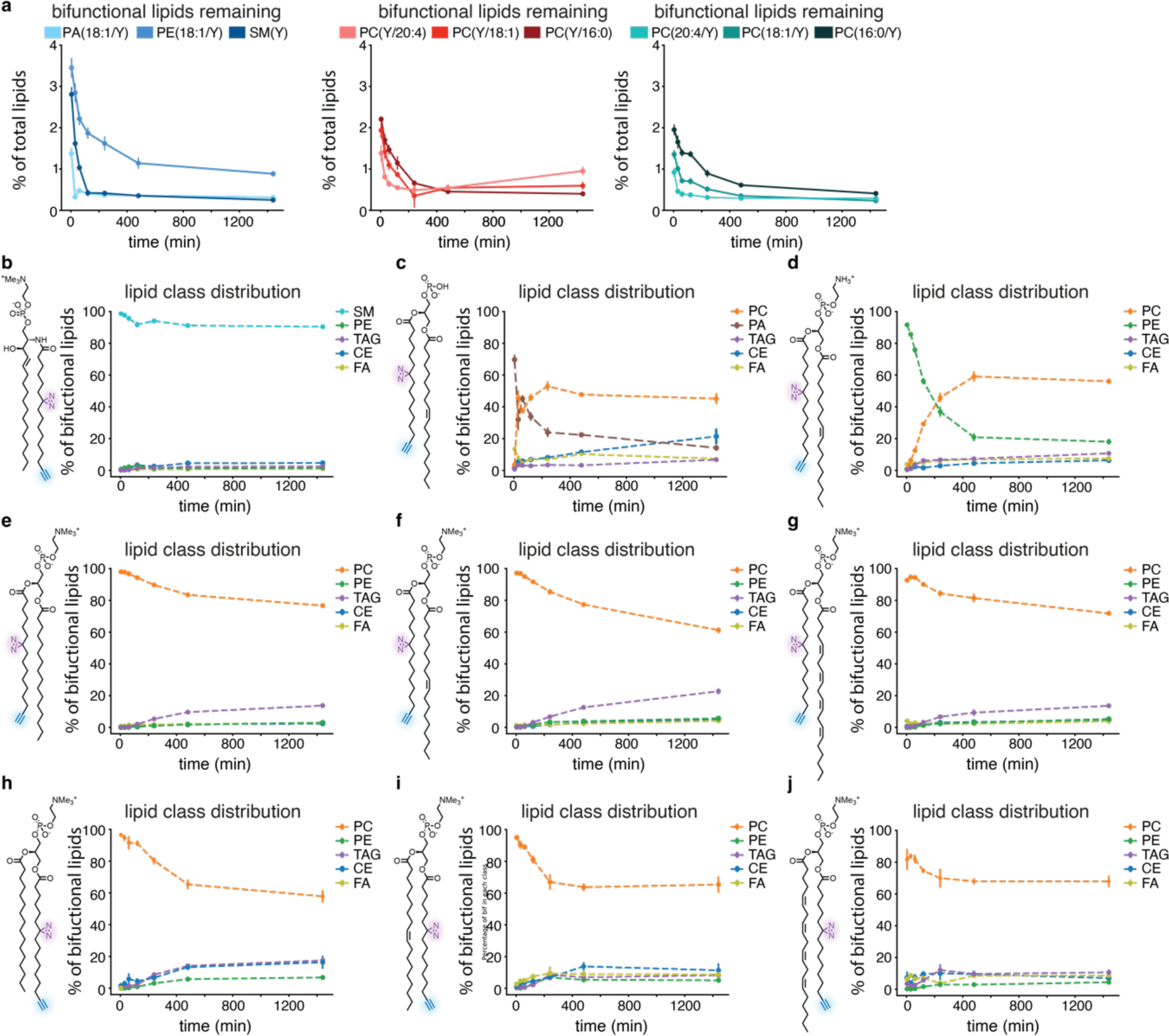
Determination of overall bifunctional lipid content and metabolism of the lipid class level. **a.** Bifunctional lipid incorporation and subsequent depletion over 24 h determined by shotgun lipidomics **b.** Development of bifunctional lipid class distribution over 24 h for all lipid probes. Note that final distributions are not identical, even for closely related species.

**Extended Data Figure 9.**
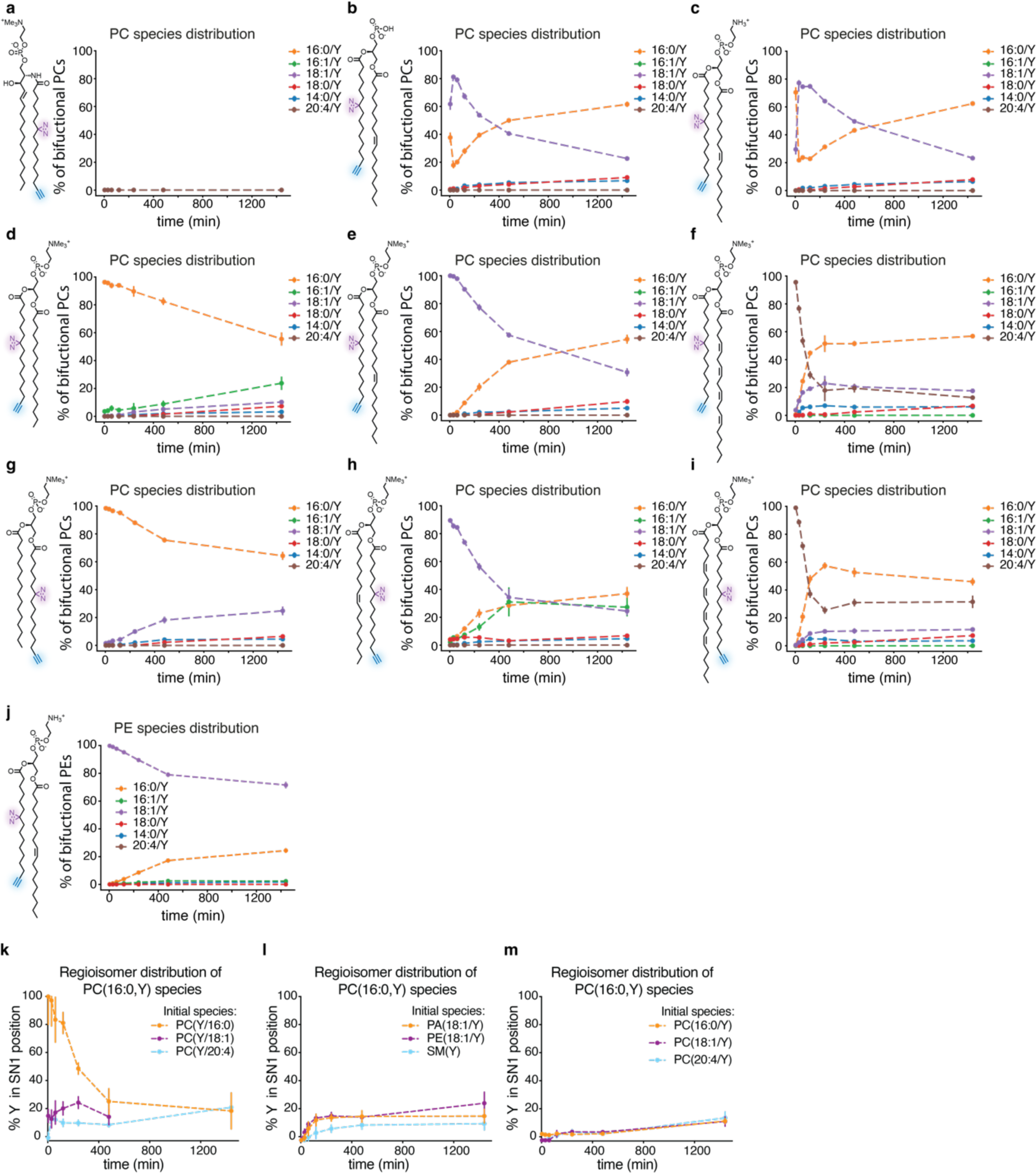
Analysis of PC species distribution. **a-i.** Development of PC species distribution over 24 h for all lipid probes. Note that some species, notably PC(16:1/Y) are only produced from a subset of the initially supplied lipids. For SM(Y), no detectable amount of PC was observed. **j.** Development of PE species distribution over 24 h after loading PE(18:1/Y). **k-m.** Development of the regioisomer distribution of the most common PC species PC(16:0,Y) estimated via the MS/MS-fatty acid neutral loss fragments. The bifunctional fatty acid is primarily incorporated at the sn-2 position.

**Extended Data Figure 10.**
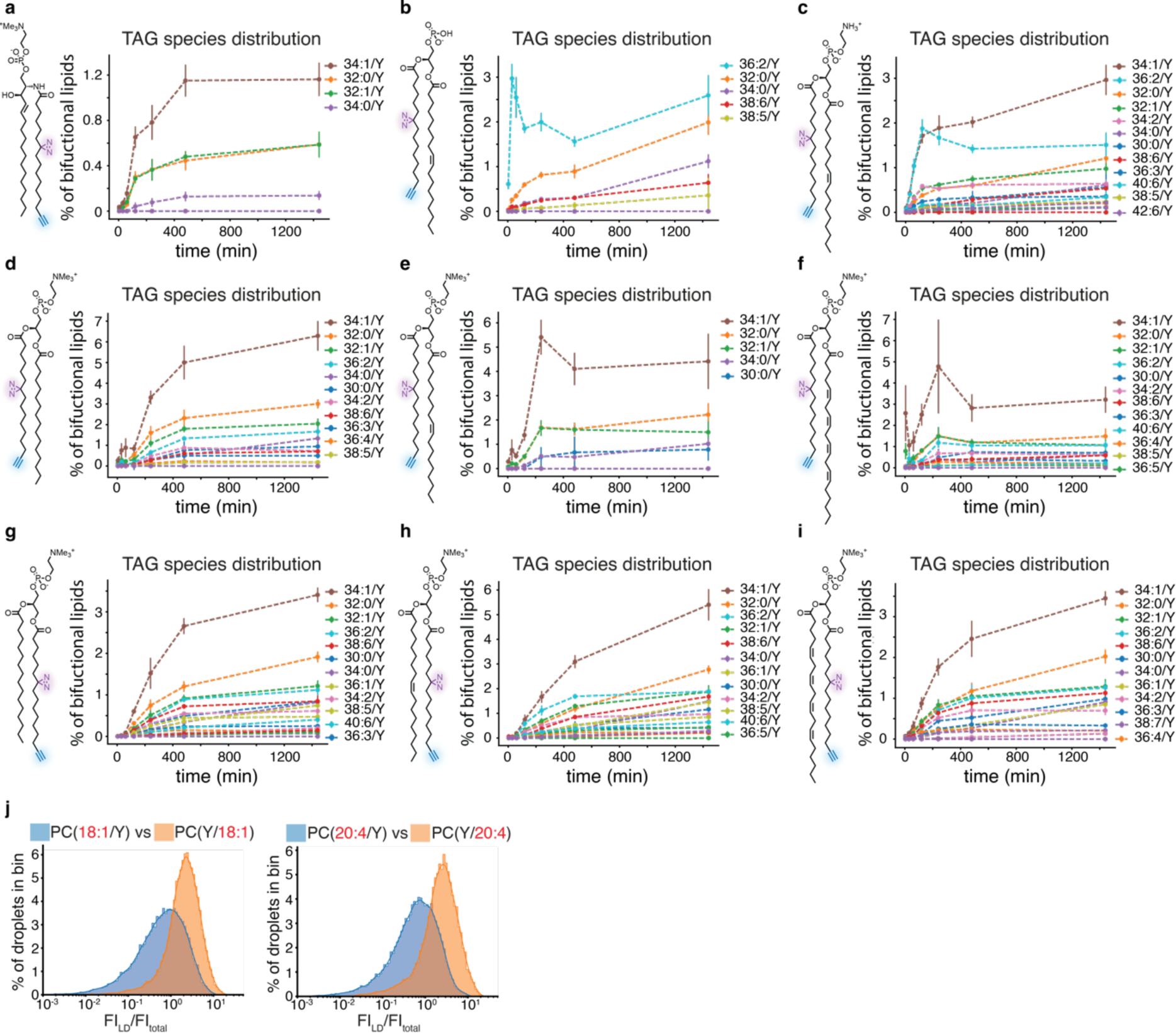
Analysis of TAG species distribution and lipid droplet populations. **a-i.** Development of TAG species distribution over 24 h for all lipid probes. Note that PA is the only lipid that initially gives rise to a single TAG species, whereas all other probes yield a spectrum of TAGs. **j.** Comparison of intensity distribution of individual lipid droplets for PC(18:1/Y) and PC(Y/18:1) and PC(20:4/Y) and PC(Y/20:4).

## References

1. Harayama, T. & Riezman, H. Understanding the diversity of membrane lipid composition. Nat Rev Mol Cell Biol 19, 281–296 (2018).

2. van Meer, G., Voelker, D. R. & Feigenson, G. W. Membrane lipids: where they are and how they behave. Nature reviews. Molecular cell biology 9, 112–124 (2008).

3. Kim, Y. & Burd, C. G. Lipid Sorting and Organelle Identity. Cold Spring Harb Perspect Biol 15, a041397 (2023).

4. 21.11101/0000-0007-FCE5-B.

5. Klose, C., Surma, M. A. & Simons, K. Organellar lipidomics — background and perspectives. Current Opinion in Cell Biology 25, 406–413 (2013).

6. Sampaio, J. L. et al. Membrane lipidome of an epithelial cell line. Proceedings of the National Academy of Sciences of the United States of America 108, 1903–7 (2011).

7. Holthuis, J. C. M. & Menon, A. K. Lipid landscapes and pipelines in membrane homeostasis. Nature 510, 48–57 (2014).

8. Reinisch, K. M. & Prinz, W. A. Mechanisms of nonvesicular lipid transport. Journal of Cell Biology 220, e202012058 (2021).

9. Koivusalo, M., Jansen, M., Somerharju, P. & Ikonen, E. Endocytic Trafficking of Sphingomyelin Depends on Its Acyl Chain Length. MBoC 18, 5113–5123 (2007).

10. Haberkant, P. & Holthuis, J. C. M. Fat & fabulous: Bifunctional lipids in the spotlight. Biochimica et Biophysica Acta - Molecular and Cell Biology of Lipids 1841, 1022–1030 (2014).

11. Höglinger, D. et al. Trifunctional lipid probes for comprehensive studies of single lipid species in living cells. Proceedings of the National Academy of Sciences of the United States of America 114, (2017).

12. Haberkant, P. et al. In vivo profiling and visualization of cellular protein-lipid interactions using bifunctional fatty acids. Angewandte Chemie - International Edition 52, 4033–4038 (2013).

13. Höglinger, D. Bi- and Trifunctional Lipids for Visualization of Sphingolipid Dynamics within the Cell. in 95–103 (2019). doi:10.1007/978-1-4939-9136-5_8.

14. Altuzar, J. et al. Lysosome-targeted multifunctional lipid probes reveal the sterol transporter NPC1 as a sphingosine interactor. Proceedings of the National Academy of Sciences 120, e2213886120 (2023).

15. Farley, S., Stein, F., Haberkant, P., Tafesse, F. G. & Schultz, C. Trifunctional Sphinganine: A New Tool to Dissect Sphingolipid Function. ACS Chem. Biol. 19, 336–347 (2024).

16. Schuhmacher, M. et al. Live-cell lipid biochemistry reveals a role of diacylglycerol side-chain composition for cellular lipid dynamics and protein affinities. Proc. Natl. Acad. Sci. U.S.A. 117, 7729–7738 (2020).

17. Höglinger, D., Nadler, A. & Schultz, C. Caged lipids as tools for investigating cellular signaling. Biochimica et Biophysica Acta - Molecular and Cell Biology of Lipids 1841, 1085–1096 (2014).

18. Jiménez-López, C. & Nadler, A. Caged lipid probes for controlling lipid levels on subcellular scales. Current Opinion in Chemical Biology 72, 102234 (2023).

19. Frank, J. A. et al. Photoswitchable diacylglycerols enable optical control of protein kinase C. Nature Chemical Biology 12, 755 (2016).

20. Morstein, J., Impastato, A. C. & Trauner, D. Photoswitchable Lipids. ChemBioChem 22, 73–83 (2021).

21. Haldar, S. & Chattopadhyay, A. Application of NBD-Labeled Lipids in Membrane and Cell Biology. in Fluorescent Methods to Study Biological Membranes (eds. Mély, Y. & Duportail, G.) 37–50 (Springer, Berlin, Heidelberg, 2013). doi:10.1007/4243_2012_43.

22. Klymchenko, A. S. & Kreder, R. Fluorescent probes for lipid rafts: from model membranes to living cells. Chemistry & biology 21, 97–113 (2014).

23. Triebl, A. & Wenk, M. R. Analytical Considerations of Stable Isotope Labelling in Lipidomics. Biomolecules 8, 151 (2018).

24. Postle, A. D. & Hunt, A. N. Dynamic lipidomics with stable isotope labelling. Journal of Chromatography B 877, 2716–2721 (2009).

25. Thiele, C. et al. Tracing fatty acid metabolism by click chemistry. ACS Chemical Biology 7, 2004–2011 (2012).

26. Thiele, C., Wunderling, K. & Leyendecker, P. Multiplexed and single cell tracing of lipid metabolism. Nat Methods 16, 1123–1130 (2019).

27. Wunderling, K., Zurkovic, J., Zink, F., Kuerschner, L. & Thiele, C. Triglyceride cycling enables modification of stored fatty acids. Nat Metab 5, 699–709 (2023).

28. Koukalová, A. et al. Lipid Driven Nanodomains in Giant Lipid Vesicles are Fluid and Disordered. Sci Rep 7, 5460 (2017).

29. Sarmento, M. J. et al. The impact of the glycan headgroup on the nanoscopic segregation of gangliosides. Biophysical Journal 120, 5530–5543 (2021).

30. Berg, S. et al. ilastik: interactive machine learning for (bio)image analysis. Nat Methods 16, 1226–1232 (2019).

31. Chang, C.-L. & Liou, J. Phosphatidylinositol 4,5-Bisphosphate Homeostasis Regulated by Nir2 and Nir3 Proteins at Endoplasmic Reticulum-Plasma Membrane Junctions. J Biol Chem 290, 14289–14301 (2015).

32. Lees, J. A. & Reinisch, K. M. Inter-organelle lipid transfer: a channel model for Vps13 and chorein-N motif proteins. Current Opinion in Cell Biology 65, 66–71 (2020).

33. Hanna, M., Guillén-Samander, A. & Camilli, P. D. RBG Motif Bridge-Like Lipid Transport Proteins: Structure, Functions, and Open Questions. Annual Review of Cell and Developmental Biology 39, 409–434 (2023).

34. Guillén-Samander, A. et al. A partnership between the lipid scramblase XK and the lipid transfer protein VPS13A at the plasma membrane. Proceedings of the National Academy of Sciences 119, e2205425119 (2022).

35. Matoba, K. et al. Atg9 is a lipid scramblase that mediates autophagosomal membrane expansion. Nat Struct Mol Biol 27, 1185–1193 (2020).

36. Li, Y. E. et al. TMEM41B and VMP1 are scramblases and regulate the distribution of cholesterol and phosphatidylserine. Journal of Cell Biology 220, e202103105 (2021).

37. Lorent, J. H. et al. Plasma membranes are asymmetric in lipid unsaturation, packing and protein shape. Nat Chem Biol 16, 644–652 (2020).

38. van der Velden, L. M. et al. Heteromeric Interactions Required for Abundance and Subcellular Localization of Human CDC50 Proteins and Class 1 P4-ATPases*. Journal of Biological Chemistry 285, 40088–40096 (2010).

39. Bryde, S. et al. CDC50 proteins are critical components of the human class-1 P 4-ATPase transport machinery. Journal of Biological Chemistry 285, 40562–40572 (2010).

40. Harayama, T. Metabolic bias: Lipid structures as determinants of their metabolic fates. Biochimie 215, 34–41 (2023).

41. Vance, J. E., Aasman, E. J. & Szarka, R. Brefeldin A does not inhibit the movement of phosphatidylethanolamine from its sites for synthesis to the cell surface. Journal of Biological Chemistry 266, 8241–8247 (1991).

42. Kaplan, M. R. & Simoni, R. D. Intracellular transport of phosphatidylcholine to the plasma membrane. Journal of Cell Biology 101, 441–445 (1985).

43. Wong, L. H., Čopič, A. & Levine, T. P. Advances on the Transfer of Lipids by Lipid Transfer Proteins. Trends in Biochemical Sciences 42, 516–530 (2017).

44. Mahmoudi, S. K. et al. Exploring the role of genetic variations in NAFLD: implications for disease pathogenesis and precision medicine approaches. Eur J Med Res 29, 190 (2024).

