## Supplementary Information for "Quantitative imaging of species-specific lipid transport in mammalian cells"

[2] École polytechnique fédérale de Lausanne, 1015 Lausanne, Switzerland

[3]J. Heyrovský Institute of Physical Chemistry, Academy of Sciences of the Czech Republic v.v.i., Dolejškova 3, 18223 Prague, Czech Republic

[4]Helmholtz Zentrum Dresden Rossendorf, Institute of Resource Ecology, Bautzner Landstraße 400, 01328 Dresden, Germany

[5]Technische Universität Dresden, Biotechnologisches Zentrum, Center for Molecular and Cellular Bioengineering (CMCB), Dresden, Germany.

[6]Cluster of Excellence Physics of Life, TU Dresden, Dresden, Germany.

These authors contributed equally: Kai Schuhmann and Kristin Böhlig

### Lipid Metabolism Annex

This section covers observations from mass spectrometric data regarding lipid flux that could not be included in the main text due to space reasons, but may be of significant interest for readers interested in general lipid metabolism. The following discussion is mostly limited to the glycerophospholipid (GPL) probes, as SM(Y) was found to be metabolically stable.

#### Nomenclature

Product species are written XX(ZZ/Y) for simplicity, as fragment ion data for the most common PC species indicate that the bifunctional fatty acid is primarily incorporated in the sn2 position (ED Figure 9, k-m). This analysis is not feasible for most other lipid species as the underlying mass spectrometric signal intensities are not high enough to conduct this type of analysis lipidome-wide. For lipid species other than PC(16:0/Y), this does not represent an annotation of acyl chain attachment site.

#### General characterization of bifunctional lipid content and composition

We used two measures to characterize the overall temporal development of bifunctional lipid content and composition over time: The fraction of initially supplied lipid probe as a fraction of the total abundance of lipids carrying the bifunctional fatty acid and the fraction of bifunctional lipids in relation to the total lipid abundance. The first measure serves as a proxy for lipid metabolism, whereas the second parameter captures all processes that lead to loss of bifunctional lipids, including shedding of membrane and dilution by ongoing cell growth. For the glycerophospholipids, the obtained kinetics are reasonably similar for both measures, pointing to a major role of lipid metabolism in depleting the bifunctional lipid pool. For SM(Y), this is not the case, pointing to a yet-unidentified alternative mechanism for the observed, relatively quick drop in overall bifunctional lipid content (Supplementary Figure 1)

Between 10 and 40 % of the bifunctional lipidome was represented by the originally supplied lipid species for GPLs (approximately 90% for SM) throughout the entire chase period of 24 h, which we did not initially expect as polyunsaturated PC species or phosphatidic acid are low-abundance lipids reported to have short half-life times in cellular membranes[1,2]. The continued presence of short-lived lipids suggests the formation of lipid pools less susceptible to metabolism.

There are at least two possible explanations for such lipid pools. The most straightforward would be an extracellular reservoir which continuously supplies the original species into the metabolic network at a slow rate. This could e.g. be due to a membrane shedding/re-uptake dynamic. Early in the chase period, extracellular vesicles[3] could be formed or membrane lost during cell migration and membrane remodelling events[4] while the original species is still overrepresented in the plasma membrane or multivesicular bodies[5]. This material would then be slowly incorporated again, as cells grow and divide, resulting in the observed quasi-steady state. Alternatively, a percentage of the initially supplied lipids could become structural lipids in lipid-protein complexes or other lipid-protein assemblies and feature much slower overall conversion.

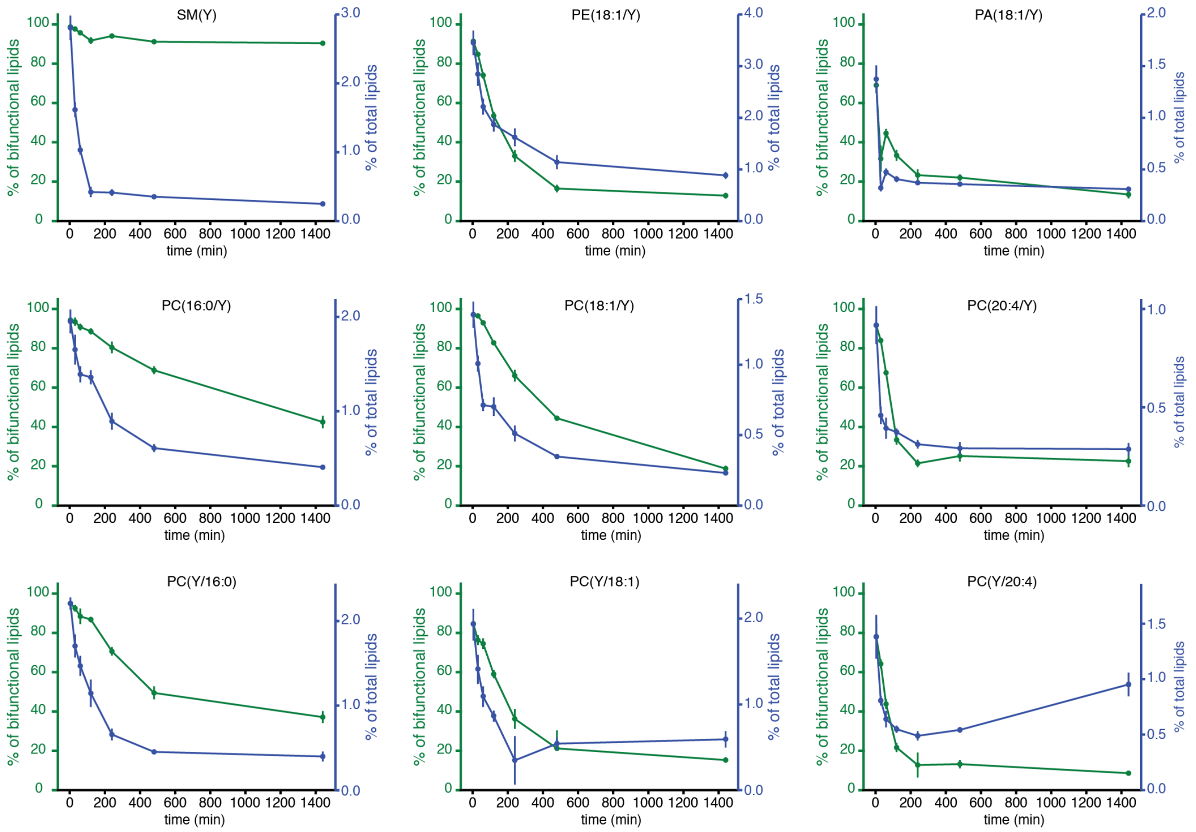

**Supplementary Figure 1 General measures of bifunctional lipid content and metabolism. Green: Fraction of originally supplied species of total bifunctional lipid abundance. Blue: Fraction of total bifunctional lipids of total lipid abundance.**

### PC metabolism

Overall, we found that between 50 and 80% of bifunctional lipids after 24 h were PC species for all initially supplied bifunctional PCs (Supplementary figures 2-7, 9, 10), with the rest mostly comprised of neutral lipids and free fatty acid. We observed only very limited conversion into other glycerophospholipids. Newly generated PC species were mostly saturated and monounsaturated species of varying chain length. Arachidonate-containing, polyunsaturated species were only detected at higher levels when the initially supplied species was also bearing an arachidonic acid side chain. Since arachidonate containing PC species are rapidly metabolized, this is most straightforwardly explained by an extracellular lipid reservoir as discussed above. Thus, the persistent presence of arachidonate containing PC in the cells supplied with this species could result from re-absorption from the medium.

Among the most commonly detected PC species, two appeared to be genuine examples of highly selective metabolic routes. PC(16:1/Y) is not generated at all after loading 4 out of 6 supplied PC species, but constitutes a major fraction (20-30% of all PCs) of the eventual product pattern for PC(Y/18:1) and PC(16:0/Y). We furthermore found that up to 8-10% of the short-chain variant PC(14:0/Y) was generated with relatively fast kinetics (min-h) from arachidonate-containing PCs, but not from more saturated PC species. This could be a spillover-effect as the available pool of endogenous 16:0- and 18:1-Lyso-PCs is depleted by incorporation of the bifunctional fatty due to quick conversion of arachidonate-containing PCs. Alternatively, the selective generation of short-chain species could be initiated to counteract the rise in polyunsaturated, long-chain species, which would require acyltransferases with a matching selectivity profile.

In contrast to the selectively generated species PC(16:1/Y) and PC(14:0/Y), PC(16:0/Y) was the most common PC species at 30-60% of total bifunctional PC abundance after 24 h for all supplied glycerophospholipid probes and PC(18:1/Y) and PC(18:0/Y) were typically present between 10 and 30% of total bifunctional PC abundance, indicating that these species were generated by non-selective bulk routes.

Taken together, we find that acyl chain remodelling and neutral lipid generation are the primary pathways observed after phosphatidylcholine loading, with examples of species-specific conversions in both routes.

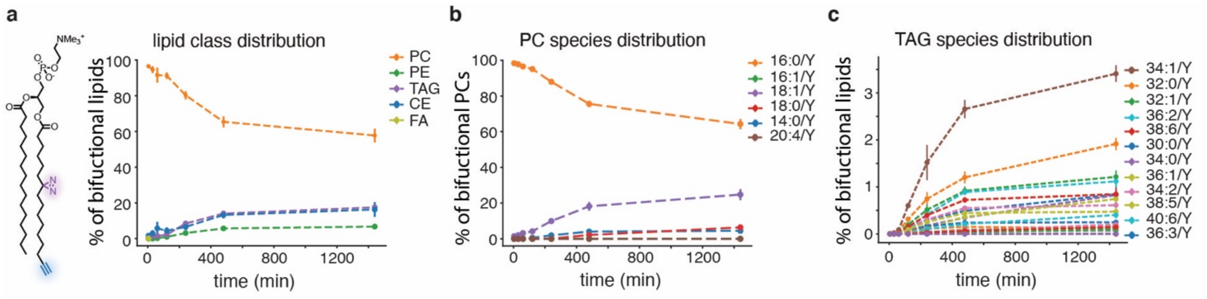

**Supplementary Figure 2 Metabolism of PC(Y/16:0).**

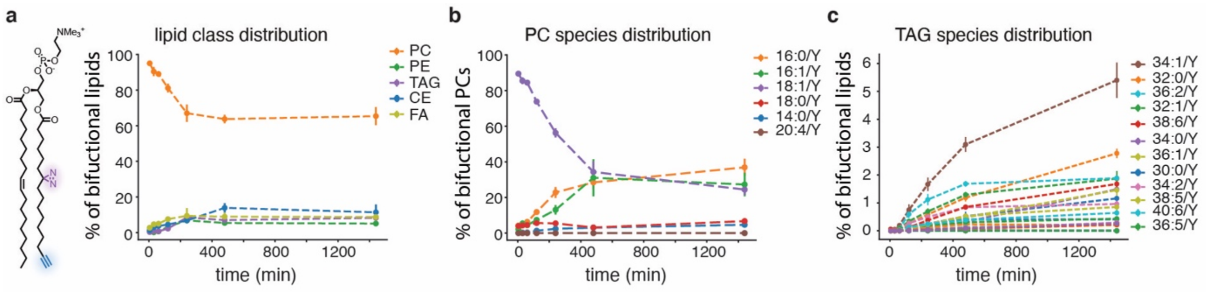

**Supplementary Figure 3 Metabolism of PC(Y/18:1).**

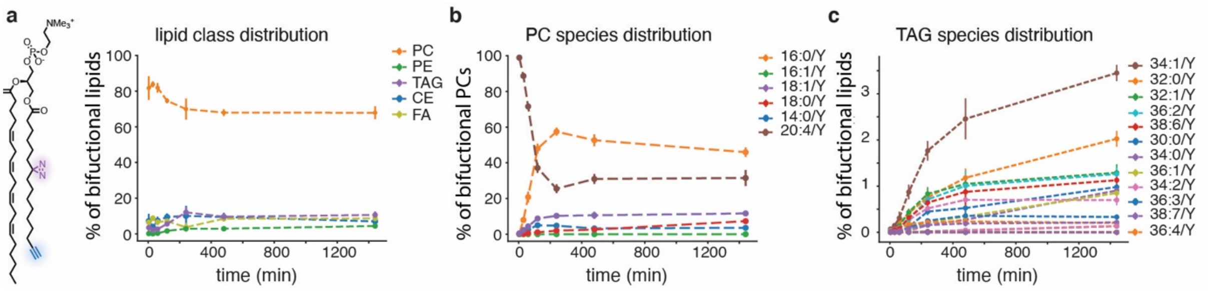

**Supplementary Figure 4 Metabolism of PC(Y/20:4).**

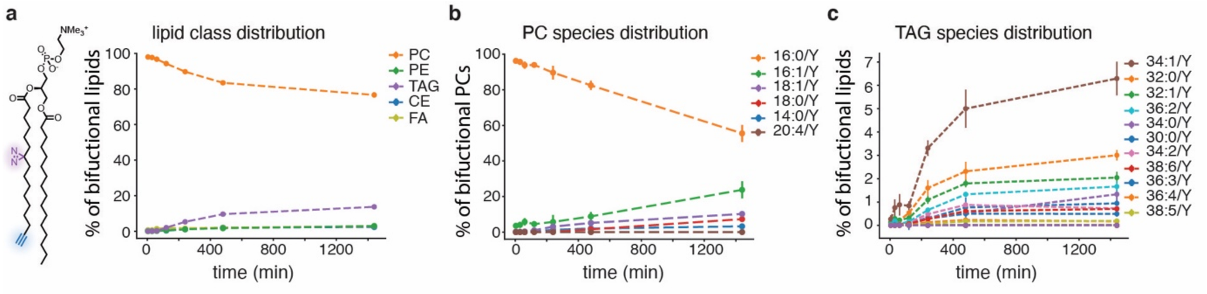

**Supplementary Figure 5 Metabolism of PC(16:0/Y).**

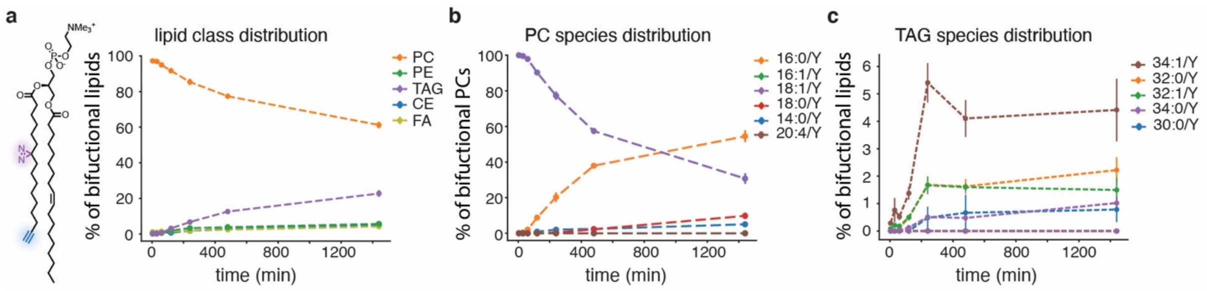

**Supplementary Figure 6 Metabolism of PC(18:1/Y).**

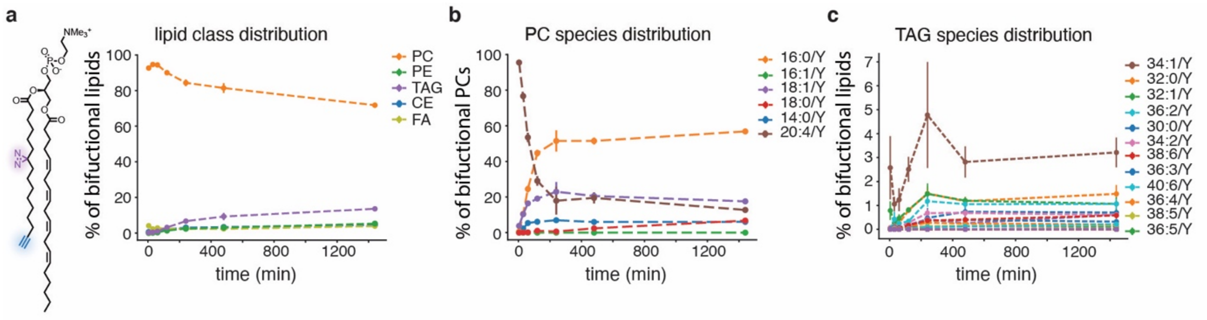

**Supplementary Figure 7 Metabolism of PC(20:4/Y).**

### PA and PE metabolism

Both PA(18:1/Y) and PE(18:1/Y) were overall metabolized with significantly faster kinetics than the corresponding PC(18:1/Y) species (Supplementary Information Fig. 1). However, the original PA and PE species were still present at approximately 20% of total bifunctional lipid abundance after 24 h, likely due to membrane shedding/re-uptake as discussed above. The main product of both PA(18:1/Y) and PE(18:1/Y) was PC. The main biosynthetic route apparently changed over time, as the fatty acid composition of the generated PC species featured complex but almost superimposable kinetics (Supplementary Information Fig. 8d, 9d). Initially, a significant amount of a PC species is generated that deviates in its fatty acid composition from the supplied lipid probe (PC(16:0/Y) vs either PE/PA(18:1/Y)), in line with cleavage of the bifunctional fatty acid and incorporation into lysophosphatidylcholine. In the following early portion of the time-course, PC(18:1/Y) becomes the predominant product, which points to headgroup modification pathways. This order is reversed again, presumably as acyl chain remodelling of the PC species generated in the interim becomes the dominant metabolic process. For both PA(18:1/Y) and PE(18:1/Y), neutral lipids are the next common metabolic products, in the case of PA(18:1/Y), more CE than TAG is generated, while the opposite is true for PE(18:1/Y).

In contrast to all supplied PC species, almost no acyl chain remodelling was observed for PA(18:Y) and only a very limited amount for PE(18:1/Y). In fact, as PE(16:0/Y) was the only additional PE species detected in meaningful amounts through the course of the experiment, this likely does not represent bona fide PE acyl chain remodelling but PE biogenesis starting from the primarily generated PC species PC(16:0/Y).

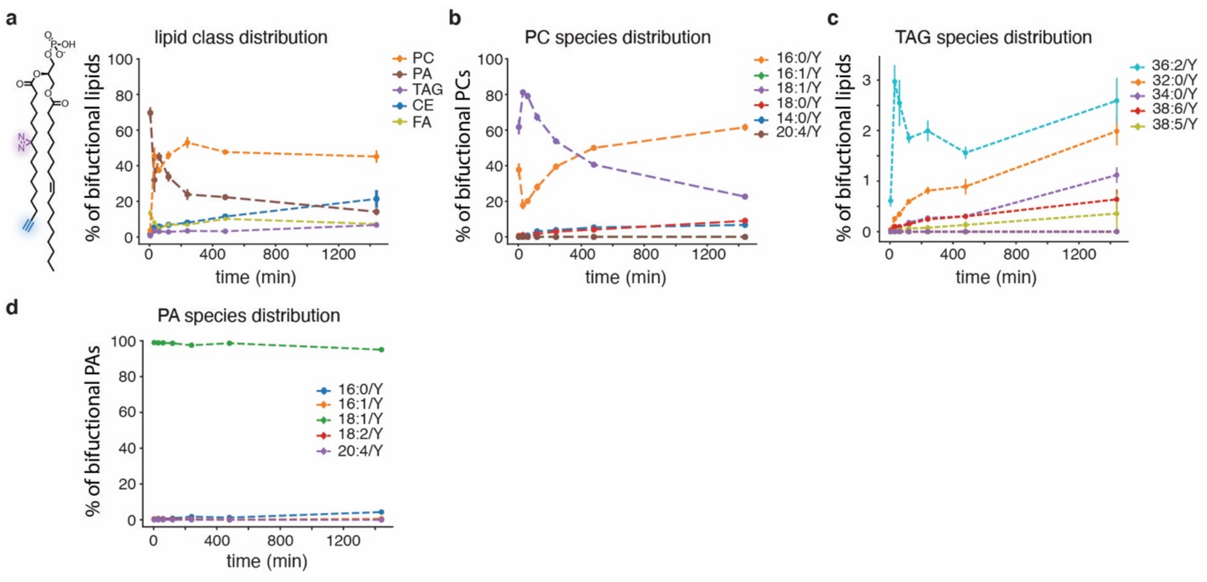

**Supplementary Figure 8 Metabolism of PA(18:1/Y).**

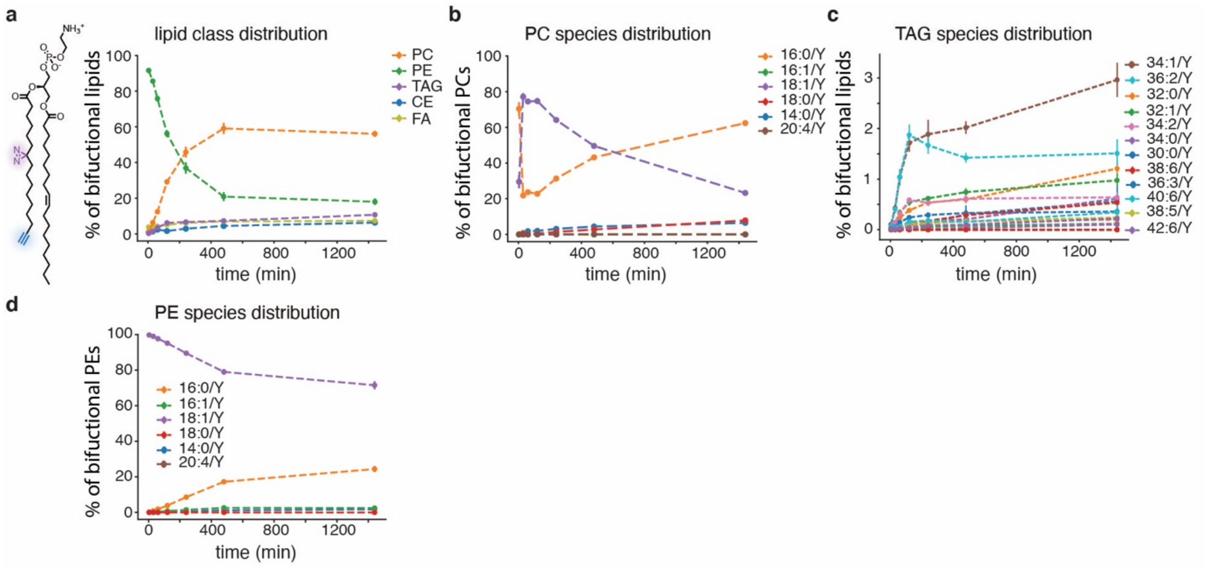

**Supplementary Figure 9 Metabolism of PE(18:1/Y).**

### SM Metabolism

SM(Y) was found to me metabolically rather stable, indicating that the cleavage of the fatty acid was the rate limiting step in this process (Supplementary Information Fig. 10).

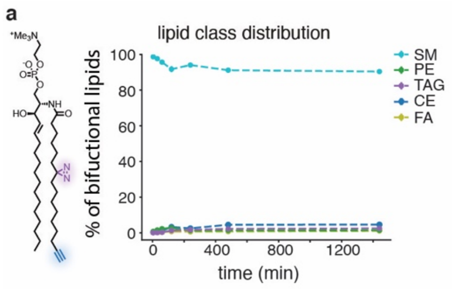

**Supplementary Figure 10 Metabolism of SM(Y).**

### Materials

#### Liposome preparation

1-palmitoyl-2-oleoyl-glycero-3-phosphocholine (POPC) (MW: 760.08) was purchased from Avanti. 100 nm polycarbonate membranes were purchased from Whatman. Cholesterol (MW: 386.65) was purchased from Sigma. LiposoFast Basic was purchased from Avestin. Chlorofom was purchased from TCI (P0018).

#### Preparation of giant unilamelar vesicles (GUVs)

1,2-dioleoyl-sn-glycero-3-phosphocholine (DOPC) (MW: 814.62), cholesterol (ovine wool) (MW: 386.65), N-stearoyl-D-erythro-sphingosylphosphorylcholine (SM) (MW: 730.59), GM1 ganglioside (bovine brain sodium salt) (MW: 1546.8), 1-palmitoyl-2-oleoyl-glycero-3-phosphocholine (POPC) (MW: 760.08) and 1,2- dipalmitoyl-sn-glycero-3-phosphoethanolamine-N-(cap biotinyl) (biotinyl-PE) (MW: 1052.66) were purchased from Avanti Polar Lipids (Alabaster, AL, USA). Fluorescent probe DiIC18(5)-DS (DiD) was purchased from Invitrogen (Carlsbad, CA, USA), while BODIPY-FL-C5-GM1 and BODIPY-564/570-C5-GM1 were synthesized by Dr. Ilya Mikhalyov and Dr. Natalia Gretskaya[6]. All used lipids were dissolved in chloroform, except of labelled and unlabelled gangliosides, which were dissolved in a mixture of chloroform and methanol in a 2:1 ratio (v/v). Organic solvents of spectroscopic grade were purchased from Merck (Darmstadt, Germany). Biotin labelled bovine serum albumin (biotinyl-BSA) and other chemicals (HEPES, D-glucose, sucrose, NaCl) were purchased from Sigma Aldrich (St. Louis, USA). Streptavidin was purchased from Iba Lifesciences (Gottingen, Germany).

#### Lipid loading onto cells

U2-OS and HCT-116 cells were purchased from ATCC. Cellview microplates, 96 well (Item No. 655891) were purchased from Greiner. Methyl-alpha-cyclodextrin (Av. MW: 1127) was purchased from Cyclodextrin-Shop. McCoy's 5A medium (16600082), FBS (A5670701) and Penicillin-Streptomycin (15140122) were purchased from Thermo Fisher Scientific. The FBS was heat inactivated by heating up the serum at 56ºC for 30 min. Hank's Balance Salt Solution (HBSS) was purchased as a salt from Merck (H6136) and prepared according to the manufacturer's instructions.

#### Cell fixation, click labelling, immunofluorescence and plasma membrane labelling

Triton X-100 (Av. MW: 625) was purchased from Serva. AF488-Picolyl-Azide (MW: 736.69), AF594-Picolyl-Azide (MW: 925.00) and AF647-Picolyl-Azide (MW: 1077.27) were purchased from Jena Bioscience. Glycine (MW: 75.07, 50046)), $CuSO_4$ (MW: 159.61, 45167), L-ascorbic acid (MW: 176.12, A92902) were purchased from Merck. MAXpack Immunostaining Media Kit was purchase from Active Motif. EZ-Link NHS-PEG4-Biotin (MW: 588.67, A39259) and Streptavidin, Alexa Fluor 405 conjugate (Av. MW: 57 056, S32351) were purchased from Thermo Fisher Scientific. The NHS-PEG4-Biotin was resuspended using acetonitrile and aliquoted, aliquots were then dried under a vacuum centrifuge and stored at -20 ºC.

##### Primary antibodies

Antibody mixes containing several antibodies were used to homogenously label organelles. In the table below, all the antibodies labelled with the same antibody mix number were combined in one solution using MAXbind Staining Solution at the specified dilution. Lipid droplets were stained with LipidSpot 610. In this study, 4 different antibody mixes were used.

1. Endosomes and Mitochondria
2. Golgi and ER
3. Lysosomes
4. Mitochondria and ER

| Antibody Mix | Antibody | Dilution | Supplier | Code |
|---|---|---|---|---|
| **1** | Rab5 (rabbit) | 1:35000 | Cell Signaling Technology | CST-3547S |
| **1** | Rab7 (rabbit) | 1:500 | Cell Signaling Technology | CST-9367S |
| **1** | Tom20 (mouse) | 1:1000 | Santa Cruz | sc-17764 |
| **1** | ATP5A (mouse) | 1:1000 | Abcam | ab14748 |
| **2** | Golgin97 (rabbit) | 1:3500 | Abcam | ab22683 |
| **2** | Giantin (rabbit) | 1:3500 | Abcam | ab80864 |
| **2** | GM130 (rabbit) | 1:3500 | Cell Signaling Technology | CST-12480S |
| **3** | Lamp1 | 1:1000 | Cell Signaling Technology | CST-9091S |
| **4** | Tom20 (rabbit) | 1:1000 | Cell Signaling Technology | CST-42406S |
| **4** | ATP5A (rabbit) | 1:1000 | Abcam | ab176569 |
| **2, 4** | Calnexin 1 (mouse) | 1:1000 | Abcam | ab112995 |
| **2, 4** | Lamin A/C (mouse) | 1:1000 | Cell Signaling Technology | CST-4777S |
| **2, 4** | BAP31 (mouse) | 1:1000 | Enzo Life Sciences | ALX-804-601-C100 |
| **2, 4** | Calreticulin (mouse) | 1:1000 | Abcam | ab84340 |

##### Secondary antibodies

| Antibody | Dilution | Supplier | Code |
|---|---|---|---|
| Goat anti-Mouse IgG AF Plus 647 | 1:1000 | Thermo Fisher Scientific | A32728 |
| Goat anti-Rabbit IgG AF-488 | 1:1000 | Thermo Fisher Scientific | A-11008 |

#### Mass spectrometry

Purified bifunctional lipids were used as standards for the original species in mass spectrometry. The following lipid standards were purchased from Avanti polar lipids and used for the metabolic products. Used solvents and other chemical were ordered from Sigma Aldrich and of LCMS quality.

| Standard | Analyte Class |
|---|---|
| CED 18:1:0 | CE |
| Cer 30:1:2 | Ceramide |
| Cer 30:1:2 | CerC16 |

| | |
|---|---|
| CholD 27:1:1 | Chol |
| CL 56:4:0 | CL |
| DGD 33:1:0 | DG |
| FFAC 18:0:0 | FFA |
| FFAC 18:0:0 | FFAC16 |
| HexCer 30:1:1 | HexCer |
| LPCD 18:1:0 | LPC |
| LPCD 18:1:0 | LPCC16 |
| LPCD 18:1:0 | LPCO |
| LPED 18:0:0 | LPE |
| LPI 13:0:0 | LPI |
| PAD 33:1:0 | PA |
| PAD 33:1:0 | PAC16 |
| PAD 33:1:0 | PAO |
| PCD 33:1:0 | PC |
| PCD 33:1:0 | PCC16 |
| PCD 33:1:0 | PCO |
| PED 33:1:0 | PE |
| PED 33:1:0 | PEO |
| PGD 33:1:0 | PG |
| PID 33:1:0 | PI |
| PSD 33:1:0 | PS |
| PSD 33:1:0 | PSO |
| SMD 36:2:1 | SM |
| SMD 36:2:1 | SMC16 |
| TGD 48:1:0 | TG |
| TGD 48:1:0 | **TGC16** |

### TMEM30 Knock-out

##### TMEM30A KO generation

Cas9 protein, guideRNAs and tracr-scaffoldRNA were obtained from Integrated DNA technologies. Neon Transfection System 100 µL Kit (MPK10096) was purchased from Thermo Scientific.

Sequences for the guideRNAs:

5'-Guide: 5'-GTTAGCATCGACTTTTCACAGGG-3'

3'-Guide 5'-GAGATTTCACGTAACGACGATGG-3'

##### TMEM30A qPCR validation

Fast Start Essential SYBR Master Mix was purchased from Roche. Primers were purchased from IDT with the following sequences:

| Name | Sequence (5' to 3') |
|---|---|
| TMEM30A Forward | CGGATGTGACACCTTGCTTTT |
| TMEM30A Reverse | ACGTAACGACGATGGTTTTGATAG |
| GAPDH Forward | GCAAATTCCATGGCACCGT |
| GAPDH Reverse | TCGCCCCACTTGATTTTGG |

### Methods

#### Liposome preparation

Liposomes containing bifunctional lipids of phosphatidylcholine (PC), phosphatidylethanolamine (PE), phosphatidic acid (PA) or sphingomyelin (SM) were prepared using a 1:2:1 ratio of POPC : bifunctional lipid : cholesterol. To this end, the different lipids were dissolved in chloroform, mixed together and dried under argon. The lipid film that formed after drying was then rehydrated with phosphate buffer saline (137 mM NaCl, 2.7 mM KCl, 10 mM $Na_2HPO_4$, 1.8 mM $KH_2PO_4$) to a final total lipid concentration of 3 mM (corresponding to 0.75 mM POPC, 1.5 mM bifunctional lipid and 0.75 mM cholesterol). After hydration with PBS, the liposomes were vortexed and sonicated repeatedly until no visible lipid aggregates remained. The solution was subsequently extruded 21 times through a 100 nm polycarbonate membrane using a LiposoFast extruder. The size of the liposomes was then measured by dynamic light scattering (DLS) in a Zetasizer Nano ZSP (Malvern Panalytical). Typical liposomes had a hydrodynamic radius of ~120 nm and a polydispersion index of 0.1.

#### Preparation of GUVs

Giant unilamellar vesicles (GUVs) were obtained by electroformation as previously described[7]. Briefly, lipid films were prepared from chloroform lipid stock solutions deposited on two titanium plates. To ensure complete solvent evaporation, the plates were placed on a heated surface at about 35°C and then subjected to vacuum for at least 1 hour. The lipid-coated plates were then assembled and the resulting chamber was filled with sucrose buffer (103 mOsm/kg). The electro-swelling was performed at 47 °C at an alternating 10 Hz electrical field starting from 0.02 V to 1.1 V (peak to peak) for the initial 45 min, followed by 2 Hz at 1.1 V for 1.5h. The generated GUVs were transferred to BSA-biotin/streptavidin pre-coated imaging chambers (8-well µ-Slide Ibidi, Munich, Germany) filled with a glucose buffer (~80 mM glucose, 10 mM HEPES and 10 mM NaCl, pH 7.2, 103 mOsm/kg).

#### Fluorescence lifetime imaging microscopy (FLIM) measurements in GUVs

Fluorescence lifetime imaging microscopy (FLIM) measurements were conducted using a custom-built setup comprising an inverted confocal microscope body (IX71, Olympus, Hamburg, Germany) and a pulsed diode laser (LDH-P-C-470, 470 nm, PicoQuant, Berlin, Germany) operating at a 10 MHz repetition rate. The laser intensity was maintained at 2 µW to prevent pile-up effects, and it was coupled to a polarization-maintaining single-mode optical fiber before being re-collimated at the output using an air space objective (UPLSAPO 4x, Olympus). The light was then directed onto a water immersion objective (UPLSAPO 60x, Olympus) via a 470/635 dichroic mirror. Donor emission signals were isolated using a 515/50 bandpass filter (Chroma Rockingham, VT) and detected using a single photon avalanche diode.

#### Characterization of microphase-separated GUVs

To characterize the impact of PC(18:1/Y) on the formation of lipid microdomains (**Extended Data Figure 1**) we employed classical fluorescence imaging (without FLIM). Specifically, microscopically phase-separated GUVs congaing DOPC/Chol/SM (60/25/15) or DOPC/Chol/SM/PC(18:1/Y) (55/25/15/5) were prepared. Whereas the liquid-disordered phase (Ld) phase of GUVs was stained by DiD (red), the liquid-ordered (Lo) phase was stained by Bodipy-FL-C5-GM1 (green). The analysis was based on a qualitative comparison of fluorescence images recorded for the GUVs with or without PC(18:1/Y).

#### Characterization of ganglioside nanodomains by FRET

The impact of PC(18:1/Y) on lipid nanodomains in GUVs was determined using the following lipid mixtures: DOPC/Chol/SM (75/25/0) as a control, and nanodomain mix A (65/25/10) or nanodomain mix B (65/25/10+4% GM1) both with 5% POPC or with 5% PC(18:1/Y). Time-resolved fluorescence (TRF) decays were analysed using BODIPY-FL-C5-GM1 and BODIPY-564/570-C5-GM1 (at the probe to lipid ratio 1:200) as donor acceptor pair. Each GUV was imaged at its cross-section with a resolution of 512x512 pixels (0.6 ms/pixel), and an experimental donor fluorescence decay was recorded. A minimum of 5 vesicles were analysed per composition, and all measurements were conducted at a temperature of 25 °C.

The analysis is outlined in detail in a previous publication[8]. It is based on the use of fluorescently labeled gangliosides, specifically Bodipy-FL-GM1 as donors and Bodipy-564/570-GM1 as acceptors, forming a FRET pair. These molecules tend to accumulate within ganglioside nanodomains, thereby bringing donors and acceptors into close proximity and enhancing FRET efficiency. The presence of nanodomains significantly influences the fluorescence deexcitation kinetics of the donors when they are in close vicinity to the acceptors. Previous studies have demonstrated that the shape of the donor decay curve carries valuable information about the average size and total area covered by the nanodomains. Thus, by solely comparing the shapes of the decay curves (Extended Data 1), insights into whether there have been alterations in nanodomain characteristics can be obtained.

#### Lipid loading onto cells

U-2 OS or HCT116 cells were seeded in a 96-well plate one day before treatment at an initial density of 7-10 x $10^3$ cells/well or 40 x $10^3$ cells/well, respectively. Cells were cultured in 200 µL of McCoy's 5A medium supplemented with 10% FBS and 5 U/mL of penicillin streptomycin.

Liposome stocks (3 mM of total lipid) were diluted to final concentration of 0.5 mM in a solution of Hank's Balance Salt Solution (HBSS) containing 4.8 mM methyl-alpha-cyclodextrin on the same day the lipid imaging experiment was carried out. The resulting solution containing 0.5 mM total lipid and 4 mM methyl-alpha-cyclodextrin was incubated for 30 minutes at 37 ºC before addition of the liposomes to the cells.

Before treating the cells with the liposome/cyclodextrin loading solution, cells were washed 2 times with pre-warmed HBSS medium. After washing, 80 µL of the loading solution were added per well and incubated for 4 minutes with the cells at 37 ºC, unless otherwise stated. For high-time-resolution experiments (PE and PA), the incubation period was reduced to 30 seconds. For the 4 min time point NHS-PEG$_4$-Biotin was added to the liposome/cyclodextrin solution to a final concentration of 5 mM immediately before loading. For PE (18:1/Y) this was not done to avoid acylation of the primary amine of the PE headgroup. In this case, biotin treatment was shorten to 30 seconds after the 30 seconds lipid incubation. The cells were then washed 2 times with pre-warmed McCoy's 5A medium.

For all timepoints after the very first one, the washing medium was exchanged with complete McCoy's 5A medium supplemented with 10% FBS and 5 U/mL of penicillin-streptomycin for the appropriate time period. 5 mM NHS-PEG4-Biotin solution in HBSS was added to the cells for 4 minute .4 minutes before UV irradiation and fixation to obtain a plasma membrane stain. The solution was removed and cells washed with pre-warmed HBSS prior to UV irradiation and fixation.

#### UV crosslinking of bifunctional probes

Bifunctional lipids within the cells were crosslinked by illuminating the corresponding wells for 10 s (unless otherwise stated) with approximately 0.1 W/cm$^2$ with a 305 nm wavelength 2x2 mm LED from beneath the 96-well plate through the bottom glass plate. The LED, supplied by Laser Componenets GmbH, had a maximum power output of 300 mW and the cells were placed at an approximately distance of 0.5 mm from the light source (Compare ED2).

#### Cell fixation, click labelling, immunofluorescence and plasma membrane labelling

After UV crosslinking, the HBSS was immediately removed and replaced with 4% formaldehyde in PBS for cell fixation. Complete removal of HBSS prior to addition of the fixative was found to be crucial for maintaining cellular architecture on the organelle level and prevented membrane blebbing. Cells were fixed for 20 min at room temperature and then washed three times with 0.1% Triton X-100 and 100 mM glycine in PBS for permeabilization. Cells were left in the last washing solution for 1 hour at room temperature to remove non-crosslinked bifunctional lipids and prevent thermal crosslinking of the remaining lipids. The cells were then washed three times with PBS to remove excess detergent. Cells could now be stored at 4ºC prior to immunostaining and click labelling.

Immunostaining was performed using the MAXpack Immunostaining Media Kit. After permeabilization, the cells were incubated with the MAXblock Blocking Medium for 1 hour at 37 ºC. This solution was then washed with MAXwash Washing Medium. The washing medium was removed and the desired primary antibody mix was added to the cells in MAXbind Staining Medium. The antibodies were incubated overnight at 4ºC. Cells were then washed three times with MAXwash Washing Medium, incubating the cells for 10 minutes at RT during each washing step. Afterwards, the secondary antibodies were added to the cells in MAXbind Staining Medium, and samples were incubated with the secondary antibody staining solution for 1 hour at RT. Cells were then washed as before with MAXwash Washing Medium.

The click reaction was performed after immunostaining in 100 mM Hepes pH 7.25. The-reaction-mix consisted of: 2 mM azide dye, 5 mM L-ascorbic acid, 0.5 mM Tris((1-hydroxy-propyl-1H-1,2,3-triazol-4-yl)methyl)amine (THPTA), 0.1 mM $Cu_2SO_4$. The cells were covered with 80 µL of the click-reaction-mix and incubated for 40 min at 37 ºC. After the click reaction the cells were washed three times with MAXwash Washing Medium.

Finally, the MAXwash Wahing Medium was removed and cells were covered with 40 µl of SlowFade diamond containing 0.1 µM of Steptavidin, Alexa Fluor 405 conjugate to label the plasma membrane. Stained samples could be were stored at 4ºC for up to 2 weeks prior to imaging, without discernible loss of fluorescence intensity.

For labelling lipid droplets, 0.2 µl of LipidSpot were added to the well after the other channels were imaged. LipidSpot was incubated for 10 min before imaging.

### Fluorescence imaging

Images were acquired using a 100X silicone immersion oil objective (UPLSAPO100XS) from Olympus mounted in an Olympus IX83 microscope. The microscope was fitted with a Yokogawa CSU-W1 SoRa unit and images were captured by an ORCA-Flash 4.0 V3 digital CMOS camera or an ORCA-Fusion from Hamamatsu. Four laser lines were used for excitation: 405, 488, 561 and 647 nm with their respective filter sets. All images for the lipid transport dataset were acquired using identical settings. The plasma membrane was always imaged in the 405 channel and the lipid in the 561 channel. The 488 and 647 channels were used for two other organelles. Cells were imaged using 20-slice z-stacks, with 0.3 µm distance in z between frames. To ensure that all images were acquired at the same starting plane, the Olympus TruFocus Z-drift compensation system was used. Each frame was exposed for 250 ms with 75% laser power.

### Image analysis

##### General information and nomenclature

Fluorescence microscopy z-stack images were analyzed in a High Performance Computing cluster using Python[9] and Ilastik[10]. The Python code and the Ilastik models used can be found in http://doi.org/21.11101/0000-0007-FCE3-D. All image stacks consisted of a 20-frame stack and 4 color channels. Images were sorted in two folders, according to the experimental conditions: one for control ("Ctrl") and another one for the images of cells containing the lipid ("Lipid"). Images can be browsed in http://doi.org/21.11101/0000-0007-FCE5-B, original images can be downloaded from http://doi.org/21.11101/0000-0007-FCE4-C.

Control ("Ctrl"):

Control images were samples not loaded with bifunctional lipids but otherwise treated the same to assess nonspecific background signal. Both +UV and -UV conditions were recorded. Control images were acquired on every 96-well plate.

Lipid images ("lipid"):

"Lipid" images were samples treated with the respective bifunctional lipid probes, both +UV and -UV conditions were recorded to ensure that the obtained signal was indeed indicative of photo-generated covalent lipid-protein conjugates. Lipid images of -UV/+lipid probe type were not considered control images as they can contain true biological signal. For instance, the C16 bifunctional fatty acid can serve as a substrate for protein acyl transferases, resulting in bifunctional palmitoylated proteins. Overall, we found that -UV/+lipid images contained little more signal than control images, except (as expected) for PE, which is chemically fixable due to its primary amino group.

##### Ilastik models

We used the pixel classification workflow of Ilastik for segmenting the ER, lysosomes, lipid droplets and the mitochondria when labelled with AF-647. This workflow uses a Random Forest classifier that was trained using annotated images antibody images. The training was performed in 2D and using different frames of the Z-stack. For the remaining organelles (PM, Golgi, Mitochondria-488 and Lipid droplets) the Autocontext workflow was used. This workflow runs the Pixel Classification twice and shows the result of the first stage to the second stage of the process. For both workflows all pixel features and scales where used. Images were segmented using the Headless Model Operation in a High Performance Computing cluster.

##### Background correction and determination of bleedthrough using control images

During image acquisition, we noticed that a certain amount of bleedthrough from the 647 channel into the 561 channel occurred if a combination of Mouse-Tom20 primary and Goat anti-Mouse IgG AF Plus 647 secondary antibodies were used. The precise cause of this effect is unclear as it was neither standard bleedthrough from a shorter-wavelength to longer wavelength channel, nor observed for other primary antibodies combined with the same secondary antibody.

To correct for this effect, we built an image processing pipeline to quantify the extent of the bleedthrough signal and correct for it in lipid images. We first analyzed control images to determine the amount of background signal in the absence of bifunctional lipid probes by segmenting each frame of the 405, 488 and 647 channels using the corresponding Ilastik model according to the antibody labelling. A probability image of the segmentation for each frame was saved for each of the three channels. This probability image contains the probability of the segmented pixel originating from the desired structure according to the model training. The probability images were then used to analyze the control images. First a Gaussian filter was applied to each frame of the control image in the 561 channel using Scikit-Image[11]. A Minimum Threshold was applied to the blurred image (Scikit-Image), if this failed, a Mean Threshold was applied. A mask of the signal in the lipid channel (561 nm) was generated using the blurred image and the threshold to identify signal in the cell-covered area. If the number of pixels in this mask was below 10000, a Mean Threshold was applied to the blurred image to generate the mask.

Using the original pixel intensities inside the mask, the distribution of the pixel intensities were calculated and then fitted to a Genlogistic curve (Scipy[12]) to extract the c, loc and scale parameters. Using these parameters, a “noise image” was generated that had the same pixel intensity distribution as the control image. The noise image was then subtracted from the lipid channel of the control image (Extended Data Figure 4c).

This new, background-corrected image was used to calculate the bleed through of the red channel to the lipid channel. For this, the lipid image and the image of the red channel were segmented using the mask of the red channel. Only the pixels inside the masks were used to calculate the bleed through. A linear regression was then fitted to a plot of the pixel intensity in the red channel versus the pixel intensity in the lipid channel to obtain the slope and intercept of the line (Extended Data Figure 4d). Finally, for each control stack a csv file was saved for later use during background subtraction of the experimental images. The csv file contained the following columns: the labelling; the timepoint of the stack; the frame; the c, loc and scale parameters, the slope and the intercept of the regression line. The csv files of each stack where then merged together to be used for background subtraction of the experimental images.

##### Processing of lipid images

The 405, 488 and 647 channels were first segmented using the corresponding Ilastik model according to the antibody labelling. A probability image of the segmentation for each frame was saved for each of the three channels. This probability image contains the probability of the segmented pixel originating from the desired structure according to the model training. The probability images were then used to create binary organelle masks for determining the distribution of the lipid signal between organelles. A threshold of 0.3 probability was applied to the probability images in the 488 and 647 channels to generate segmentation masks. For the plasma membrane, imaged in the 405 channel, a probability of at least 0.8 for U-2 OS cells and 0.9 for HCT116 cells was used (Extended Data Figure 4e). The probability values need to be adjusted because of the different morphology of the cell lines and their interaction with the plate bottom. Background subtraction was then carried frame by frame using the values calculated for the control images. The mean of the saved parameters of all experiments (c, loc, scale, slope, intercept) was calculated per frame. With the parameters from the Genlogistic distribution, a noise image was generated which was then used to subtract the background signal from the lipid image (561 channel) to obtain the background-corrected image for all experiments that did not include labelled mitochondria in the 647 channel.

If the labeling included the mitochondria in the 647 channel, the values obtained for intercept and slope derived from the were used to generate an image of the expected bleedthrough in the absence of lipid signal. This image was subtracted from the lipid image obtained after the subtraction of the noise image and the resulting image used for lipid signal assignment (Extended Data Figure 4e).

#### Determination of cell covered area

After background subtraction, the cell covered area was determined. For this a maximum intensity projection of the plasma membrane channel (405) after background subtraction was used. To the max projection, a median filter was applied. Another image was generated by summing the maximum intensity projections of the green and the red channels' mask. To remove illumination artifacts, contrast limited adaptive histogram equalization (CLAHE) was applied to the max intensity projection image of the plasma membrane (Opencv[13]), followed by a median filter. A minimum plasma membrane intensity mask was generated by taking the mean intensity of the equalized image inside the plasma membrane mask at the frame with most plasma membrane and subtracting it by 2 standard deviations of the pixel intensity of the equalized image inside the plasma membrane mask at the same frame. Then the equalized plasma membrane image was subtracted by the min intensity mask to remove background from the plasma membrane image. A mean threshold was then applied to this image to generate a mask, called seed. In this seed mask, all values with 1 in the maximum intensity projection of the green and red channels were also included. An exclusion zone was determined applying a binary dilation to the seed. A background marker was defined as the pixel intensity in the 20$^{th}$ percentile of the equalized image intensity inside the seed region or as the 50$^{th}$ percentile if an ER stain is included. This background marker value was used to generate a rough background mask, by setting all values in the equalized image lower or equal than the background marker as 1 and all values higher as 0. The edges of the cells were then estimated using a Scharr filter. A markers image was made using the seed mask (value=2) and the background mask (value = 1). Segmentation of the total cell area was then performed using a Watershed algorithm. Finally small objects and holes were removed using morphology filters. This gave the maximum area that was occupied by the cells in a max intensity projection of the stacks, and therefore the region of interest for the image quantification.

#### Allocation of lipid signal to individual organelles

The image stack was then analyzed frame by frame using only the pixels inside the cell region. For this, masks were identified for the 405, 488 and 647 channels that contained no overlap between the other channels. Then, masks were made of the overlap between the three channels and between two channels. Using regions where the binary masks did not overlap, a kernel density estimation was performed (Scikit-learn[14]) to estimate the probability density function of the lipid signal for each segmented organelle. The distribution of the signal in the overlapping regions (identified in the different overlapping masks) was then determined by the calculated probability density function. That is, in a region were the 405 channel and the 488 channeled overlapped, the amount of lipid intensity attributed to each organelle was determined by the probability that the signal comes from the 405 or the 488 channel. To generate an image of the lipid signal corresponding to each channel the sum of the lipid signal of the regions where the marker channel did not overlap plus the estimate signal in the overlapping regions was taken. Using the images of the lipid distribution in the different organelles, the fraction of lipid signal assigned to a given organelle was calculated by dividing the sum of the signal intensity allocated to that organelle divided by the total lipid signal.

##### Lipid droplet analysis

We followed a different approach to quantify the amount of signal in lipid droplets for the different PC species. First, to avoid bleadtrhough of LipidSpot to other channels, we acquired two set of images: one before the addition of lipid spot and another set after. Image segmentation was then performed as previously described using Ilastik[10] to obtain a probability map for the lipid droplets. A stack was then made by using the 405, 488 and 561 channels of the first set and combining it with the 647 channel of the second set using image cross-correlation between the 405 channels of the two sets for alignment (OpenCV[13]).

The probability map was then used to make a mask of the lipid droplets (threshold probability: 0.2). A 3D label image for the droplets was made using Scikit-Image[11]. Background subtraction of the lipid image and signal allocation to the lipid droplets as performed as explained in the previous sections. Using the allocated signal to the lipid droplets and the 3D label image, the mean signal intensity of each droplet was calculated. We then estimated the enrichment of signal in the lipid droplets by dividing the signal of each droplet to the mean intensity lipid signal in all the cell.

#### Mass spectrometry data acquisition, analysis and data representation

Cells were seeded in a 6-well-plate up to 24 hours before the experiment. For the 4 min to 8 h timepoints, a density of $5 \times 10^5$ cells per well was used. For the 24 hours time point, $2.5 \times 10^5$ cells were seeded to keep the cells in exponential growth during the experimental time. Liposomes were prepared by mixing POPC : bifunctional lipid : cholesterol in a 1 : 2 : 1 ratio. After drying the chloroform solution containing the POPC, the bifunctional lipid and the cholesterol, the lipid film was hydrated using warmed filtered PBS to a final concentration of 3 mM total lipid (0.75 mM POPC, 1.5 mM Bifunctional lipid, 0.75 mM cholesterol).

The liposome stocks were diluted to 0.5 mM in McCoy's 5A medium and incubated for at least 30 min at 37 ºC in the presence of 4 mM methyl-alpha-cyclodextrin. This solution was then used for loading the lipid onto the cells. Before loading the liposomes, cells were washed 2 times with McCoy's 5A to remove FBS. 850 µL of the liposome/cyclodextrin solution was loaded onto the cells and incubated at 37 C for 4 min. The cells were then washed two times with McCoy's 5A. If the cells were going to be examined at a later time point (i.e. 8 h), the medium was removed and the cells were incubated at 37 C in McCoy's 5A supplemented with 10% FBS and penicillin-streptomycin (100 U/mL).

At the desired time point, the cells were taken from the incubator and placed on ice. The cell medium was removed and the cells were washed 2 times with 2 mL of ice-cold PBS. The last washed was removed and 1 mL of PBS was added. The cells were scrapped and collected in 1.5 mL Eppendorf tubes. The samples were centrifuged at 1200 x g for 5 min at 4 C. The PBS supernatant was removed and the cell pellet was resuspended in 500 µL of ice-cold 50% 2-propanol. A small amount of zirconia beads were added to the samples for homogeneization on a tissue lyzer (Qiagen). The homogeneization was performed two times at 30 Hz for 5 min at 4 ºC. were stored at -80 oC for further processing..

Cells were treated with 500µL isopropanol / H2O 1:1 (v/v) and lyzed using 0.5mm stainless steel beads in a TissueLyser II (Qiagen). An aliquot equivalent to 3 × 105 cells was transferred to the new 1.5 ml Eppendorf tube and dried in a Speedvac. For lipid extraction, 700 µL of the internal standard mixture (see Materials) in methyl-tert-butyl ester (MTBE) / MeOH (10:3; v/v) and 140 µL water were added and the mixture incubated for 1.5 hours at 4 ºC followed by 30 min centrifugation at 13.4 rpm at 4 ºC[15]. <Matyash et al, PMID: 18281723>

Before the analysis, 50 µL of organic (MTBE / methanol) layer were transferred to 96-well plate, dried in a vacuum desiccator and re-solubilized in 100 µL of MS mix solvent containing isopropanol/methanol/chloroform (4:2:1; v/v/v) with 7.5 mM ammonium formate. Samples were analysed by shotgun lipidomics on Q Exactive hybrid mass spectrometer (Thermo Fisher Scientific, Bremen) featured with robotic nanoflow ion source Triversa Nanomate HD (Advion Biosciences, Ithaca NY)[16] <ref PMID: 21634439> and Booster X2 data processing system (SpectroSwiss, Lausanne. ). RF S-lens level was set to 10% and capillary temperature to 275° C..

For ultra-high resolution FT MS with a target mass resolution Rs m/z = 200 of 106 (Full Width at Half Maximum, FWHM), analyte and dummy scans were combined to acquire 2 sec transients[17] <PMID: 38127459>. Spectra were acquired in targeted single ion monitoring (t-SIM) mode as successive 40 Th m/z mass isolation windows spaced by 20 Th[18] <ref. PMID: 29111682>.

For FT MSMS, regular 0.5s transients were recorded by Booster at Rs m/z=200 of 2.8 × 105. For each MS/MS experiment, the mass range of m/z 400 - 1000 was fragmented using 1Th precursor isolation window at normalized collision energy (NCE) of 15% and 22% in positive and negative mode, respectively.

Transients were converted to spectra by the PeakbyPeak software (SpectroSwiss) in aFT mode. Peak intensities were normalized to corresponding ion injection times. Transients with the same scan header were averaged. t-SIM spectra were processed by in-house software Simtrim that removed overlapping ranges in neighbouring mass windows[18] <ref. PMID: 29111682> and saved the resulting spectra in mzML format.

Lipids were identified by Lipidxplorer 1.2.4 software[19] <PMID: 22272252>: Chol, CE, TG, were quantified by UHR FTMS in positive mode; DG (FA neutral losses), PE (head group neutral loss fragment 141.019), PEO (NL 141), LPE (NL 141) by positive mode FT MS/MS; Cer, HexCer, SM, PC, PCO, LPC, LPCO, PI, LPI, CL by negative mode FT MS; PS (head group neutral loss fragment 87.032), PSO (NL 87), LPS (NL 87) by negative mode FT MS/MS. For analysing Y-modified lipids with LipidXplorer, additional MFQLs were generated for the all lipid classes to identify analytes with additional two nitrogen atoms of the diazirine group FTMS and FTMSMS spectra were re-calibrated using internal standards to improve mass accuracy. Further data analysis, including peak intensities normalization to internal standards, filtering for blank peaks and detection frequency within technical replicates was performed using Jupyter Notebook and Python 3.9.7

#### Mathematical modelling

The entire modeling process can be reproduced using the Matlab scripts provided separately in http://doi.org/21.11101/0000-0007-FCE3-D ('Optimization Toolbox' is needed for lsqcurvefit).

At t0 = 0 the total quantity of each lipid resides in the extracellular space ("external reservoir"), representing the situation where the loading solution has just been added ($E_{t0}$ = 100 %) and the plasma membrane ($M_{t0}$ = 0 %) is free of lipid. M can be populated by lipids from E with a specific rate ($k_E > 0$ ). Furthermore, lipid from M can be transported to an acceptor membrane ($A_{t0}$ = 0 %) with a specific rate ($k_A > 0$). Both processes can be combined to a set of ordinary differential equation (ODE) together with the initial conditions (initial value problem).

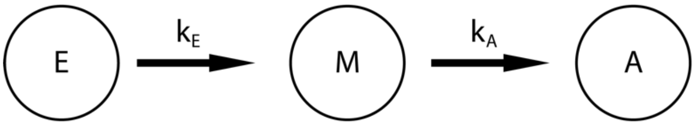

$$
\begin{aligned}
\frac{dE}{dt} &= -k_E \cdot E(t) \\
\frac{dM}{dt} &= k_E \cdot E(t) - k_A \cdot M(t) \\
\frac{dA}{dt} &= k_A \cdot M(t)
\end{aligned}
$$

The differential equations for the individual organelles were constructed successively (see Kinetic Models section and file:“DGLModel_MSNorm.m”)

To extract lipid transport rate constants, the ODE system - which was solved numerically using ode45 solver in Matlab R2020b - was embedded in a parameter optimization routine based on Matlabs nonlinear least-square solver (lsqcurvefit). The rapid lipid loading rate ($k_{Lipo \rightarrow PM}$) cannot be extracted from the dataset as the first datapoint is acquired after lipid loading is completed, so a fixed value (1000) was used to avoid overfitting.

After deriving the transport rates, the model was calculated (solid non-transparent lines in the model plots). The robustness of the model was then tested using a wild Bootstrap / Monte Carlo with normally distributed weights[20,21]. For the resampling, synthetic noise was generated in the same order of magnitude as the residuals of the fit. This was added to the model and the transport rate optimization was restarted (100 MC runs). The new models are shown with transparency (1% opacity) in the corresponding plots. This allowed us to check the reproducibility of the fit and to estimate the variance of the rates (standard deviation of the 100 MC runs).

As the model is a superposition of different transport processes, it is difficult to compare individual sub-processes directly with each other. To estimate the efficiency of vesicular and non-vesicular transport pathways from the PM to the endosomes (Endo) and to the ER, a model for these compartments was calculated (Extended Data Figure 7) that only includes the optimized transport rates (from the 100 MC runs) from the plasma membrane to Endosomes ($k_{PM \rightarrow Endo}$) and the ER ($k_{PM \rightarrow ER}$), respectively

### TMEM30 Knock-out

##### KO Generation using CRISPR/Cas9

The TMEM30A KO HCT116 cell line was generated by 1bp insertion into Exon3 leading to a 139 AA truncated protein. For the target region, the sequence was analyzed for appropriate guideRNA binding sites using the Geneious software and the CRISPOR design webtool (http://crispor.tefor.net/). Four combinations of guideRNA pairs were tested in advance, and the pair with the best performance was used to generate KO clones. Cuts were made using guideRNA pairs flanking the sequence

(GGCAACGTGTTTATGTATTATGGACTGTCTAATTTCTATCAAAACCATCGTCGTT ACGTGAAATCTCGAGATGATAGTCAACTAAATGGAGATTCTAGTGCTTTGCTT).

Cas9 protein, guideRNAs (ordered as crRNA, see Materials) and tracr-scaffoldRNA were obtained from IDT (Integrated DNA Technologies).

HCT116 cells were transfected with Cas9 ribonucleoprotein complexes using the Neon electroporation kit and device (Invitrogen). For the preparation of the crRNA-tracrRNA duplexes, RNA oligos (crRNA and tracrRNA) were mixed in equimolar amounts (100 µM). The mixes were incubated for 3 mins at 95 °C and subsequently slowly cooled to RT. For the formation of the Cas9 ribonucleoprotein complexes, crRNA-tracrRNA duplexes were mixed with Cas9 in a 1.4:1 molar ratio followed by a 15 min incubation at RT. Afterwards, the electroporation mixture was prepared by combining Cas9 ribonucleoprotein and cell suspension in electroporation buffer to a 12 µL final volume. From this mixture, 10 µL were used for electroporation that was performed with the following settings: 1530 V, 20 ms pulse width, 1 pulse. After 72h, cells were harvested and sorted as single cells into 96-well paltes (167008, Thermo Fisher Scientific) by FACS. Single-cell clones were screened for insertion by PCR using primers spanning the targeting region. Insertion was identified by Sanger Sequencing and RNA levels were investigated by qPCR.

##### KO validation qPCR

RNA was isolated using RNAeasy Mini Kit (74104, Qiagen) and RNA quality was assessed by RNA integrity number (RIN). RIN was determined with the Bioanalyzer 2100 (Agilent). Only RNA with a RIN >9 was used for qPCR. cDNA synthesis of 1 ug of total RNA was carried out for all samples (3 biological replicates for WT and KO) following the SuperScript II protocol (NEB, Ipswich, MA). Primers were designed using Primer Blast (NCBI, Bethesda, MD) with the following criteria: Tm 60 degrees, GC content between 40 and 80%, and amplicon length between 80 and 150 bp. GAPDH was used for housekeeping.

The Fast Start Essential SYBR Master Mix (Roche, Basel, Switzerland) was used for detection. All primers were tested and qualified for their efficiency, melting curve, and dependency of Cq from the template concentration according to MIQE guidelines. The plate was run on the Roche Light Cycler 96 (Roche, Basel, Switzerland) with the following thermal profile: heat activation at 95 degrees for 10 min, 40 cycles of melting (95 degrees for 10 sec), annealing (60 degrees for 10 sec) and elongation (72 degrees for 15 sec), and a melting curve. The data was analysed with the LC96 software (Roche, Basel, Switzerland) and plotted with R studio.

### Kinetic Models and model performance

Models 1a and 1b are "transport only" models (see Fig 2b, ED6, Supplementary Fig 11) that describe only the exchange lipids between cellular membranes and do not account for depletion of bifunctional lipids over time. The fraction of lipid signal (total assigned intensity normalized to 100%) in each organelle derived from imaging data is used as input and the sum does not change over time. Model 1a is the "minimal" model for the purpose of this study which contains the required minimum number of parameters required to fit the data.

Model 1a captures vesicular retrograde transport along the secretory pathway from the PM via Endosomes and the Golgi to the ER and retrograde non-vesicular lipid transport from the PM to the ER. Anterograde lipid transport is captured by a summary rate $k_{ER\rightarrow PM}ER(t)$ which encompasses all lipid transport (vesicular and non-vesicular). Lipid exchange between the mitochondria and other cellular membranes is assumed to occur only via the ER, based on the assumption that this is the main pathway for lipid flux into the mitochondria.

Model 1b also includes anterograde transport along the secretory pathway, that is from the ER to the PM via the Golgi and Endosomes (blue in Model 1a/b). Consequently, the rate $k_{ER\rightarrow PM}ER(t)$ describes non-vesicular transport from the ER to the PM in this model and is no summary rate.

Model 1 a/b

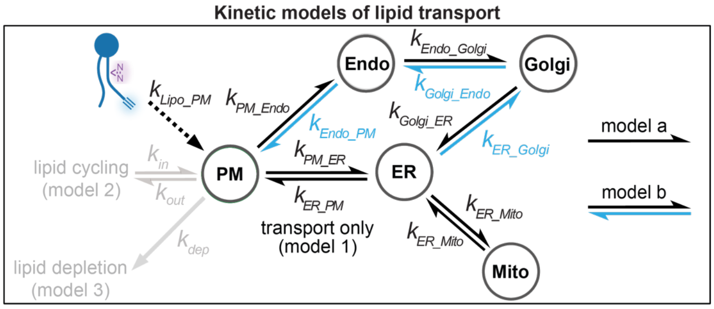

$$\frac{dLipo}{dt} = -k_{Lipo\rightarrow PM}Lipo(t)$$

$$\frac{dPM}{dt} = k_{Lipo\rightarrow PM}Lipo(t) + k_{ER\rightarrow PM}ER(t) - k_{PM\rightarrow Endo}PM(t) - k_{PM\rightarrow ER}PM(t) + k_{Endo\rightarrow PM}Endo(t)$$

$$\frac{dEndo}{dt} = k_{PM\rightarrow Endo}PM(t) - k_{Endo\rightarrow Golgi}Endo(t) + k_{Golgi\rightarrow Endo}Golgi(t) - k_{Endo\rightarrow PM}Endo(t)$$

$$\frac{dER}{dt} = k_{PM\rightarrow ER}PM(t) + k_{Mito\rightarrow ER}Mito(t) + k_{Golgi\rightarrow ER}Golgi(t) - k_{ER\rightarrow PM}ER(t) - k_{ER\rightarrow Mito}ER(t) - k_{ER\rightarrow Golgi}ER(t)$$

$$\frac{dMito}{dt} = k_{ER\rightarrow Mito}ER(t) - k_{Mito\rightarrow ER}ER(t)$$

$$\frac{dGolgi}{dt} = k_{Endo\rightarrow Golgi}Endo(t) - k_{Golgi\rightarrow ER}Golgi(t) + k_{ER\rightarrow Golgi}ER(t) - k_{Golgi\rightarrow Endo}Golgi(t)$$

Values for $k_{ER\rightarrow PM}ER(t)$ can thus not be directly compared between "a" and "b" models, whereas all other shared parameters describe the same transport steps and are directly comparable.

**Supplementary figure 11: Schematic description of model 1a/b and corresponding ordinary differential equations.**

Models 2a and 2b include information about the amount of bifunctional lipids in the cells from mass spectrometric measurements. To this end, the input data timepoint for model 1a/b (fraction of lipid signal in organelle) was scaled by a function accounting for the loss of bifunctional lipids over time. This function was derived by fitting a monoexponential decay to mass spectrometric determination of bifunctional lipid abundance in the cellular lipidome (Extended Data Figure 6. a). The input data for models 2a and 2b thus reflect the actual bifunctional lipid abundance and are not solely reflecting relative intensity differences between the different organelles.

To account for the decreasing total lipid amount, we included two additional rates (red in Supplementary Fig. 12) in the reaction network describing exchange of material between the extracellular space and the cell. The (re-)uptake reaction was required as the bifunctional lipid content did not converge to 0 during 24 h. The description of intracellular lipid transport pathways is the same as in Model 1a/b

Model 2 a/b (extensions of model 1 a/b)

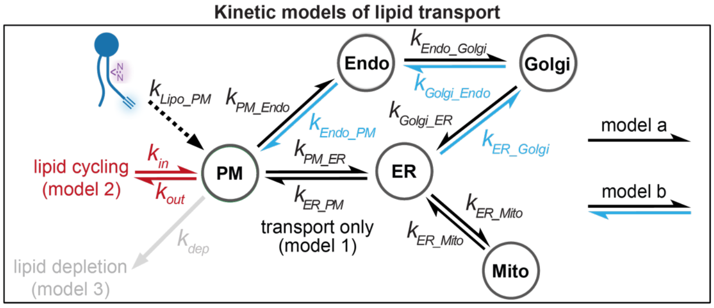

$$\frac{dPM}{dt} = \left(\frac{dPM}{dt}\right)_{model\ 1} - k_{out}PM(t) + k_{in}Medium(t)$$

$$\frac{dMedium}{dt} = k_{out}PM(t) - k_{in}Medium(t)$$

**Supplementary figure 12: Schematic description of models 2a/b and ordinary differential equations of additional rates.**

Models 3a and 3b are designed to minimize the influence of ongoing metabolism. To this end, the input data for model 1a/b (fraction of lipid signal in organelle) were scaled by a function accounting for the decrease of originally supplied bifunctional lipid species over time. This function was derived by fitting a biexponential decay to mass spectrometric determination of the abundance of the originally supplied bifunctional lipid species in the cellular lipidome. The input data for models 3a and 3b thus correspond to the abundance of the originally supplied species. To account for the decreasing abundance of the originally supplied species, we included a depletion rate in the reaction network, which summarily captures all lipid loss processes (metabolism, membrane shedding, dilution through lipid biogenesis during cell growth). The description of intracellular lipid transport pathways is the same as in Model 1a/b

Model 3 a/b (extensions of model 1 a/b)

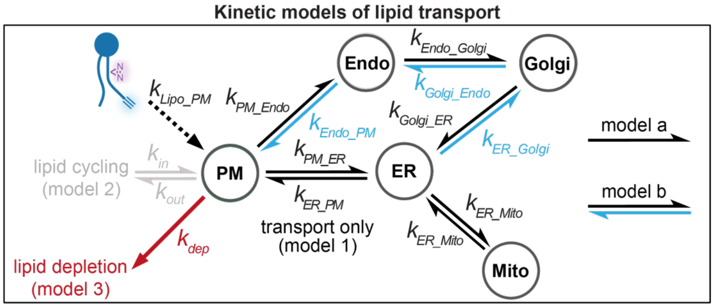

$$\frac{dPM}{dt} = \left(\frac{dPM}{dt}\right)_{model\ 1} - k_{dep}PM(t)$$

**Supplementary figure 13: Schematic description of models 3a/b and ordinary differential equations of additional rate.**

In all models, the rate constant for lipid transfer from the liposomes to the plasma membrane is not accessible for two reasons. First, there are no data points for the incorporation period (<2 minutes) and second, the exact fraction of total liposomal lipid transferred to the cells is unclear due to the sample preparation. Since all of the lipid later detected was rapidly transferred from the liposomes to the plasma membrane during the incorporation process, the same fast rate constant was fixed for all lipids.

Overall, all kinetic parameters were well identifiable for models 1-3a, which featured a summary rate for lipid transport in the anterograde direction. This was no longer true for models 1-3b which explicitly distinguished between vesicular and non-vesicular transport in the anterograde direction. Parameters differed significantly between model runs for many transport reactions, leading to large standard deviations (see Table S1). This is likely due to the fact that the additional parameters required to describe the individual transport steps lead to a model overparameterization. Models 1-3b can be understood as a test for the information content of the acquired dataset. While transport in the anterograde direction can be dissected into vesicular and non-vesicular routes, the same does not hold true for the anterograde direction. To do so, it would likely be necessary to chase probe localization starting from the ER and acquire a complementary dataset, which is technically not yet feasible for molecularly distinct lipid species.

Therefore, conclusions in the manuscript are therefore based on parameter sets based on models 1-3a.

#### Model for HCT116 WT and TMEM30A Knockout

A simplified model was used to quantify the change in PE transport in HCT116 wildtype and TMEM30A KO cells. For this model we only considered lipid transport from the liposomes to the plasma membrane and the reversible transport from the PM and the ER. Fluorescence intensity was normalized so that the sum of PM and ER signal is equal to 1.

$$\frac{dLiposome}{dt} = -k_{Lipo \to PM} Liposome(t)$$

$$\frac{dPM}{dt} = k_{Lipo \to PM} Liposomes(t) + k_{ER \to PM} ER(t) - k_{PM \to ER} PM(t)$$

$$\frac{dER}{dt} = k_{PM \to ER} PM(t) - k_{ER \to PM} ER(t)$$

### Supplementary Tables

**Supplementary Table 1:** Model 1 normalized to the sum of the lipid intensity

| | ***Lipids*** | | | | | | | |
|---|---|---|---|---|---|---|---|---|
| ***Rates (min$^{-1}$)*** Mean ± STD | PC(Y/16:0) | PC(Y/18:1) | PC(Y/20:4) | PC(16:0/Y) | PC(18:1/Y) | PC(20:4/Y) | PE(18:1/Y) | SM(Y) |
| ***PM→Endo*** | 0.00663±0.00224 | 0.00823±0.00415 | 0.0132±0.00607 | 0.00916±0.004 | 0.0125±0.00439 | 0.0213±0.00742 | 0.0324±0.00648 | 0.0219±0.00948 |
| ***Endo→Golgi*** | 0.0338±0.015 | 0.0421±0.0304 | 0.0829±0.0428 | 0.0448±0.0271 | 0.0659±0.0353 | 0.119±0.042 | 0.149±0.0445 | 0.0491±0.0242 |
| ***Golgi→ER*** | 0.0569±0.032 | 0.0608±0.0441 | 0.0986±0.0537 | 0.0303±0.0254 | 0.0368±0.0234 | 0.0582±0.0238 | 0.127±0.0353 | 0.136±0.0823 |
| ***PM→ER*** | 0.0292±0.00608 | 0.0396±0.0071 | 0.124±0.0179 | 0.0471±0.0157 | 0.0747±0.0216 | 0.195±0.0283 | 0.347±0.0458 | 0.141±0.0347 |
| ***ER→PM*** | 0.0262±0.0073 | 0.034±0.008 | 0.0806±0.0135 | 0.035±0.0159 | 0.0545±0.0227 | 0.125±0.0229 | 0.224±0.0491 | 0.146±0.0463 |
| ***ER→Mito*** | 0.252±0.431 | 0.12±0.47 | 0.38±0.605 | 0.0974±0.429 | 0.0246±0.0413 | 0.311±0.681 | 0.201±0.438 | 0.0177±0.0254 |
| ***Mito→ER*** | 2.62±4.29 | 0.554±2.17 | 2.53±4.01 | 0.386±1.72 | 0.0733±0.127 | 0.941±2.15 | 0.753±1.72 | 0.0593±0.0816 |

**Supplementary Table 2:** Model 1b normalized to the sum of the lipid intensity

| | ***Lipids*** | | | | | | | |
|---|---|---|---|---|---|---|---|---|
| ***Rates ($min^{-1}$)*** Mean ± STD | PC(Y/16:0) | PC(Y/18:1) | PC(Y/20:4) | PC(16:0/Y) | PC(18:1/Y) | PC(20:4/Y) | PE(18:1/Y) | SM(Y) |
| ***PM→Endo*** | 0.0291±0.0159 | 0.077±0.033 | 0.0747±0.0491 | 0.0187±0.0123 | 0.019±0.019 | 0.0409±0.0315 | 0.0803±0.0246 | 0.0061±0.0089 |
| ***Endo→PM*** | 0.1845±0.1486 | 0.6585±0.407 | 0.3679±0.2965 | 0.1037±0.1508 | 0.1754±0.2313 | 0.4289±0.3531 | 0.5067±0.3479 | 0.1615±0.0886 |
| ***Endo→Golgi*** | 0.0534±0.186 | 0.781±0.3295 | 0.2636±0.3671 | 0.5455±0.2398 | 0.4804±0.262 | 0.6107±0.3172 | 0.1938±0.3036 | 0.0415±0.0772 |
| ***Golgi→Endo*** | 0.0296±0.0902 | 0.3772±0.2307 | 0.0994±0.156 | 0.8418±0.2888 | 0.812±0.313 | 0.7386±0.3275 | 0.0582±0.1298 | 0.4731±0.2705 |
| ***Golgi→ER*** | 0.1324±0.2282 | 0.1183±0.2174 | 0.5318±0.3608 | 0.142±0.2455 | 0.1461±0.2617 | 0.166±0.2718 | 0.3344±0.2854 | 0.1873±0.3097 |
| ***ER→Golgi*** | 0.0265±0.0416 | 0.0267±0.0346 | 0.0932±0.0749 | 0.0108±0.0195 | 0.0192±0.0237 | 0.0216±0.0227 | 0.0516±0.0439 | 0.0844±0.0481 |
| ***PM→ER*** | 0.0424±0.0117 | 0.0575±0.0174 | 0.1903±0.0409 | 0.0244±0.0081 | 0.0374±0.008 | 0.115±0.0177 | 0.3177±0.0388 | 0.1357±0.0289 |
| ***ER→PM*** | 0.025±0.0115 | 0.0295±0.0206 | 0.1175±0.0328 | 0.0185±0.0098 | 0.0199±0.0141 | 0.0578±0.0235 | 0.1891±0.0454 | 0.0631±0.0497 |
| ***ER→Mito*** | 0.0543±0.1723 | 0.067±0.1805 | 0.2636±0.2866 | 0.0977±0.248 | 0.077±0.1909 | 0.3271±0.3913 | 0.0597±0.0455 | 0.0249±0.0438 |
| ***Mito→ER*** | 3.3854±0.4471 | 2.6372±0.3773 | 2.7642±0.2151 | 7.503±1.7698 | 4.1293±0.5777 | 5.8881±1.2214 | 3.4777±0.3541 | 3.3666±0.6608 |

**Supplementary Table 3:** Model 2 scaled by the bifunctional lipid content

| | *Lipids* | | | | | | | |
|---|---|---|---|---|---|---|---|---|
| ***Rates (min$^{-1}$)*** Mean ± STD | PC(Y/16:0) | PC(Y/18:1) | PC(Y/20:4) | PC(16:0/Y) | PC(18:1/Y) | PC(20:4/Y) | PE(18:1/Y) | SM(Y) |
| ***PM→Endo*** | 0.0074±0.0012 | 0.0101±0.0025 | 0.0171±0.0068 | 0.0042±0.0011 | 0.0052±0.0021 | 0.0054±0.0013 | 0.0293±0.0065 | 0.0086±0.0017 |
| ***Endo→Golgi*** | 0.0388±0.0083 | 0.0574±0.0217 | 0.116±0.0594 | 0.0228±0.0097 | 0.0246±0.0139 | 0.032±0.0108 | 0.1482±0.0555 | 0.0262±0.0082 |
| ***Golgi→ER*** | 0.03±0.0089 | 0.0417±0.02 | 0.0675±0.0328 | 0.0431±0.0533 | 0.1411±0.9964 | 0.0379±0.016 | 0.1393±0.0484 | 0.0761±0.066 |
| ***PM→ER*** | 0.065±0.0155 | 0.112±0.0243 | 0.1718±0.0234 | 0.0351±0.0053 | 0.053±0.0088 | 0.1178±0.0151 | 0.4154±0.0552 | 0.154±0.02 |
| ***ER→PM*** | 0.0674±0.0218 | 0.1145±0.0318 | 0.147±0.0265 | 0.0446±0.0089 | 0.0682±0.0153 | 0.0942±0.0172 | 0.3432±0.0677 | 0.224±0.0423 |
| ***ER→Mito*** | 0.0075±0.0024 | 0.0095±0.0048 | 0.0191±0.0188 | 0.0051±0.0036 | 0.006±0.0038 | 0.0087±0.0089 | 0.0706±0.0732 | 0.3204±0.5266 |
| ***Mito→ER*** | 0.0241±0.011 | 0.026±0.0166 | 0.0512±0.059 | 0.031±0.0266 | 0.0247±0.0227 | 0.0465±0.0674 | 0.2929±0.3398 | 1.7001±2.8706 |
| ***out*** | 0.0087±0.0006 | 0.0229±0.0014 | 0.0824±0.0043 | 0.0118±0.0004 | 0.0191±0.0011 | 0.0528±0.0029 | 0.0217±0.0022 | 0.0435±0.0015 |
| ***in*** | 0.0002±0.0003 | 0.0026±0.0007 | 0.0107±0.0014 | 0.0001±0.0002 | 0.0012±0.0006 | 0.0088±0.001 | 0.0038±0.001 | 0.0017±0.0004 |

**Supplementary Table 4:** Model 2b scaled by the bifunctional lipid content

| | ***Lipids*** | | | | | | | |
|---|---|---|---|---|---|---|---|---|
| ***Rates (min$^{-1}$)*** Mean ± STD | PC(Y/16:0) | PC(Y/18:1) | PC(Y/20:4) | PC(16:0/Y) | PC(18:1/Y) | PC(20:4/Y) | PE(18:1/Y) | SM(Y) |
| ***PM→Endo*** | 0.0276±0.0193 | 0.0446±0.0294 | 0.0773±0.0454 | 0.0124±0.0097 | 0.006±0.0031 | 0.0142±0.0129 | 0.0774±0.0182 | 0.0049±0.0044 |
| ***Endo→PM*** | 0.2414±0.2005 | 0.4133±0.3069 | 0.6063±0.3464 | 0.2171±0.16 | 0.0186±0.0381 | 0.1575±0.1865 | 0.6584±0.2865 | 0.0724±0.0697 |
| ***Endo→Golgi*** | 0.022±0.1106 | 0.056±0.1438 | 0.0201±0.0365 | 0.0448±0.0948 | 0.0314±0.072 | 0.023±0.1008 | 0.1339±0.2188 | 0.146±0.1056 |
| ***Golgi→Endo*** | 0.021±0.0583 | 0.0243±0.0643 | 0.0163±0.0234 | 0.918±0.1964 | 0.0254±0.1204 | 0.0201±0.0886 | 0.0776±0.1722 | 0.8853±0.244 |
| ***Golgi→ER*** | 0.0535±0.1516 | 0.0408±0.111 | 0.0178±0.0158 | 0.7807±0.2832 | 0.0188±0.0145 | 0.0188±0.0172 | 0.2445±0.2035 | 0.1892±0.2494 |
| ***ER→Golgi*** | 0.013±0.0238 | 0.0102±0.0193 | 0.0071±0.0047 | 0.0409±0.0255 | 0.0027±0.0043 | 0.0044±0.0028 | 0.0491±0.0341 | 0.0353±0.0225 |
| ***PM→ER*** | 0.0571±0.0133 | 0.1046±0.0203 | 0.1642±0.0188 | 0.0324±0.005 | 0.0506±0.0077 | 0.1175±0.0142 | 0.3925±0.0463 | 0.1359±0.0156 |
| ***ER→PM*** | 0.0489±0.017 | 0.0989±0.0325 | 0.1196±0.0216 | 0.0166±0.0169 | 0.0594±0.0146 | 0.089±0.0168 | 0.2959±0.0648 | 0.1605±0.0445 |
| ***ER→Mito*** | 0.008±0.0037 | 0.0108±0.0092 | 0.0172±0.0097 | 0.0049±0.0017 | 0.0067±0.0036 | 0.008±0.0031 | 0.0257±0.0155 | 0.0061±0.0049 |
| ***Mito→ER*** | 2.9007±0.7059 | 2.4071±0.5122 | 2.3432±0.2497 | 3.5396±0.8676 | 2.9077±0.6549 | 3.5539±0.5044 | 3.4511±0.4797 | 2.5242±1.6395 |
| ***out*** | 0.009±0.0007 | 0.024±0.0022 | 0.0875±0.0042 | 0.012±0.0006 | 0.0194±0.0012 | 0.0551±0.0032 | 0.0212±0.0024 | 0.044±0.0021 |
| ***in*** | 0.0003±0.0005 | 0.003±0.0011 | 0.0112±0.001 | 0.0002±0.0003 | 0.0011±0.0006 | 0.0093±0.001 | 0.0035±0.0012 | 0.0017±0.0005 |

**Supplementary Table 5:** Model 3 scaled by the bifunctional lipid content

| | ***Lipids*** | | | | | | | |
|---|---|---|---|---|---|---|---|---|
| ***Rates (min$^{-1}$)*** Mean ± STD | PC(Y/16:0) | PC(Y/18:1) | PC(Y/20:4) | PC(16:0/Y) | PC(18:1/Y) | PC(20:4/Y) | PE(18:1/Y) | SM(Y) |
| ***PM→Endo*** | 0.0116±0.0023 | 0.0127±0.003 | 0.0146±0.005 | 0.0077±0.0013 | 0.0081±0.0023 | 0.004±0.0011 | 0.0283±0.0078 | 0.0085±0.0023 |
| ***Endo→Golgi*** | 0.0573±0.0146 | 0.0553±0.0146 | 0.0913±0.0453 | 0.0396±0.0102 | 0.0399±0.018 | 0.0208±0.0102 | 0.1438±0.0645 | 0.0206±0.0096 |
| ***Golgi→ER*** | 0.049±0.017 | 0.0401±0.0147 | 0.0578±0.0278 | 0.0644±0.0178 | 0.0626±0.0305 | 0.0239±0.0153 | 0.1333±0.0427 | 0.11±0.633 |
| ***PM→ER*** | 0.0744±0.0179 | 0.1182±0.0278 | 0.1954±0.0279 | 0.0401±0.0065 | 0.0595±0.0128 | 0.126±0.0145 | 0.4696±0.0608 | 0.1623±0.0334 |
| ***ER→PM*** | 0.087±0.027 | 0.1273±0.0388 | 0.1785±0.0339 | 0.0556±0.0114 | 0.0856±0.0234 | 0.1228±0.0188 | 0.4354±0.075 | 0.2342±0.069 |
| ***ER→Mito*** | 0.0284±0.0259 | 0.1829±0.558 | 0.0936±0.0657 | 0.1261±0.4073 | 0.019±0.0332 | 0.0049±0.0023 | 0.0565±0.0296 | 0.4126±0.703 |
| ***Mito→ER*** | 0.0969±0.0946 | 0.5437±1.6557 | 0.3115±0.2267 | 0.7401±2.3576 | 0.0868±0.204 | 0.0201±0.0148 | 0.2373±0.1379 | 1.8638±3.1903 |
| ***dep*** | 0.0116±0.0009 | 0.0238±0.0017 | 0.0653±0.0032 | 0.0162±0.0006 | 0.0257±0.0012 | 0.0549±0.0021 | 0.0284±0.0018 | 0.0413±0.0021 |

**Supplementary Table 6:** Model 3b scaled by the bifunctional lipid content

| | ***Lipids*** | | | | | | | |
|---|---|---|---|---|---|---|---|---|
| ***Rates (min$^{-1}$)*** Mean ± STD | PC(Y/16:0) | PC(Y/18:1) | PC(Y/20:4) | PC(16:0/Y) | PC(18:1/Y) | PC(20:4/Y) | PE(18:1/Y) | SM(Y) |
| ***PM→Endo*** | 0.024±0.0127 | 0.0338±0.0237 | 0.0929±0.0418 | 0.0175±0.0113 | 0.0067±0.0047 | 0.0201±0.0176 | 0.0731±0.0178 | 0.0065±0.0063 |
| ***Endo→PM*** | 0.1961±0.1609 | 0.2977±0.2496 | 0.6932±0.3095 | 0.2844±0.1661 | 0.0535±0.0781 | 0.3156±0.3304 | 0.646±0.2206 | 0.0733±0.0943 |
| ***Endo→Golgi*** | 0.0954±0.2113 | 0.0788±0.1532 | 0.0539±0.0502 | 0.5908±0.3471 | 0.3136±0.3299 | 0.0137±0.0202 | 0.0657±0.1759 | 0.3758±0.2466 |
| ***Golgi→Endo*** | 0.063±0.1104 | 0.0411±0.0718 | 0.0069±0.0125 | 0.6965±0.3098 | 0.3266±0.322 | 0.0066±0.0084 | 0.0442±0.1025 | 0.6732±0.3453 |
| ***Golgi→ER*** | 0.1636±0.2277 | 0.0477±0.0942 | 0.015±0.0149 | 0.05±0.1444 | 0.074±0.0883 | 0.0057±0.0066 | 0.0255±0.0416 | 0.1082±0.1233 |
| ***ER→Golgi*** | 0.0538±0.0677 | 0.0261±0.0439 | 0.0034±0.0044 | 0.0383±0.0287 | 0.025±0.0231 | 0.0023±0.0017 | 0.0114±0.0122 | 0.0475±0.0394 |
| ***PM→ER*** | 0.0718±0.0151 | 0.1148±0.0205 | 0.1792±0.0185 | 0.0376±0.0065 | 0.0625±0.0095 | 0.1271±0.014 | 0.4461±0.0572 | 0.1419±0.0253 |
| ***ER→PM*** | 0.0645±0.029 | 0.1035±0.0313 | 0.1499±0.0231 | 0.0139±0.0217 | 0.0754±0.0281 | 0.1191±0.019 | 0.3784±0.0745 | 0.1736±0.0714 |
| ***ER→Mito*** | 0.0572±0.1042 | 0.0631±0.1177 | 0.0316±0.0382 | 0.1355±0.2641 | 0.0213±0.0207 | 0.0055±0.001 | 0.0684±0.0501 | 0.0115±0.0104 |
| ***Mito→ER*** | 2.6348±0.3873 | 1.9245±0.3363 | 2.0827±0.5305 | 3.4506±0.5468 | 3.0218±0.5378 | 2.3816±0.5583 | 3.3044±0.6132 | 1.4867±0.7754 |
| ***dep*** | 0.0116±0.0009 | 0.0235±0.0018 | 0.0672±0.0028 | 0.0162±0.0007 | 0.0258±0.0011 | 0.0563±0.0021 | 0.0285±0.0017 | 0.042±0.0023 |

### Chemical synthesis

#### General synthetic procedures

Solvents for flash chromatography were obtained from VWR and dry solvents from Sigma. Deuterated solvents were obtained from Deutero GmbH, Karlsruhe, Germany. TLC was performed on precoated plates of silica-gel (Merck, 60 F254) using UV-light (254 nm or 365 nm) or staining solution of phosphomolybdic acid in EtOH (10g phosphomolybdic acid in EtOH) for analysis. Preparative column chromatography was performed using silica gel from Merck (silica 60, grain size 0.063-0.200 nm) with a pressure of 1 bar. $^{1}H$-, $^{13}C$- and $^{31}P$-NMR spectra were measured on 400 MHz Advance$^{TM}$ III HD Nanobay Bruker spectrometer. Chemical shift of $^{1}H$- and $^{13}C$-NMR spectra are referenced indirectly to tetramethylsilane. J values are given in Hz and chemical shifts in ppm. Splitting patterns are mentioned as follows: s, singlet; d, doublet; t, triplet; q, quartet; m, multiplet; $m_c$, centered multiplet. $^{13}C$-NMR spectra were broadband hydrogen decoupled. Mass spectra (ESI) were accorded using a QExactive instrument (Thermo Fisher Scientific) equipped with a robotic nanoflow ion source.

#### Synthesis of the bifunctional fatty acid

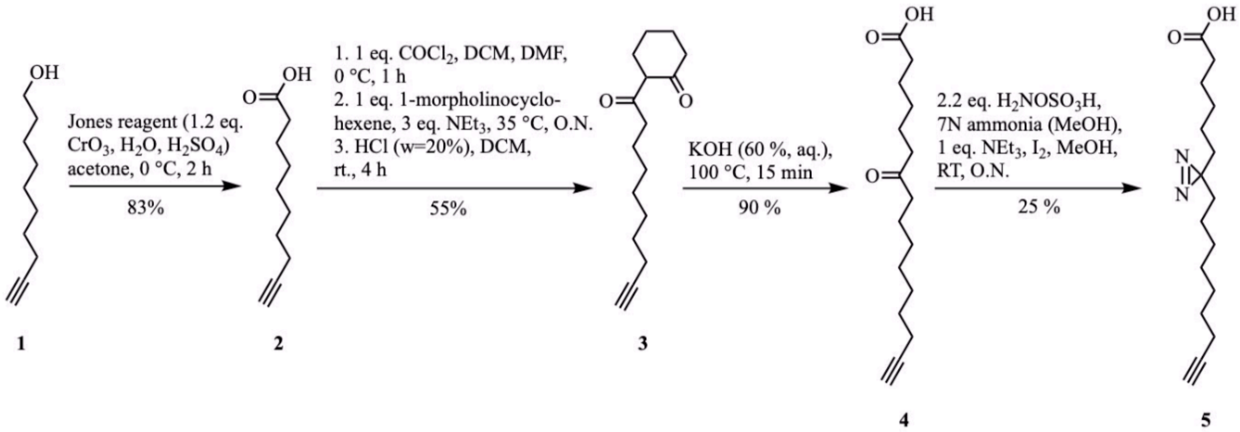

Scheme A: Synthesis of the bifunctional fatty acid 5. Dec-9-ynoic acid (2) was synthesized according to literature protocols[22]. The analytical data were in accordance with those previously reported[23,24]. All other protocols and analytical data are listed below.

### 2-(Dec-9-ynoyl)cyclohexan-1-one (3)

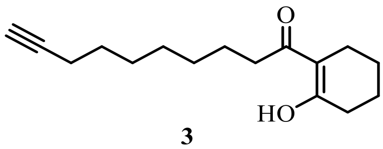

**3**

A solution of Dec-9-ynoic acid (**2**) (5.30 g, 31.5 mmol, 1.0 eq.) in 30 ml dry DCM was treated with 6 drops of dry DMF under an inert gas atmosphere. The reaction mixture was cooled to 0 °C and oxalyl chloride (4.14 g, 2.8 ml, 32.6 mmol, 1.0 eq.) was added dropwise. After removing the ice bath, the reaction mixture was stirred for 1 h at room temperature. The resulting yellow acid chloride solution was added slowly to a solution of 5.14 g of 1-morpholinocyclohexene (30.7 mmol, 1.0 eq.) and 13.2 ml $NEt_3$ (9.60 g, 94.9 mmol, 3.0 eq.) in 35 ml dry DCM placed at 35 °C. Afterwards, the reaction mixture was stirred over night at room temperature. Subsequently, 70 ml aqueous HCl (w/w=20%) and 120 ml chloroform were added and the mixture was stirred for further 4 h at room temperature. Organic and aqueous layer were separated and the organic phase was washed 3x with water. The combined aqueous layers were reextracted with DCM. The combined organic layers were dried over $Na_2SO_4$ and the solvent removed under reduced pressure. The obtained light brownish oil was purified by flash chromatography twice (CyHex/EtOAc, 95:5). 2-(Dec-9-ynoyl)cyclohexan-1-one (**3**) was isolated as a colourless liquid.

$^1$H NMR (400 MHz, $CD_3OD$) δ = 2.44 (t, $J$ = 7.4 Hz, 2H), 2.40 – 2.25 (m, 4H), 2.21 – 2.10 (m, 3H), 1.69 ($m_c$, 4H), 1.64 – 1.55 (m, 2H), 1.55 – 1.46 (m, 2H), 1.46 – 1.25 (m, 6H) ppm.

$^{13}$C NMR{$^1$H} (101 MHz, $CD_3OD$) δ = 203.44, 181.84, 107.90, 85.03, 69.40, 37.95, 31.81, 30.31, 30.04, 29.65, 29.64, 25.32, 24.85, 23.95, 22.74, 19.00 ppm.

HR-MS (ESI negative) m/z calculated for $C_{16}H_{24}O_2$: 248.17763; found: 247.170 $[M-H]^-$.

Yield: 4.3 g (17.31 mmol, 55 %).

### 7-Oxohexadec-15-ynoic acid (4)

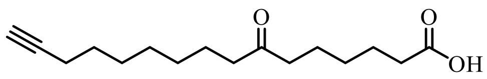

**4**

Neat 2-(Dec-9-ynoyl)cyclohexan-1-one (**3**) (6.30 g, 25.4 mmol, 1 eq.) was heated to 100 °C and treated with 20 ml of a 60% (w/v) aqueous KOH solution. After 15 min the oil bath was removed and the reaction mixture was allowed to cool down to room temperature. The reaction mixture was diluted with 200 ml water, neutralized with saturated aqueous HCl solution and extracted with chloroform (3x). The organic layer was dried over. $Na_2SO_4$ and the solvent removed under reduced pressure. The crude product was purified by flash chromatography (DCM/MeOH, 97:3). The product was isolated as a colourless solid.

$^1$H NMR (400 MHz, $CD_3OD$) δ = 2.46 (t, *J* = 7.2 Hz, 2H), 2.45 (t, *J* = 7.3 Hz, 2H), 2.28 (t, *J* = 7.4 Hz, 2H), 2.21 – 2.12 (m, 3H), 1.67 – 1.45 (m, 8H), 1.45 – 1.37 (m, 2H), 1.37 – 1.23 (m, 6H) ppm.*

*Signals at 2.46 ppm and 2.45 ppm overlap.

$^{13}$C NMR{$^1$H} (101 MHz, $CD_3OD$) δ = 214.18, 179.69, 85.05, 69.36, 43.43, 43.32, 36.55, 30.17, 30.01, 29.98, 29.64, 29.60, 26.53, 24.83, 24.63, 18.97 ppm.

HR-MS (ESI positive) m/z calculated for $C_{16}H_{26}O_3$: 266.18819; found: 267.20$[M+H]^+$.

Yield: 6.10 g (22.9 mmol, 90 %).

### 6-(3-(non-8-yn-1-yl)-3H-diazirin-3-yl)hexanoic acid (5)

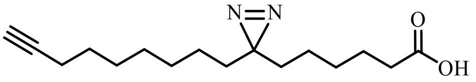

**5**

Molecular sieves (3Å, powder) were activated in a round bottom flask under vacuum using a heat gun. A 7N solution of ammonia in MeOH (20 ml) was added under inert gas atmosphere. After adding a solution of 1.18 g of 7-Oxohexadec-15-ynoic acid (4.43 mmol, 1.0 eq.) in 14 ml dry MeOH the reaction mixture was stirred for 4 h at room temperature. Subsequently, 1.08 g of $H_2NOSO_3H$ (9.55 mmol, 2.2 eq.) were dried under high vacuum for 3 h, then dissolved in 11 ml dry MeOH and added to the reaction mixture in a dropwise fashion at 0 °C. The reaction mixture was stirred over night at RT. Molecular sieves were removed by filtration. The solvents were removed under reduced pressure and the residue was dissolved in 20 ml MeOH and cooled down to 0 °C. 6 ml $NEt_3$ (4.43 mmol, 1 eq.) were added dropwise and the mixture was stirred for 1 h at 0 °C. Subsequently, small portions of iodine were added at 0 °C until the mixture remained brownish. The excess iodine was removed by addition of saturated $Na_2S_2O_3$-solution. The reaction mixture was extracted with DCM (3x), the combined organic layers dried over $Na_2SO_4$ and the solvent removed under reduced pressure. The crude product was purified by flash chromatography (DCM/MeOH, 97:3) and HPLC using a Macherey Nagel VP 250/21 Nucleodur C18 Pyramid column at 10 ml/min eluting with a gradient. The solvent system used was A (75 % MeOH, 25 %water, + 4 % AcOH) and B (50 % MeCN, 40 % *i*-propanol, 10 % MeOH, + 4 % AcOH). Gradient: 0-2 min: 0-45 % B, 2-8 min: 45-75 % B, 8-11 min: 75-100 % B, 11-27 min: 100-0% B, 27-30 min: 0 % B. The retention time of the product was found to be 10.5 min. The bifunctional fatty acid **5** was isolated as a yellowish oil.

$^1$H NMR (400 MHz, $CDCl_3$) δ = 2.33 (t, $J$ = 7.4 Hz, 2H), 2.17 (td, $J$ = 7.0, 2.4 Hz, 2H), 1.94 ($m_c$, 1H), 1.59 (tt, $J_1$ = 7.6 Hz, $J_2$ = 7.6 Hz, 2H), 1.51 (tt, $J_1$ = 7.2 Hz, $J_2$ = 7.2 Hz, 2H), 1.43 – 1.16 (m, 12H), 1.16 – 0.98 (m, 4H) ppm.

$^{13}$C NMR{$^1$H} (101 MHz, $CDCl_3$) δ = 178.77, 84.82, 68.28, 33.75, 33.00, 32.85, 29.20, 28.99, 28.86, 28.73, 28.69, 28.52, 24.55, 23.92, 23.67, 18.50 ppm.

HR-MS (ESI negative) m/z calculated for $C_{16}H_{26}N_2O_2$: 278.19943; found: 277.19 $[M-H]^-$.

Yield: 310 mg (1.11 mmol, 25 %).

### Synthesis of Phosphatidylethanolamine(18:1/Y)

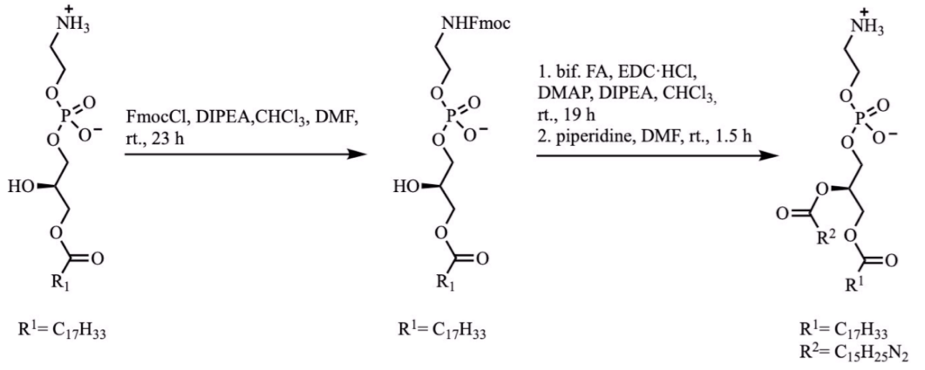

Scheme B: Synthesis of the bifunctional PE(18:1/Y).

***(2R)-3-(((2-((((9H-fluoren-yl)methoxy)carbonyl)amino)ethoxy)(hydroxy)phosphoryl) oxy)-2-hydroxypropyl oleate (7)***

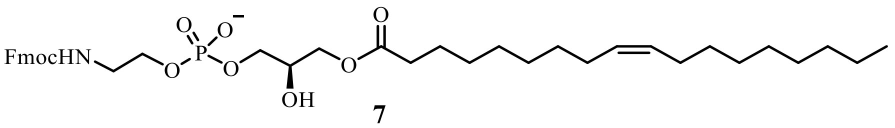

**7**

A solution of 200 mg of *sn1*-Oleoyl-lyso-PE (**6**) (417 µmol, 1.0 eq.) in 6 ml dry chloroform was treated with 107 µl of DIPEA (79.1 mg, 614 µmol, 1.5 eq.) for 15 minutes at RT under an inert atmosphere. Subsequently, a solution of 116 mg FmocCl (448 µmol, 1.1 eq.) in 3 ml dry chloroform was added and the reaction mixture was stirred for 22 h under inert conditions. The solvent was removed under reduced pressure and the crude product was purified by flash chromatography ($CHCl_3$/MeOH/$H_2O$, 65:35:2). Fmoc- *sn1*-Oleoyl-lyso-PE (**7**) was isolated as a white solid.

$^1$H NMR (400 MHz, $CD_3OD$) δ = 7.77 (d, $J$ = 7.5 Hz, 2H), 7.64 (d, $J$ = 7.4 Hz, 2H), 7.36 (dd, $J_1$ = 7.4 Hz, $J_2$ = 7.4 Hz ), 7.29 (dd, $J_1$ = 7.4 Hz, $J_2$ = 7.4 Hz ), 5.31 ($m_c$, 2H), 4.28 (d, $J$ = 14.6 Hz, 2H), 4.24 – 4.01 (m, 3H), 4.00 – 3.82 (m, 5H), 3.36 (t, $J$ = 5.2 Hz, 2H), 2.28 (t, $J$ = 7.5 Hz, 2H), 2.05 – 1.89 (m, 4H), 1.54 ($m_c$, 2H), 1.40 – 1.19 (m, 20H), 0.88 (t, $J$ = 6.8 Hz, 3H) ppm.

$^{13}$C NMR{$^1$H} (101 MHz, $CD_3OD$) δ = 175.30, 158.85, 145.28, 142.55, 130.84, 130.77, 128.77, 128.15, 126.22, 120.92, 69.86, 67.92, 67.56, 66.20, 65.47, 42.64, 34.86, 33.04, 30.82, 30.78, 30.59, 30.43, 30.32, 30.30, 30.18, 28.11, 25.93, 23.72, 14.48 ppm.*[1,2]

*[1] Signals at 69.8, 67.5 65.4 and 42.6 ppm appear as two peaks likely due to $^{13}$C-$^{31}$P coupling (see additional zoomed spectra A and B in NMR characterization section at the end of the Supplementary Information document).

*[2] Signals between 30.2 ppm and 30.8 ppm overlap, not all signals are resolved individually.

$^{31}$P NMR{$^1$H} (162 MHz, $CD_3OD$) δ = -1.41 ppm.

HR-MS (ESI negative) m/z calculated for $C_{38}H_{56}NO_9P$: 701.36927; found: 700.363 $[M-H]^-$.

Yield: 257 mg (366 µmol, 88 %).

***PE(18:1/Y) (8)***

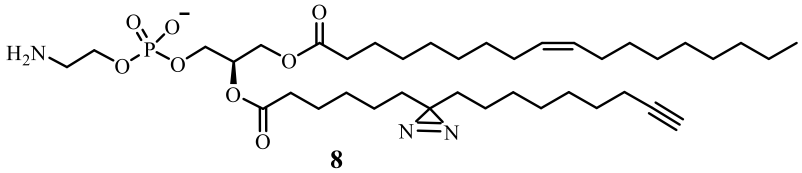

**8**

A solution of 57.0 mg of the bifunctional fatty acid **5** (186 µmol, 1.0 eq.) in 3 ml dry DMF was treated with 64.4 mg EDC·HCl (336 µmol, 1.8 eq.), 11.8 mg DMAP (96.6 µmol, 0.5 eq.) and 107 µl of DIPEA (614 µmol, 3.3 eq.) for 12 minutes at RT. After adding a solution of 133 mg Fmoc-*sn1*-Oleoyl-lyso-PE (**7**) (190 µmol, 1.0 eq.) in 3 ml dry DMF and 5 ml dry chloroform, the reaction mixture was stirred for 19 h under inert gas atmosphere. The solvent was removed under reduced pressure, and most impurities were removed by flash chromatography ($CHCl_3$/MeOH, 7:3). The intermediate Fmoc-protected bifunctional PE was difficult to fully purify in an efficient manner, the obtained product-containing fractions were directly used for the next step without further purification.

The partially purified intermediate was dissolved in 3 ml dry DMF and 2 drops of piperidine were added. The reaction was stopped after 1.5 h by removal of the solvent under reduced pressure. The crude product was purified by flash chromatography ($CHCl_3$/MeOH/$H_2O$, 65:35:2). PE(18:1/Y) (**8**) was isolated as yellowish oil.

$^{1}$H NMR (400 MHz, $CD_3OD$) δ = 5.35 ($m_c$, 2H), 5.23 ($m_c$, 1H), 4.44 (dd, $J$ = 12.0, 3.2 Hz, 1H), 4.18 (dd, $J$ = 12.0, 6.7 Hz, 1H), 4.04 ($m_c$, 2H), 4.00 (t, $J$ = 5.9 Hz, 2H), 3.17 (t, $J$ = 4.9 Hz, 2H), 2.33 (t, $J$ = 7.3 Hz, 2H), 2.32 (t, $J$ = 7.4 Hz, 2H), 2.21 – 2.08 (m, 3H), 2.09 – 1.99 (m, 4H), 1.69 – 1.53 (m, 4H), 1.53 – 1.45 (m, 2H), 1.43 – 1.22 (m, 32H), 1.16 – 1.02 (m, 4H), 0.91 (t, $J$ = 6.7 Hz, 3H) ppm.*[1]

*[1] Signals (triplets) at 2.33 ppm and 2.32 ppm overlap.

$^{13}$C NMR{$^{1}$H} (101 MHz, $CD_3OD$) δ = 174.92, 174.45, 130.93, 130.79, 85.02, 71.92, 69.44, 64.93, 63.60, 62.95, 41.71, 41.65, 34.90, 33.83, 33.74, 33.07, 30.85, 30.84, 30.62, 30.46, 30.36, 30.33, 30.22, 30.19, 29.97, 29.69, 29.63, 29.50, 28.15, 26.00, 25.75, 24.87, 24.66, 23.75, 18.99, 14.48 ppm.*[2,3]

*[2] Signals at 71.9, 64.9, 62.9 and 33.8 ppm appear as two peaks likely due to $^{13}$C-$^{31}$P coupling (see additional zoomed spectra C and D in NMR characterization section at the end of the Supplementary Information document).

*[3] Signals between 29.4 and 31.8 ppm overlap, not all signals are resolved individually.

$^{31}$P NMR{$^{1}$H} (162 MHz, $CD_3OD$) δ = 0.12 ppm.

HR-MS (ESI negative) m/z calculated for $C_{39}H_{70}N_3O_8P$: 739.49005; found: 738.482 $[M-H]^-$.

Yield: 42.5 mg (57.4 µmol, 30 % over 2 steps).

### Synthesis of bifunctional Phosphatidylcholines

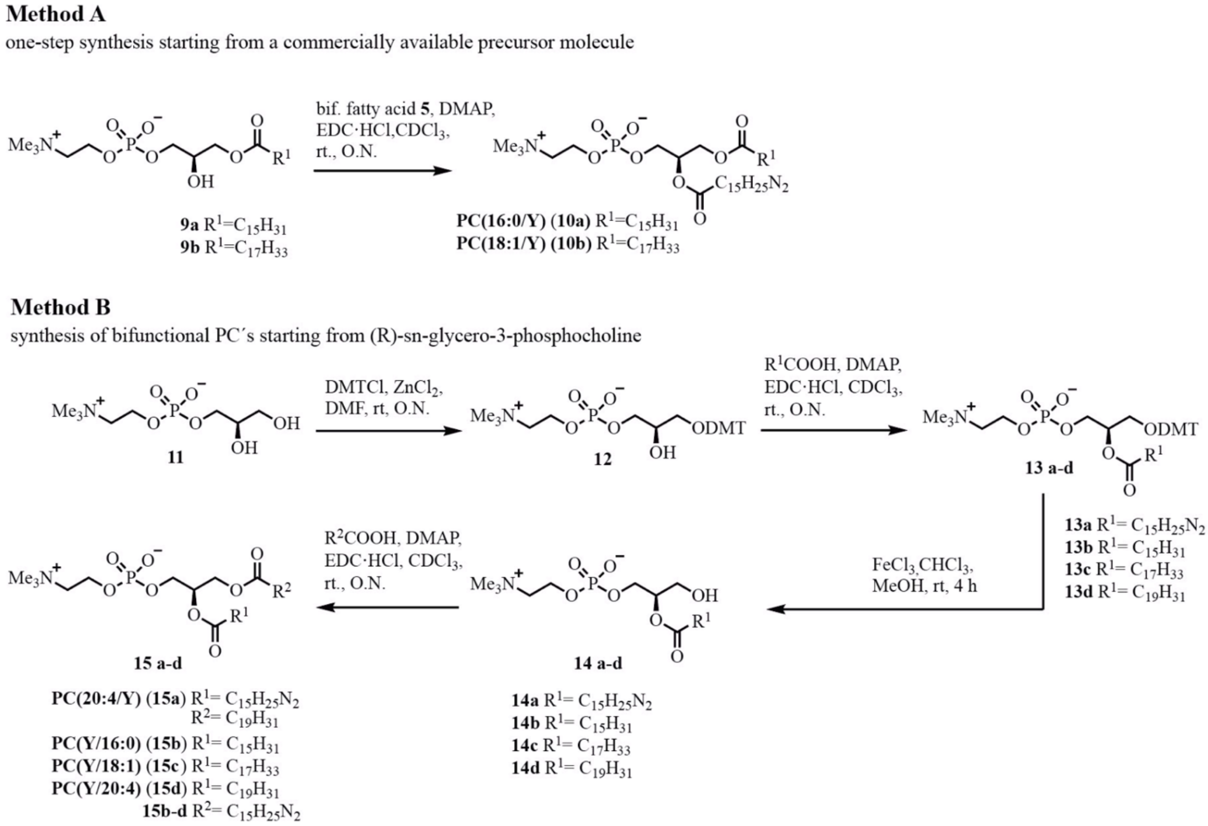

Scheme C: Synthesis of the bifunctional PCs.

##### *Synthesis of 1-DMT-(R)-sn-glycero-3-phosphocholine 12*

$ZnCl_2$ was dried for 4h under high vacuum prior to starting the reaction. 1.88 g (R)-*sn*-glycero-3-phosphocholine (**11**) (7.28 mmol, 1.0 eq.), 2.74 g DMTCl (8.01 mmol, 1.1 eq) and 1.0 g of $ZnCl_2$ (7.28 mmol, 1.0 eq.) were placed in a 100 ml flask under argon. 20 ml of dry DMF were added and the reaction mixture was stirred at room temperature for 20 h. The mixture was transferred into a separation funnel charged with sat. $NH_4HCO_3$ solution and extracted 3x with chloroform. The combined organic layers were dried over $Na_2SO_4$ and the solvent removed under reduced pressure. The crude product was purified by flash chromatography ($CHCl_3$/MeOH/$H_2O$/$NEt_3$, 70:30:5:0.2). DMT protected (R)-*sn*-glycero-3-phosphocholine **12** was isolated as a colourless solid in a yield of 41 % (1.66 g, 3.21 mmol).

##### *General procedure for the synthesis of 1-DMT-2-acyl-phosphaditylcholines 13 a-d*

A mixture of 2.7 eq. EDC·HCl, 0.5 eq. DMAP and 2.0 eq. of the respective fatty acid (bifunctional fatty acid, palmitic acid, oleic acid, arachidonic acid) was dissolved in 10 ml dry $CHCl_3$ after drying under high vacuum. A solution of 1.0 eq. 1-Dimethoxytrityl-(R)-*sn*-glycero-3-phosphocholine (**12**) in $CHCl_3$ (140 mM) was added after 10 min. The mixture was stirred over night at room temperature. The solvent was removed under reduced pressure and the crude products was purified by flash chromatography ($CHCl_3$/MeOH/$H_2O$, 70:30:5). 1-DMT-2-O-acyl-phosphaditylcholine **13 a-d** were isolated in yields of 62-75 %.

***General procedure for the synthesis of 2-Acyl-lysophosphatidylcholines 14 a-d***

1.1 eq. $FeCl_3$ were dissolved in a MeOH/chloroform (1:1) mixture (6 mM) and added to 1.0 eq. of 1-DMT-2-acyl-phosphaditylcholine **13 a-d**. The reaction mixture was stirred for 4h at room temperature. After removing the solvents under reduced pressure, the crude product was purified by flash chromatography ($CHCl_3$/MeOH/$H_2O$, 70:30:5). The deprotected 2-Acyl-lysophosphatidylcholines **14 a-d** were isolated in quantitative yields.

***General procedure for the synthesis of bifunctional phosphatidylcholines 15 a-d***

2.0 eq. of the respective fatty acid (bifunctional fatty acid, arachidonic acid) were dissolved in $CDCl_3$ (180 mM) and activated with 2.3 eq. EDC·HCl and 0.5 eq. DMAP. After 15 min, the solution of activated fatty acid was added to a mixture of 1.0 eq. of 2-Acyl-lysophosphatidylcholines **14 a-d** in $CDCl_3$ (50 mM). The reaction mixture was stirred overnight at room temperature. After removing the solvent under reduced pressure, the crude product was purified by flash chromatography ($CHCl_3$/MeOH/$H_2O$, 70:30:2). Bifunctional PCs **15 a-d** were isolated in yields of 10-50 %.

The analytical data of **12**, **13b** and **14b** were in accordance with those previously reported.[2] All other analytical data are listed below.

***PC(16:0/Y) (10a)***

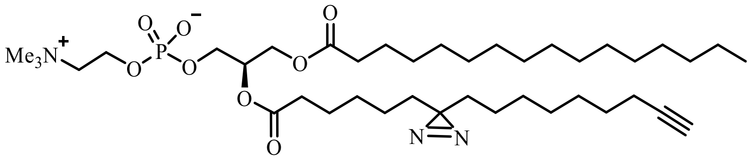

A solution of 160 mg of the bifunctional fatty acid **5** (575 µmol, 1.1 eq.) in 10 ml dry DMF was treated with 156 mg EDC·HCl (814 µmol, 1.6 eq.) and 31 mg DMAP (253.7 µmol, 0.5 eq.). After 10 min, the solution of the activated bifunctional fatty acid **5** was added to a solution of 253 mg 1-Palmitoyl-lyso-PC (**9a**) (510 µmol, 1.0 eq.) in 10 ml dry DMF. The reaction mixture was stirred overnight at room temperature. After removing the solvents under reduced pressure, the crude product was purified by HPLC using a Macherey Nagel VP 250/21 Nucleodur 100-5 C8ec Column at 12 ml/min eluting with a gradient. The solvent system used was A (MeOH/ water 3:1 + 4 % AcOH) and B (MeCN/MeOH/THF 4:3:1 + 4% AcOH). Gradient: 0-5 min: 0-45 % B, 5-15 min: 45-75 % B, 15-20 min: 75-100 % B, 20-26 min: 100 % B, 26-30 min: 100-0 % B. The retention time of the product was found to be 17.5 min. The product was isolated as a colourless oil.

$^{1}$H NMR (400 MHz, $CD_3OD$) δ = 5.24 ($m_c$, 1H), 4.44 (dd, $J$ = 12.0, 3.2 Hz, 1H), 4.32 – 4.22 (m, 2H), 4.17 (dd, $J$ = 12.0, 6.8 Hz, 1H), 4.00 ($m_c$, 2H), 3.68 – 3.58 (m, 2H), 3.23 (s, 9H), 2.33 (t, $J$ = 7.3 Hz, 2H), 2.32 (t, $J$ = 7.4 Hz, 2H), 2.20 – 2.09 (m, 3H), 1.67 – 1.43 (m, 6H), 1.43 – 1.20 (m, 36H), 1.15 – 1.02 (m, 4H), 0.90 (t, $J$ = 6.8 Hz, 3H) ppm.*[1]

*[1] Signals (triplets) at 2.33 and 2.32 ppm overlap.

$^{13}$C NMR{$^{1}$H} (101 MHz, $CD_3OD$) δ = 174.95, 174.45, 85.01, 71.90, 69.43, 67.52, 64.91, 63.62, 60.49, 54.68, 34.89, 33.83, 33.73, 33.10, 30.82, 30.78, 30.66, 30.50, 30.47, 30.22, 30.20, 29.98, 29.69, 29.64, 29.52, 26.02, 25.76, 24.88, 24.67, 23.76, 18.99, 14.46 ppm.*[2,3]

*[2] Signals at 71.9, 67.5, 64.9, 60.5 and 54.7 ppm appear as two peaks likely due to $^{13}$C-$^{31}$P coupling (see additional zoom spectrum E)

*[3] signals between 29.3 and 31.0 ppm overlap, not all signals are resolved individually.

HR-MS (ESI positive): calculated m/z for $C_{40}H_{74}N_3O_8P$: 755.52135, found: 756.528 $[M+H]^+$.

$^{31}$P NMR{$^{1}$H} (162 MHz, $CD_3OD$) δ = -0.57 ppm.

Yield: 98.0 mg (129.6 µmol, 25 %).

### ***PC(18:1/Y) (10b)***

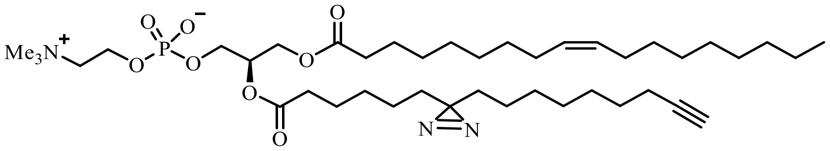

A solution of 156 mg of the bifunctional fatty acid **5** (560 µmol, 1.2 eq.) in 10 ml dry $CDCl_3$ was treated with 138 mg EDC·HCl (720 µmol, 1.5 eq.) and 33 mg DMAP (270 µmol, 0.6 eq.). After 10 min, the solution of the activated bifunctional fatty acid **5** was added to a solution of 191 mg 1-Oleoyl-lyso-PC (**9b**) (469 µmol, 1 eq.) in 10 ml dry DMF. The reaction mixture was stirred overnight at room temperature. After removing the solvent under reduced pressure, the crude product was purified by HPLC using a Macherey Nagel VP 250/21 Nucleodur 100-5 C8ec Column at 12 ml/min eluting with a gradient. The solvent system used was A (MeOH/ water 3:1 + 4 % AcOH) and B (MeCN/MeOH/*i*-propanol 5:1:4 + 4% AcOH). Gradient: 0-2 min: 0-45 % B, 2-8 min: 45-75 % B, 8-11 min: 75-100 % B, 11-26 min: 100 % B, 26-30 min: 100-0 % B. The retention time of the product was found to be 16.5 min. The product was isolated as a colourless oil.

$^{1}$H NMR (400 MHz, $CD_3OD$) δ = 5.35 ($m_c$, 2H), 5.24 ($m_c$, 1H), 4.44 (dd, J = 12.0, 3.2 Hz, 1H), 4.31 – 4.24 (m, 2H), 4.18 (dd, *J* = 12.0, 6.8 Hz, 1H), 4.00 ($m_c$, 2H), 3.68 – 3.62 (m, 2H), 3.23 (s, 9H), 2.33 (t, *J* = 7.2 Hz, 2H), 2.32 (t, *J* = 7.3 Hz, 2H), 2.19 – 2.12 (m, 3H), 2.09 – 2.00 (m, 4H), 1.66 – 1.53 (m, 4H), 1.52 – 1.45 (m, 2H), 1.45 – 1.21 (m, 32H), 1.16 – 1.04 (m, 4H), 0.91 (t, J = 6.8 Hz, 3H) ppm.*[1]

*[1] Signals (triplets) at 2.33 and 2.32 ppm overlap.

$^{13}$C NMR{$^{1}$H} (101 MHz, $CD_3OD$) δ = 174.90, 174.43, 130.94, 130.79, 85.02, 71.91, 71.83, 69.44, 67.48, 64.91, 64.86, 63.63, 60.49, 60.44, 54.73, 54.70, 54.66, 34.90, 33.84, 33.74, 33.08, 30.86, 30.63, 30.46, 30.36, 30.34, 30.23, 30.19, 29.97, 29.71, 29.69, 29.64, 29.63, 29.51, 28.16, 26.00, 25.76, 24.87, 24.66, 23.75, 19.00, 14.48 ppm.*[2,3]

*[2] Signals at 71.9, 67.5, 64.9, 60.5 and 54.7 ppm appear as two peaks likely due to $^{13}$C-$^{31}$P coupling (see additional zoomed spectrum F).

*[3] Signals between 29.4 and 31.0 ppm overlap, not all signals are resolved individually.

$^{31}$P NMR{$^{1}$H} (162 MHz, $CD_3OD$) δ = -0.57 ppm.

HR-MS (ESI positive): calculated m/z for $C_{42}H_{76}N_3O_8P$: 781.53700, found: 782.549 $[M+H]^+$.

Yield: 110 mg (141 µmol, 30 %).

***Bifunctional 1-DMT-2-acyl-phosphaditylcholine (13a)***

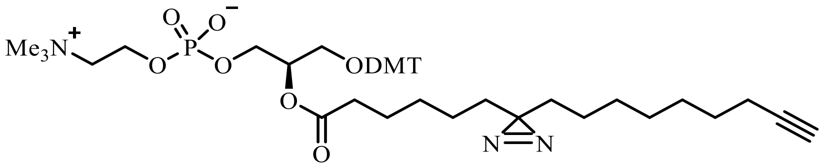

$^{1}$H NMR (400 MHz, $CD_3OD$) δ = 7.47 – 7.37 (m, 2H), 7.36 – 7.25 (m, 6H), 7.25 – 7.17 (m, 1H), 6.90 – 6.81 (m, 4H), 5.26 ($m_c$, 1H), 4.23 – 4.11 (m, 2H), 4.01 (t, $J$ = 5.7 Hz, 2H), 3.79 (s, 6H), 3.62 – 3.49 (m, 2H), 3.30 – 3.26 (m, 2H), 3.18 (s, 9H), 2.38 (t, $J$ = 7.3 Hz, 2H), 2.20 – 2.09 (m, 3H), 1.69 – 1.52 (m, 2H), 1.51 – 1.42 (m, 2H), 1.42 – 1.13 (m, 12H), 1.13 – 0.96 (m, 4H) ppm.

$^{13}$C NMR{$^{1}$H} (101 MHz, $CD_3OD$) δ = 174.75, 160.17, 146.26, 137.06, 131.40, 131.27, 129.38, 129.27, 128.83, 127.89, 114.15, 87.42, 85.07, 73.61, 69.42, 67.43, 65.70, 63.70, 60.40, 55.75, 54.67, 35.19, 33.80, 33.59, 30.12, 29.91, 29.74, 29.60, 29.52, 25.91, 24.79, 24.67, 18.97 ppm.*[1,2]

*[1] Signals at 73.6, 67.4, 65.7, 60.4, 54.7 ppm appear as two or more peaks because of the presence of two conformers (see additional zoomed spectrum G).

*[2] Signals between 29.4 ppm and 30.2 ppm overlap, not all signals resolved individually.

$^{31}$P NMR{$^{1}$H} (162 MHz, $CD_3OD$) δ = -0.56 ppm.

HR-MS (ESI positive): calculated m/z for $C_{45}H_{62}N_3O_9P$: 819.42237, found: 820.436 $[M+H]^+$.

Yield: 20.7 mg (25.2 µmol, 20 %).

##### *1-DMT-2-oleoyl-phosphaditylcholine (13c)*

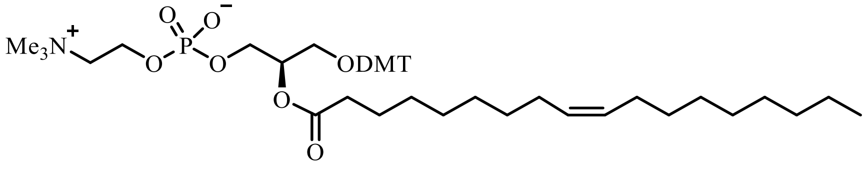

$^{1}$H NMR (400 MHz, $CD_3OD$) δ = 7.46 – 7.37 (m, 2H), 7.37 – 7.22 (m, 6H), 7.22 – 7.12 (m, 1H), 6.90 – 6.75 (m, 4H), 5.41 – 5.20 (m, 3H), 4.23 – 4.10 (m, 2H), 4.02 (t, $J$ = 5.7 Hz, 2H), 3.76 (s, 6H), 3.59 – 3.49 (m, 2H), 3.30 – 3.26 (m, 2H), 3.17 (s, 9H), 2.38 (t, $J$ = 7.3 Hz, 2H), 2.10 – 1.90 (m, 4H), 1.70 – 1.55 (m, 2H), 1.40 – 1.16 (m, 20H), 0.88 (t, $J$ = 6.8 Hz, 3H) ppm.*[1]

$^{13}$C NMR{$^{1}$H} (101 MHz, $CD_3OD$) δ = 174.49, 160.04, 146.22, 137.00, 136.96, 131.20, 130.00, 129.94, 129.43, 129.19, 129.15, 128.89, 128.81, 128.74, 127.85, 114.14, 87.36, 73.54, 73.46, 67.34, 65.62, 65.57, 63.65, 60.34, 60.29, 55.73, 54.63, 34.67, 32.59, 30.41, 28.16, 27.60, 26.57, 26.54, 25.98, 23.59, 14.51 ppm.*[2]

*[1] Signals at 73.5, 67.3, 65.6, 60.4, 54.6 ppm appear as two likely due to $^{13}$C-$^{31}$P coupling (see additional zoomed spectrum H).

*[2] Signals between 30.1 and 30.9 ppm overlap, not all signals are resolved individually.

$^{31}$P NMR{$^{1}$H} (162 MHz, $CD_3OD$) δ = -0.57 ppm.

HR-MS (ESI positive): calculated m/z for $C_{47}H_{70}NO_9P$: 823.47882, found: 824.493 $[M+H]^+$.

Yield: 226 mg (274 µmol, 71 %).

##### *1-DMT-2-arachidonyl PC (13d)*

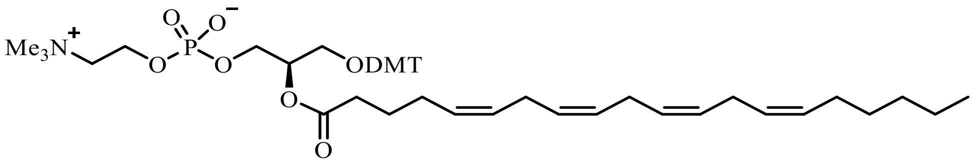

$^{1}H$ NMR (400 MHz, $CD_3OD$) δ = δ 7.44 – 7.32 (m, 2H), 7.32 – 7.19 (m, 6H), 7.19 – 7.05 (m, 1H), 6.88 – 6.68 (m, 4H), 5.40 – 5.17 (m, 9H), 4.22 – 4.06 (m, 2H), 4.01 (t, $J$ = 5.6 Hz, 2H), 3.70 (s, 6H), 3.57 – 3.44 (m, 2H), 3.29 – 3.20 (m, 2H), 3.13 (s, 9H), 2.76 ($m_c$, 6H), 2.36 (t, $J$ = 7.6 Hz, 2H), 2.14 – 1.92 (m, 4H), 1.67 ($m_c$, 2H), 1.36 – 1.16 (m, 6H), 0.84 (t, $J$ = 6.8 Hz, 3H) ppm.*[1]

*[1] The integrals of double bond protons and neighbouring methylene groups (m, 5.4-5.17 ppm and $m_c$, 2.76) were found to be lower than expected, possibly due to H-D exchange. However, the product was confirmed by MS and after DMT deprotection in the next step the expected integrals for the same protons were again observed.

$^{13}C$ NMR{$^{1}H$} (101 MHz, $CD_3OD$) δ = 174.49, 160.04, 146.22, 137.04, 137.00, 136.96, 131.33, 131.20, 130.00, 129.94, 129.43, 129.19, 129.15, 128.89, 128.81, 128.74, 127.85, 114.14, 87.36, 73.54, 67.34, 65.62, 63.65, 60.34, 55.73, 54.63, 34.67, 32.59, 30.41, 28.16, 27.60, 26.57, 26.54, 25.98, 23.59, 14.51 ppm.*[2,3]

*[2] signals at 73.5, 67.3, 65.6, 60.3, 54.6 ppm appear as two likely due to $^{13}C$-$^{31}P$ coupling (see additional zoomed spectrum I).

*[3] signals between 26.5 and 26.7 ppm overlap, not all signals resolved individually.

$^{31}P$ NMR{$^{1}H$} (162 MHz, $CD_3OD$) δ = -0.57 ppm.

HR-MS (ESI positive): calculated m/z for $C_{49}H_{68}NO_9P$: 845.463, found: 846.670 $[M+H]^+$.

Yield: 203 mg (240 µmol, 62 %).

##### *sn2-BIF-sn-lysophosphocholine (14a)*

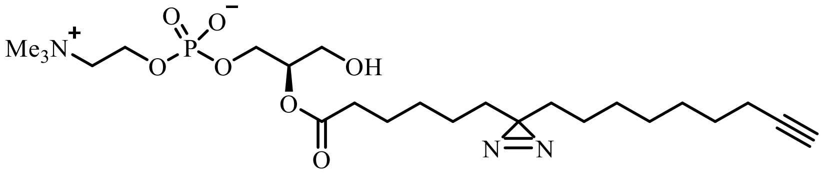

$^{1}$H NMR (400 MHz, $CD_3OD$) δ = δ 5.00 ($m_c$, 1H), 4.28 ($m_c$, 2H), 4.00 ($m_c$, 2H), 3.74 – 3.64 (m, 4H), 3.24 (s, 9H), 2.35 (t, $J$ = 7.5 Hz, 2H), 2.22 – 2.11 (m, 3H), 1.65 – 1.44 (m, 4H), 1.44 – 1.22 (m, 12H), 1.15 – 1.04 (m, 4H) ppm.

$^{13}$C NMR{$^{1}$H} (101 MHz, $CD_3OD$) δ = 174.90, 85.06, 74.76, 69.44, 67.51, 64.78, 61.22, 60.50, 54.73, 49.85, 49.64, 49.43, 49.21, 49.00, 48.79, 48.57, 48.36, 34.92, 34.73, 33.77, 33.67, 30.14, 29.92, 29.70, 29.58, 25.71, 24.81, 24.62, 18.96 ppm.*[1,2]

*[1] Signals at 74.8, 67.5, 64.8, 60.5, 54.7 ppm appear as two peaks likely due to $^{13}$C-$^{31}$P coupling (see additional zoomed spectrum J).

*[2] Signals between 29.4 and 30.2 ppm overlap, not all signals are resolved individually.

$^{31}$P NMR{$^{1}$H} (162 MHz, $CD_3OD$) δ = -0.81 ppm.

HR-MS (ESI positive): calculated m/z for $C_{24}H_{44}N_3O_7P$: 517.29169, found: 518.303 $[M+H]^+$.

Yield: 13.0 mg (25.1 µmol, quant.).

***sn2-Oleoyl-sn-lysophosphocholine (14c)***

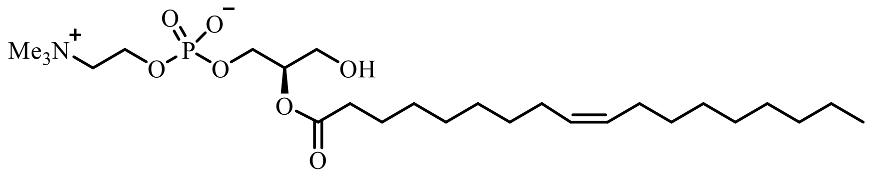

$^1$H NMR (400 MHz, $CD_3OD$) δ = 5.34 ($m_c$, 2H), 5.01 ($m_c$, 1H), 4.31 ($m_c$, 2H), 4.02 ($m_c$, 2H), 3.76 – 3.64 (m, 4H), 3.26 (s, 9H), 2.36 (t, $J$ = 7.3 Hz, 2H), 2.09 – 1.98 (m, 4H), 1.62 ($m_c$, 2H), 1.42 – 1.23 (m, 20H), 0.90 (t, $J$ = 6.8 Hz, 3H) ppm.

$^{13}$C NMR{$^1$H} (101 MHz, $CD_3OD$) δ = 174.95, 130.87, 79.51, 74.67, 67.45, 64.80, 61.18, 60.59, 54.73, 35.09, 33.03, 30.82, 30.58, 30.41, 30.34, 30.31, 30.20, 28.12, 28.11, 25.96, 23.71, 14.46 ppm.[1,2]

*[1] Signals at 74.7, 67.5, 64.8, 60.6 and 54.7 ppm appear as two peaks likely due to $^{13}$C-$^{31}$P coupling (see additional zoomed spectrum K).

*[2] Signals between 30.1 and 30.9 ppm overlap, not all signals are resolved individually.

$^{31}$P NMR{$^1$H} (162 MHz, $CD_3OD$) δ = -1.44 ppm.

HR-MS (ESI positive): calculated m/z for $C_{26}H_{52}NO_7P$: 521.34814, found: 522.358 $[M+H]^+$.

Yield: 56.6 mg (109 µmol, quant.).

***sn2-Arachidonyl-sn-lysophosphocholine (14d)***

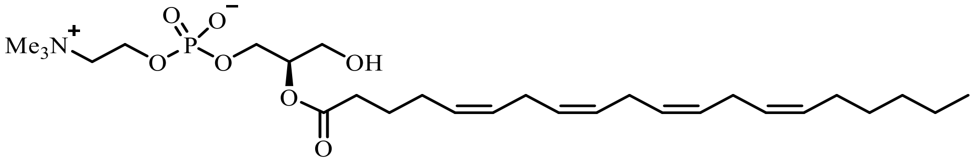

Compound is not stable enough for storage and needs to be used immediately for the next step after purification.

$^{1}$H NMR (400 MHz, $CD_3OD$) δ = 5.48 – 5.24 (m, 8H), 5.01 ($m_c$, 1H), 4.33 – 4.23 (m, 2H), 4.00 ($m_c$, 2H), 3.74 – 3.62 (m, 4H), 3.24 (s, 9H), 2.90 – 2.75 (m, 6H), 2.38 (t, $J$ = 7.5 Hz, 2H), 2.19 – 2.03 (m, 4H), 1.69 ($m_c$, 2H), 1.42 – 1.24 (m, 6H), 0.91 (t, $J$ = 6.8 Hz, 3H) ppm.

$^{13}$C NMR{$^{1}$H} (101 MHz, $CD_3OD$) δ = 174.70, 131.19, 130.03, 129.88, 129.45, 129.17, 128.88, 128.75, 79.49, 74.73, 67.43, 64.73, 61.18, 60.48, 54.69, 34.51, 32.63, 30.44, 28.17, 27.53, 26.54, 25.89, 23.61, 14.46 ppm.*[1,2]

*[1] Signals at 74.7, 67.4, 64.7, 60.5 and 54.7 ppm appear as two peaks likely due to $^{13}$C-$^{31}$P coupling (see additional zoomed spectrum L).

*[2] Signals between 26.4 and 26.7 ppm overlap, not all signals are resolved individually.

$^{31}$P NMR{$^{1}$H} (162 MHz, $CD_3OD$) δ = -0.40 ppm.

HR-MS (ESI positive): calculated m/z for $C_{28}H_{50}NO_7P$: 543.332, found: 544.339 $[M+H]^+$.

Yield: 26.4 mg (48.6 µmol, 63 %).

***PC(20:4/Y) (15a)***

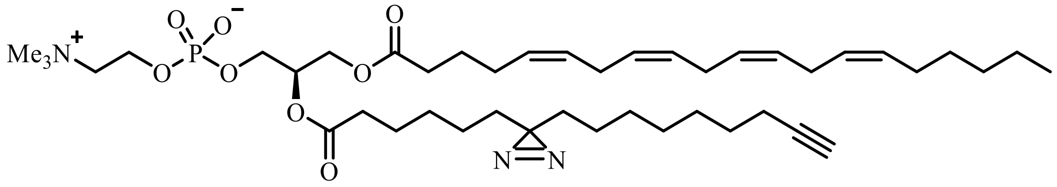

$^1$H NMR (400 MHz, $CD_3OD$) δ = 5.45 – 5.29 (m, 8H), 5.24 ($m_c$, 1H), 4.43 (dd, $J$ = 12.0, 3.2 Hz, 1H), 4.32 – 4.22 (m, 2H), 4.18 (dd, $J$ = 12.0, 6.8 Hz, 1H), 4.00 (t, $J$ = 5.8 Hz, 2H), 3.67 – 3.59 (m, 2H), 3.23 (s, 9H), 2.91 – 2.79 (m, 6H), 2.38 – 2.28 (m, 4H), 2.20 – 2.04 (m, 7H), 1.75 – 1.63 (m, 2H), 1.62 – 1.43 (m, 4H), 1.43 – 1.22 (m, 18H), 1.15 – 1.02 (m, 4H), 0.91 (t, $J$ = 6.9 Hz, 3H) ppm.

$^{13}$C NMR{$^1$H} (101 MHz, $CD_3OD$) δ = 174.72, 174.46, 131.23, 130.01, 129.95, 129.49, 129.21, 129.17, 128.91, 128.79, 85.03, 71.89, 69.45, 67.47, 64.91, 63.70, 60.49, 54.69, 34.89, 34.28, 33.82, 33.73, 32.68, 30.49, 30.19, 29.97, 29.68, 29.63, 29.53, 28.22, 27.54, 26.60, 25.88, 25.76, 24.87, 24.66, 23.65, 18.99, 14.48 ppm.*[2,3]

*[1] Signals at 71.9, 67.5, 64.9, 60.5 and 54.7 ppm appear as two peaks likely due to $^{13}$C-$^{31}$P coupling (see additional zoomed spectrum M).

*[2] Signals between 29.4 and 29.8 ppm and 26.4 and 26.8 ppm overlap, not all signals are resolved individually.

$^{31}$P NMR{$^1$H} (162 MHz, $CD_3OD$) δ = -0.57 ppm.

HR-MS (ESI positive): calculated m/z for $C_{44}H_{74}N_3O_8P$: 803.52135, found: 804.535 $[M+H]^+$.

Yield: 8.2 mg (10 μmol, 41 %).

### *PC(Y/16:0) (15b)*

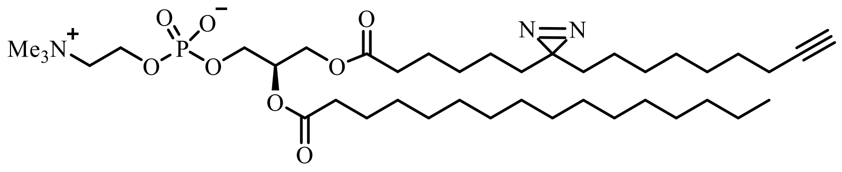

[1]H NMR (400 MHz, $CD_3OD$) δ = 5.24 ($m_c$, 1H), 4.43 (dd, *J* = 12.0, 3.1 Hz, 1H), 4.28 ($m_c$, 2H), 4.17 (dd, *J* = 12.1, 7.0 Hz, 1H), 4.00 (t, *J* = 5.9 Hz, 2H), 3.68 – 3.61 (m, 2H), 3.23 (s, 9H), 2.35 (t, *J* = 7.6 Hz, 2H), 2.31 (t, *J* = 7.6 Hz, 2H), 2.20 – 2.11 (m, 3H), 1.66 – 1.43 (m, 6H), 1.43 – 1.23 (m, 36H), 1.17 – 1.03 (m, 4H), 0.90 (t, *J* = 6.8 Hz, 3H) ppm.*[1]

*[1] Signals (triplets) at 2.35 and 2.31 ppm overlap.

[13]C NMR{[1]H} (101 MHz, $CD_3OD$) δ = 174.77, 174.65, 85.02, 71.83, 69.43, 67.48, 64.92, 63.72, 60.50, 54.70, 35.09, 34.72, 33.82, 33.73, 33.09, 30.81, 30.78, 30.67, 30.49, 30.19, 29.97, 29.71, 29.64, 29.63, 29.50, 26.05, 25.73, 24.87, 24.64, 23.75, 18.99, 14.45 ppm.*[2,3]

*[2] Signals at 71.8, 67.5, 64.9, 60.5 and 54.7 ppm appear as two peaks likely due to $^{13}C$-$^{31}P$ coupling (see additional zoomed spectrum N).

*[3] Signals between 29.4 and 30.9 ppm overlap, not all signals are resolved individually.

[31]P NMR{[1]H} (162 MHz, $CD_3OD$) δ = -0.57 ppm.

HR-MS (ESI positive): calculated m/z for $C_{40}H_{74}N_3O_8P$: 755.52135, found: 756.535 $[M+H]^+$.

Yield: 16.7 mg (22.1 µmol, 55 %).

***PC(Y/18:1) (15c)***

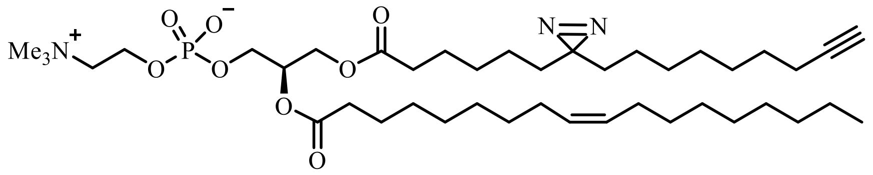

$^{1}$H NMR (400 MHz, $CD_3OD$) δ = 5.35 ($m_c$, 2H), 5.24 ($m_c$, 1H), 4.44 (dd, *J* = 12.0, 2.8 Hz, 1H), 4.34 – 4.22 (m, 2H), 4.17 (dd, *J* = 12.0, 6.9 Hz, 1H), 4.00 (t, *J* = 5.9 Hz, 2H), 3.70 – 3.60 (m, 2H), 3.23 (s, 9H), 2.35 (t, *J* = 7.6 Hz, 2H), 2.31 (t, *J* = 7.6 Hz, 2H), 2.22 – 2.11 (m, 3H), 2.11 – 1.96 (m, 4H), 1.69 – 1.44 (m, 6H), 1.44 – 1.19 (m, 32H), 1.15 – 1.03 (m, 4H), 0.91 (t, *J* = 6.4 Hz, 3H) ppm.*[1]

*[1] Signals (triplets) at 2.35 and 2.31 ppm overlap.

$^{13}$C NMR{$^{1}$H} (101 MHz, $CD_3OD$) δ = 174.80, 174.65, 130.94, 130.79, 85.04, 71.83, 69.44, 67.48, 64.91, 63.73, 60.49, 54.71, 49.85, 49.64, 49.43, 49.28, 49.21, 49.00, 48.79, 48.58, 48.36, 35.08, 34.72, 33.81, 33.72, 33.07, 30.84, 30.61, 30.45, 30.35, 30.24, 30.17, 29.95, 29.70, 29.61, 29.51, 28.15, 26.03, 25.73, 24.86, 24.63, 23.74, 18.98, 14.47 ppm.*[2,3]

*[2] Signals at 71.8, 67.5, 64.9, 60.5 and 54.7 ppm appear as multiplets likely due to $^{13}$C-$^{31}$P coupling (see additional zoomed spectrum O).

*[3] Signals between 29.5 and 30.9 ppm overlap and are not resolved individually.

$^{31}$P NMR{$^{1}$H} (162 MHz, $CD_3OD$) δ = -0.57 ppm.

HR-MS (ESI positive): calculated m/z for $C_{42}H_{76}N_3O_8P$: 781.53700, found: 782.550 $[M+H]^+$.

Yield: 26.0 mg (33.3 mmol, 51 %).

***PC(Y/20:4) (15d)***

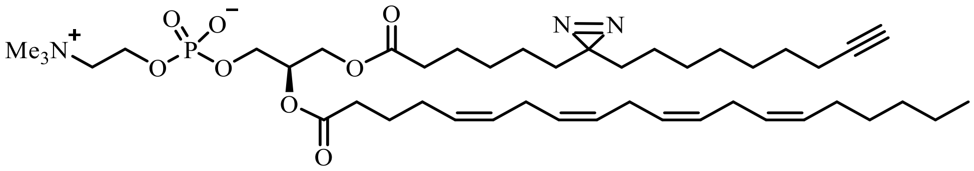

Coupling reaction needs to be performed at 0 °C.

$^{1}$H NMR (400 MHz, $CD_3OD$) δ = 5.49 – 5.30 (m, 8H), 5.25 ($m_c$, 1H), 4.44 (dd, $J$ = 12.0, 3.2 Hz, 1H), 4.28 ($m_c$, 2H), 4.17 (dd, $J$ = 12.0, 6.8 Hz, 1H), 4.00 (t, $J$ = 5.8 Hz, 2H), 3.69 – 3.59 (m, 2H), 3.23 (s, 9H), 2.91 – 2.78 (m, 6H), 2.37 (t, $J$ = 7.4 Hz, 2H), 2.31 (t, $J$ = 7.4 Hz, 2H), 2.21 – 1.97 (m, 7H), 1.78 – 1.64 (m, 2H), 1.64 – 1.43 (m, 4H), 1.43 – 1.20 (m, 18H), 1.18 – 1.01 (m, 4H), 0.94 – 0.88 (m, 3H) ppm.*[1]

*[1] Signals (triplets) at 2.37 and 2.31 ppm overlap.

$^{13}$C NMR{$^{1}$H} (101 MHz, $CD_3OD$) δ = 174.72, 174.38, 131.23, 130.01, 129.98, 129.50, 129.24, 129.18, 128.92, 128.78, 85.03, 71.88, 69.47, 67.46, 64.89, 63.69, 60.49, 54.68, 34.70, 34.47, 33.82, 33.73, 32.69, 30.49, 30.19, 29.97, 29.71, 29.63, 29.52, 28.22, 27.53, 26.60, 25.94, 25.88, 25.76, 25.73, 24.88, 24.65, 23.66, 19.00, 14.49 ppm.*[2,3]

*[2] Signals at 71.9, 67.5, 64.9, 60.5 and 54.7 ppm appear as two peaks likely due to $^{13}$C-$^{31}$P coupling (see additional zoomed spectrum N).

*[3] Signals between 29.5 and 30.9 ppm overlap, not all signals are resolved individually.

$^{31}$P NMR{$^{1}$H} (162 MHz, $CD_3OD$) δ = -0.56 ppm.

HR-MS (ESI positive): calculated m/z for $C_{44}H_{74}N_3O_8P$: 803.52135, found: 804.536 $[M+H]^+$.

Yield: 20.0 mg (24.9 µmol, 21 %).

### Synthesis of Sphingomyelin(Y) (16)

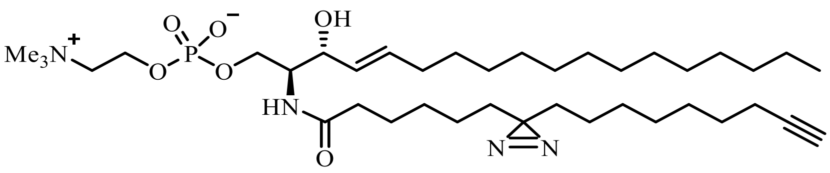

16.5 mg of the bifunctional fatty acid **5** (59.3 µmol, 1.2 eq.), 28.2 mg HBTU (74.4 µmol, 1.5 eq.) and 1.4 mg of HOBt (10.4 µmol, 0.2 eq.) were dissolved in 4 ml $CDCl_3$. After adding 16 µl of DIPEA (91.9 µmol, 1.9 eq.) the reaction mixture was stirred for 10 min at 40 °C. Subsequently a solution of 23 mg Lyso-Sphingomyelin (49.5 µmol, 1.0 eq.) in 6 ml $CDCl_3$ was added and the mixture was stirred for 4 h at 40 °C, allowed to reach RT and stirred overnight. After removing the solvent under reduced pressure, the crude product was purified by HPLC using a Macherey Nagel VP 250/21 Nucleodur C18 Pyramid column (10µm, 250mm) at 10 ml/min eluting with a gradient. The solvent system used was A (50 % MeOH, 50 % water) and B (62 % *i*-propanol, 19% MeCN, 19 % THF). Gradient: 0-18 min: 100-80 % B, 18-26 min: 80-100 % B, 26-45 min: 100-20% B, 45-50 min: 20-0 % B. The retention time of the product was found to be 38 min. Bifunctional sphingomyelin **16** was isolated as a colourless oil.

$^{1}H$ NMR (400 MHz, $CD_3OD$) δ = 5.70 (dt, $J_1$ = 15.3 Hz, $J_2$ = 6.8 Hz, 1H), 5.45 (dd, $J$ = 15.3, 7.7 Hz, 1H), 4.28 (m, 2H), 4.15 – 4.00 (m, 2H), 4.00 – 3.87 (m, 2H), 3.63 (t, $J$ = 4.5 Hz, 2H), 3.22 (s, 9H), 2.23 – 2.08 (m, 5H), 2.03 (td, $J_1$ = 7.0 Hz, $J_2$ = 7.0 Hz, 2H), 1.65 – 1.44 (m, 4H), 1.44 – 1.18 (m, 36H), 1.18 – 1.01 (m, 4H), 0.90 (t, $J$ = 6.8 Hz, 3H) ppm.

$^{13}C$ NMR{$^{1}H$} (101 MHz, $CD_3OD$) δ = 175.70, 135.10, 131.22, 85.00, 72.59, 69.42, 67.47, 65.83, 60.44, 55.31, 54.69, 37.13, 33.86, 33.78, 33.46, 33.10, 30.84, 30.80, 30.75, 30.50, 30.42, 30.21, 29.99, 29.64, 29.52, 26.88, 24.89, 24.77, 23.75, 18.99, 14.46 ppm.*[1,2]

*[1] Signals at 67.5 65.8, 60.4, 55.3 and 54.7 ppm appear as two peaks likely due to $^{13}C$-$^{31}P$ coupling (see additional zoom picture Q).

*[2] signals between 29.4 and 31.0 ppm overlap, not all signals are resolved individually.

$^{31}P$ NMR{$^{1}H$} (162 MHz, $CD_3OD$) δ = 0.03 ppm.

HR-MS (ESI positive): calculated m/z for $C_{39}H_{73}N_4O_6P$: 724.52677, found: 725.531 $[M+H]^+$.

Yield: 5.0 mg (6.9 µmol, 14 %).

### Synthesis of Phosphatidic acid (18:1/Y)

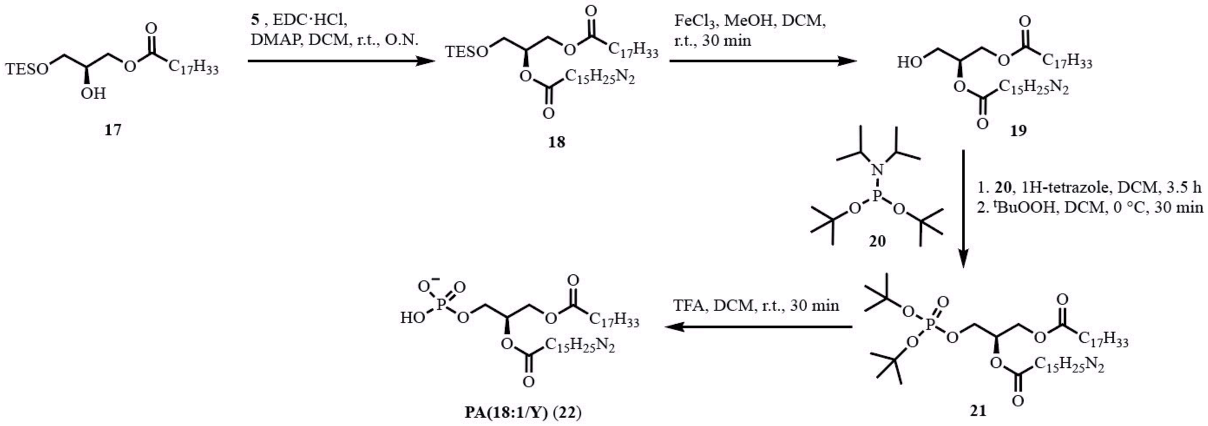

The synthesis of 17 was performed according to [3].

***(R)-2-((6-(3-(non-8-yn-1-yl)-3H-diazirin-3-yl)hexanoyl)oxy)-3-((triethyl-silyl)oxy)propyl oleate (18)***

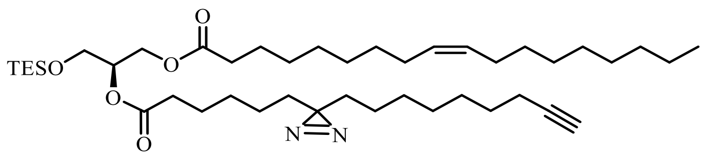

A solution of 50.0 mg of the bifunctional fatty acid **5** (409 µmol, 1.6 eq.), 80.0 mg EDC·HCl (417 µmol, 1.6 eq.) and 15.0 mg DMAP (53.9 µmol, 0.2 eq.) in 3 ml dry DCM was stirred for 10 min under an inert atmosphere. Subsequently, a solution of 120 mg of **17** (255 µmol, 1 eq.) in 5 ml dry DCM was added and the reaction mixture was stirred overnight. The solvent was removed under reduced pressure and the residue was purified by flash chromatography (CyHex/EtOAc, 9:1). The TES protected DAG **18** was isolated as a colourless oil.

$^{1}$H NMR (400 MHz, $CDCl_3$) δ = 5.34 ($m_c$, 2H), 5.06 ($m_c$, 1H), 4.35 (dd, $J$ = 11.9, 3.7 Hz, 1H), 4.16 (dd, $J$ = 11.8, 6.2 Hz, 1H), 3.71 ($m_c$, 2H), 2.29 (t, $J$ = 7.5 Hz, 2H), 2.28 (t, $J$ = 7.4 Hz, 2H), 2.17 (td, $J$ = 7.1, 2.6 Hz, 2H), 2.08 – 1.95 (m, 4H), 1.93 (t, $J$ = 2.6 Hz, 1H), 1.67 – 1.44 (m, 6H), 1.44 – 1.14 (m, 32H), 1.14 – 1.01 (m, 4H), 0.94 (t, $J$ = 8.3 Hz, 9H), 0.88 (t, $J$ = 6.7 Hz, 3H), 0.59 (q, $J$ = 7.9 Hz, 6H) ppm.*[1]

*[1] Signals (triplets) at 2.29 ppm and 2.28 ppm overlap.

$^{13}$C NMR{$^{1}$H} (101 MHz, $CDCl_3$) δ = 173.55, 172.99, 130.17, 129.88, 84.78, 72.02, 68.27, 62.57, 61.37, 34.30, 34.25, 33.04, 32.90, 32.06, 29.92, 29.88, 29.68, 29.48, 29.46, 29.35, 29.27, 29.21, 29.00, 28.83, 28.79, 28.70, 28.54, 27.38, 27.34, 25.06, 24.81, 23.93, 23.70, 22.83, 18.51, 14.24, 6.79, 4.47 ppm.*[2]

*[2] Signals between 28.5 and 30.0 ppm overlap, not all signals are resolved individually.

HR-MS (ESI positive): calculated m/z for $C_{43}H_{78}N_2O_5Si$: 730.56800, found: 748.602 $[M+NH_4]^+$.

Yield: 80.0 mg (109 µmol, 43 %).

***(S)-3-hydroxy-2-((6-(3-(non-8-yn-1-yl)-3H-diazirin-3-yl)hexanoyl)oxy) propyl oleate (19)***

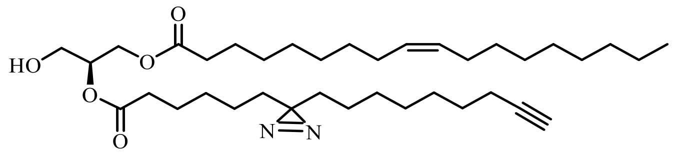

A solution of 55.0 mg of **18** (67.7 µmol, 1 eq.) in 10 ml 5 mM $FeCl_3$ solution (50 µmol, 0.7 eq.) in MeOH/DCM 3:1 was stirred for 30 min at room temperature. The reaction mixture was transferred onto a mixture of EtOAc and $H_2O$. The layers were separated and the organic layer was washed with sat. NaCl solution and dried over $Na_2SO_4$. After removing the solvents under reduced pressure, the residue was purified by flash chromatography (CyHex/EtOAc, 4:1) to yield 35.0 mg of pure product **19**.

$^1$H NMR (400 MHz, $CDCl_3$) δ = 5.34 ($m_c$, 2H), 5.07 (dddd, $J_{1-4}$ = 5.0 Hz, 1H), 4.32 (dd, $J$ = 11.9, 4.6 Hz, 1H), 4.23 (dd, $J$ = 11.9, 5.6 Hz, 1H), 3.76 – 3.66 (m, 2H), 2.33 (t, $J$ = 7.5 Hz, 2H), 2.32 (t, $J$ = 7.5 Hz, 2H), 2.17 (td, $J$ = 7.1, 2.6 Hz, 2H), 2.09 – 1.95 (m, 4H), 1.94 (t, $J$ = 2.6 Hz, 1H), 1.67 – 1.55 (m, 4H), 1.55 – 1.44 (m, 2H), 1.44 – 1.16 (m, 32H), 1.16 – 1.00 (m, 4H), 0.88 (t, $J$ = 6.8 Hz, 3H) ppm.*[1]

*[1] Signals (triplets) at 2.33 ppm and 2.32 ppm overlap.

$^{13}$C NMR{$^1$H} (101 MHz, $CDCl_3$) δ = 173.90, 173.28, 130.19, 129.86, 84.80, 72.34, 68.28, 62.08, 61.66, 34.22, 34.18, 33.02, 32.81, 32.05, 29.91, 29.85, 29.67, 29.46, 29.32, 29.24, 29.20, 28.99, 28.87, 28.73, 28.69, 28.52, 27.37, 27.32, 25.02, 24.79, 23.92, 23.67, 22.83, 18.50, 14.26 ppm.*[2]

*[2] Signals between 28.5 and 30.0 ppm overlap, not all signals are resolved individually.

HR-MS (ESI positive): calculated m/z for $C_{37}H_{64}N_2O_5$: 616.48152, found: 634.521 $[M+NH_4]^+$.

Yield: 35.0 mg (56.7 µmol, 84 %).

***(R)-3-((di-tert-butoxyphosphoryl)oxy)-2-((6-(3-(non-8-yn-1-yl)-3H-diazirin-3-yl)hexanoyl)oxy)propyl oleate (21)***

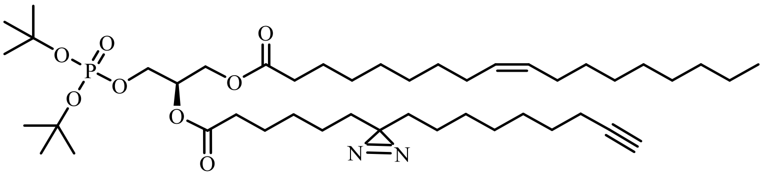

Molecular sieves (3Å, powder) were placed in a round-bottom flask and activated with heat under high vacuum. A solution of 35.0 mg of **19** (56.7 µmol, 1.0 eq.) in 4 ml dry DCM was added under inert conditions. After adding 50 µl of Di-tert-butyl *N,N*-diisopropyl-phosphoramidite (**20**) (159 µmol, 2.8 eq.), the reaction mixture was stirred for 30 min at room temperature. A solution of 1H-tetrazole (450 µl, 0.45 M in MeCN, 202 µmol, 3.6 eq.) was added and the reaction mixture was stirred for 3.5 h at room temperature. Subsequently, 40 µl of $^{t}$BuOOH (5.5 M in decane, 221 µmol, 3.9 eq.) was added dropwise at 0 °C and the reaction mixture was stirred for 30 min at 0 °C. The reaction was quenched with sat. $Na_2S_2O_3$ solution. Molecular sieves were removed by filtration. The reaction mixture was extracted 3x with DCM, the combined organic layers dried over $Na_2SO_4$ and the solvent removed under reduced pressure. The crude product was purified by flash chromatography (CyHex/EtOAc, 9:1). The product **21** was isolated as a light yellowish oil.

$^1$H NMR (400 MHz, $CD_3OD$) δ = 5.35 (m$_c$, 2H), 5.25 (m$_c$, 1H), 4.40 (dd, *J* = 12.0, 3.8 Hz, 1H), 4.17 (dd, *J* = 12.0, 6.2 Hz, 1H), 4.14 – 4.04 (m, 2H), 2.34 (2xt, *J* = 7.3 Hz, 4H), 2.21 – 2.11 (m, 3H), 2.12 – 1.96 (m, 4H), 1.69 – 1.54 (m, 4H), 1.54 – 1.44 (m, 20H), 1.44 – 1.18 (m, 32H), 1.19 – 1.01 (m, 4H), 0.93 – 0.85 (m, 3H) ppm. [*1]

$^{13}$C NMR{$^1$H} (101 MHz, $CD_3OD$) δ = 174.69, 174.09, 130.94, 130.79, 85.01, 84.91, 84.83, 84.77, 71.22, 69.44, 66.13, 63.05, 34.87, 33.85, 33.73, 33.08, 30.87, 30.82, 30.63, 30.48, 30.38, 30.31, 30.20, 30.16, 29.98, 29.65, 29.48, 28.17, 28.15, 26.01, 25.72, 24.88, 24.63, 23.76, 19.01, 14.50 ppm.[*2]

[*1]The two triplets from the $CH_2$-groups next to the carboxyl ester overlap.

[*2] Signals at 71.2 and 66.1 ppm appear as two peaks likely due to $^{13}$C-$^{31}$P coupling (see additional zoomed spectrum R).

$^{31}$P NMR{$^1$H} (162 MHz, $CD_3OD$) δ = -10.61 ppm.

HR-MS (ESI positive): calculated m/z for $C_{45}H_{81}N_2O_8P$: 808.57305, found: 826.614 $[M+NH_4]^+$.

Yield: 24.0 mg (29.7 µmol, 52 %).

***PA(18:1/Y) (22)***

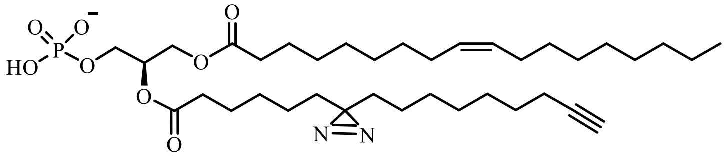

A solution of 24.0 mg of **21** (29.7 µmol, 1.0 eq.) in 1.5 ml dry DCM was treated with 40 µl of TFA (519 µmol, 17 eq.) and the reaction mixture was stirred at room temperature for 30 min. Subsequently, all volatiles were removed under high vacuum. As the $^{1}$HNMR spectra indicated remaining starting material, the protocol was repeated (1.5 ml dry DCM, 40 µl TFA, 30 min r.t.). After removing all volatiles under high vacuum, the product was isolated in quantitative yield and high purity as a light yellowish oil.

$^{1}$H NMR (400 MHz, $CD_3OD$) δ = 5.35 ($m_c$, 2H), 5.23 ($m_c$, 1H), 4.40 (dd, $J$ = 12.0, 3.4 Hz, 1H), 4.18 (dd, $J$ = 12.0, 6.4 Hz, 1H), 4.14 – 4.01 (m, 2H), 2.34 (t, $J$ = 7.1 Hz, 4H), 2.33 (t, $J$ = 7.3 Hz, 4H), 2.24 – 2.11 (m, 3H), 2.11 – 1.88 (m, 4H), 1.73 – 1.44 (m, 6H), 1.45 – 1.23 (m, 32H), 1.23 – 0.98 (m, 4H), 0.91 (t, $J$ = 6.6 Hz, 4H) ppm.[*1]

*[1] Triplets at 2.34 and 2.33 ppm overlap.

$^{13}$C NMR{$^{1}$H} (101 MHz, $CD_3OD$) δ = 174.82, 174.30, 130.93, 130.79, 85.01, 71.46, 69.41, 65.52, 63.19, 34.86, 33.83, 33.74, 33.08, 30.86, 30.83, 30.63, 30.47, 30.37, 30.32, 30.21, 30.19, 29.97, 29.67, 29.64, 29.49, 28.16, 25.99, 25.72, 24.87, 24.64, 23.75, 19.00, 14.48 ppm.[*2, 3]

[*2] signals at 71.5 and 65.5 ppm appear as doublets likely due to $^{13}$C-$^{31}$P coupling (see additional zoom spectrum S).

[*3] signals between 30.9 and 29.5 ppm overlap and are not resolved individually.

$^{31}$P NMR{$^{1}$H} (162 MHz, $CD_3OD$) δ = -0.16 ppm.

HR-MS (ESI negative): calculated m/z for $C_{37}H_{64}N_2O_8P^-$: 695.44058, found: 695.432 $[M]^-$.

Yield: 20.7 mg (29.7 µmol, quant.).

### Synthesis of $^{13}$C-labeled PC

### *PC(18:1/16:0[$^{13}C_{16}$]) (23)*

50.0 mg of ($^{13}C_{16}$)-palmitic acid (184 µmol, 1.7 eq.) were dissolved in 0.5 ml $CDCl_3$ and activated with 60.5 mg EDC·HCl (316 µmol, 3.0 eq.) and 6.4 mg DMAP (52.4 µmol, 0.5 eq.) for 15 minutes at room temperature. 55.0 mg Lyso-PC 18:1 (105 µmol, 1 eq.) was dissolved in 1 ml $CDCl_3$ and added. The reaction mixture was stirred at room temperature overnight. The solvent was removed and the crude product was purified by flash chromatography (eluents chloroformCHCl$_3$/MeOH/H$_2$O 70/30/2). The product **23** was isolated as a colourless oil.

$^{1}$H NMR (400 MHz, $CD_3OD$) δ = 5.35 ($m_c$, 2H), 5.25 ($m_c$, 1H), 4.44 (dd, $J$ = 12.0, 3.1 Hz, 1H), 4.28 ($m_c$, 2H), 4.18 (dd, $J$ = 12.0, 7.0 Hz, 1H), 4.00 (t, $J$ = 6.2 Hz, 2H), 3.68 – 3.60 (m, 2H), 3.23 (s, 9H), 2.55 – 2.44 (m, 1H), 2.39 – 2.28 (m, 2H), 2.24 – 2.12 (m, 1H), 2.04 ($m_c$, 4H), 1.77 ($m_c$, 1H), 1.67 – 1.55 (m, 2H), 1.56 – 1.39 (m, 12H), 1.39 – 1.22 (m, 22H), 1.23 – 1.00 (m, 13.5 H), 0.90 (t, $J$ = 6.9 Hz, 3H), 0.79 – 0.69 (m, 1.5 H) ppm.

$^{13}$C NMR{$^{1}$H} (101 MHz, $CD_3OD$) δ = 175.21, 174.90, 174.64, 174.33, 130.93, 130.77, 71.86, 67.52, 64.87, 63.70, 60.50, 54.70, 35.55, 35.38, 35.21, 35.04, 34.98, 34.81, 34.64, 34.47, 33.42, 33.08, 32.82, 30.77, 30.47, 28.16, 26.36, 26.02, 25.69, 24.08, 23.73, 23.39, 14.61, 14.47, 14.26 ppm.

$^{31}$P NMR{$^{1}$H} (162 MHz, $CD_3OD$) δ = -0.57 ppm.

HR-MS (ESI positive): calculated m/z for $(C_{12})_{26}(C_{13})_{16}H_{82}NO_8P$: 775.63210, found: 776.642 $[M+H]^+$.

Yield: 9.7 mg (13 µmol, 12 %).

### NMR Characterization of new compounds

##### 2-(dec-9-ynoyl)cyclohexan-1-one (3)

***[1]H spectrum***

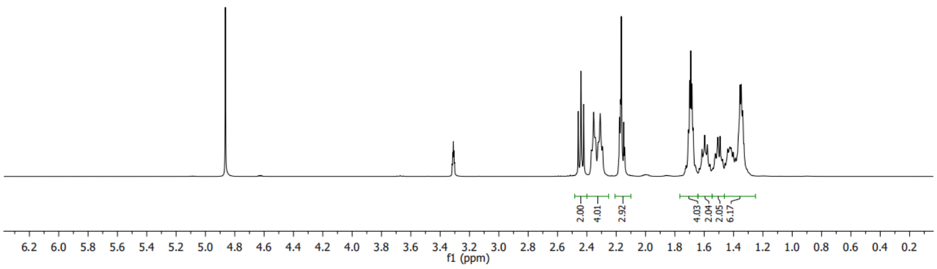

***[13]C spectrum***

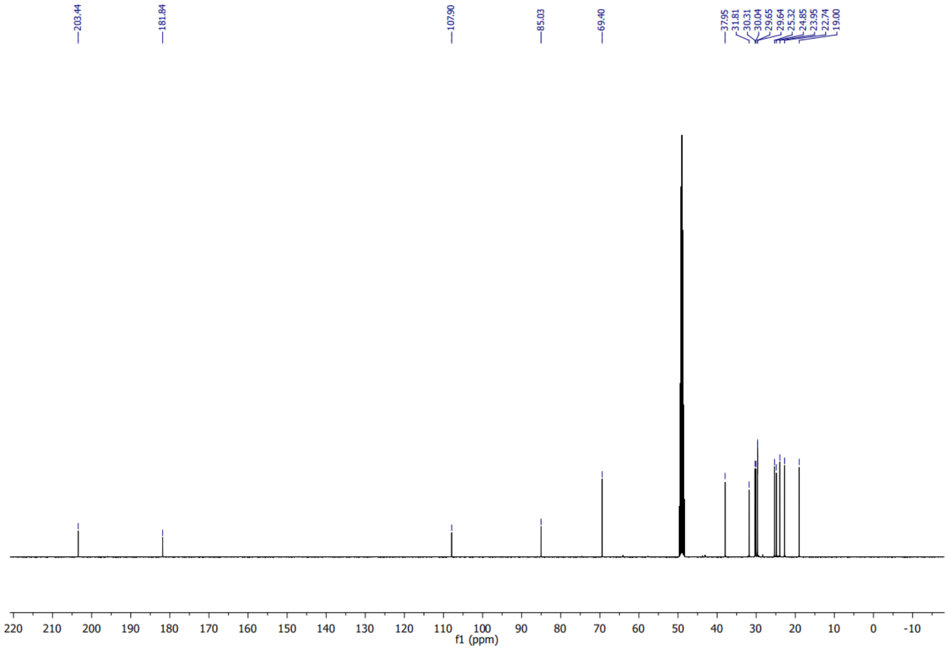

### 7-Oxohexadec-15-ynoic acid (4)

#### *¹H spectrum*

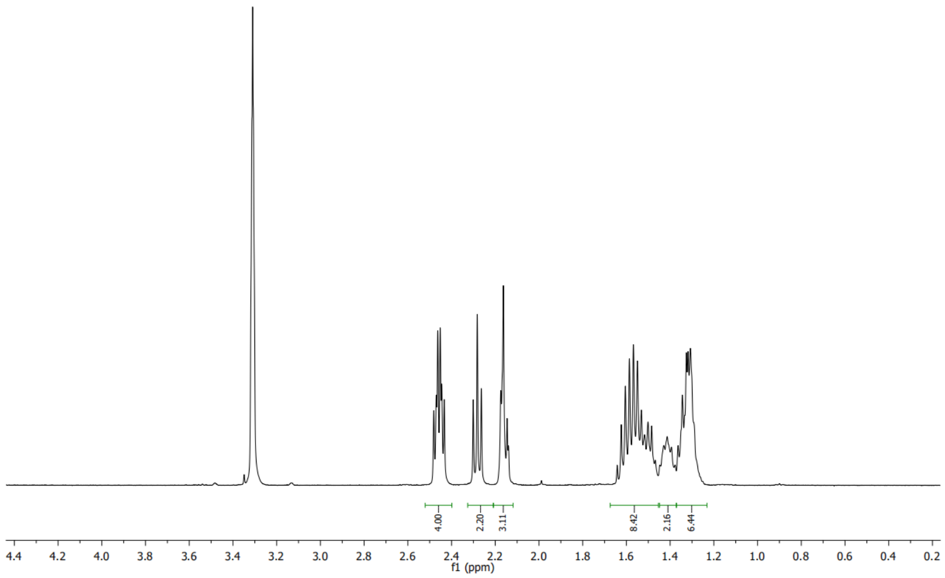

#### *¹³C spectrum*

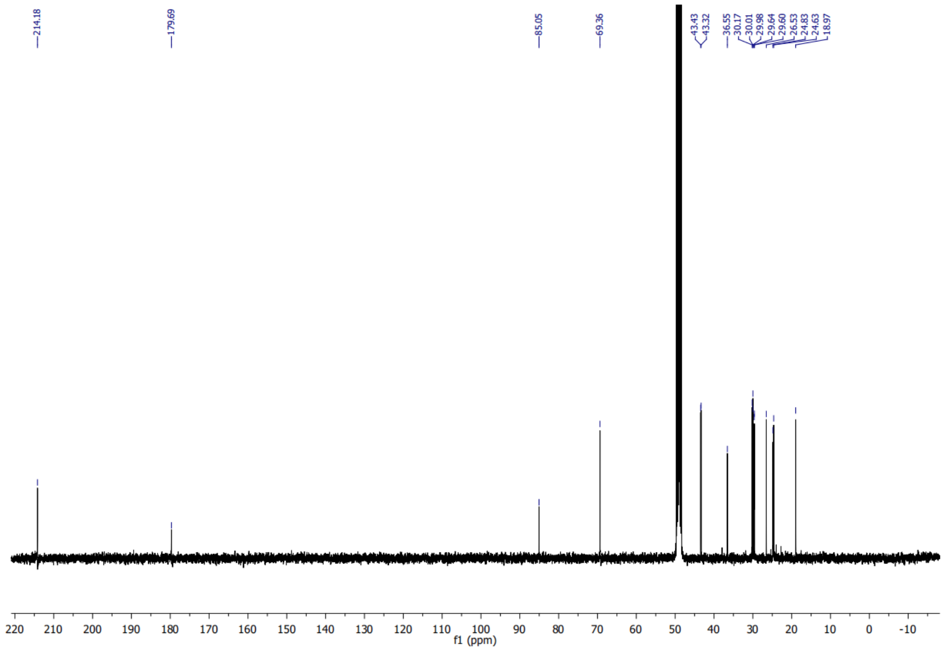

### 6-(3-(non-8-yn-1-yl)-3H-diazirin-3-yl)hexanoic acid (5)

#### *[1]H spectrum*

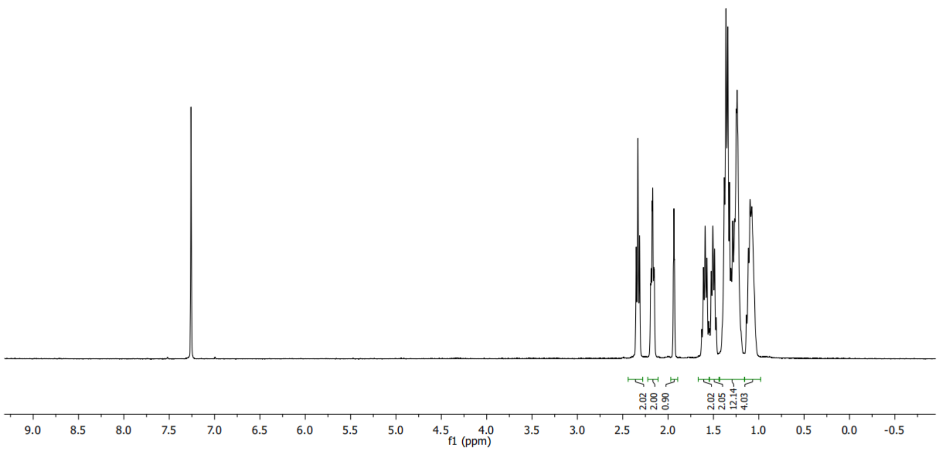

#### *[13]C spectrum*

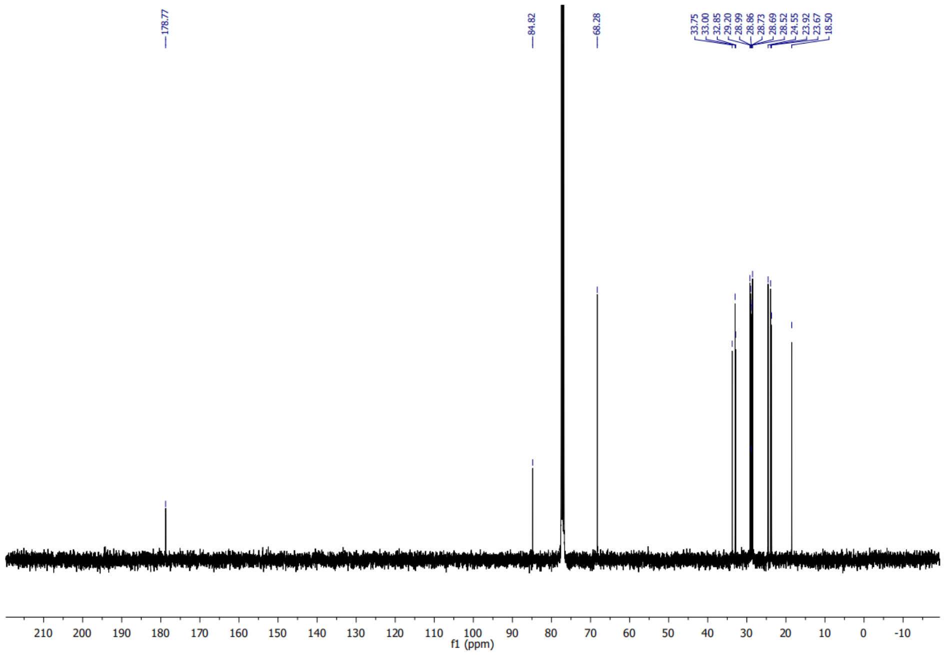

#### (2R)-3-(((2-((((9H-fluoren-9-yl)methoxy)carbonyl)amino)ethoxy)(hydro-xy) phosphoryl)oxy)-2-hydroxypropyl oleate (7)

##### *[1]H spectrum of (7)*

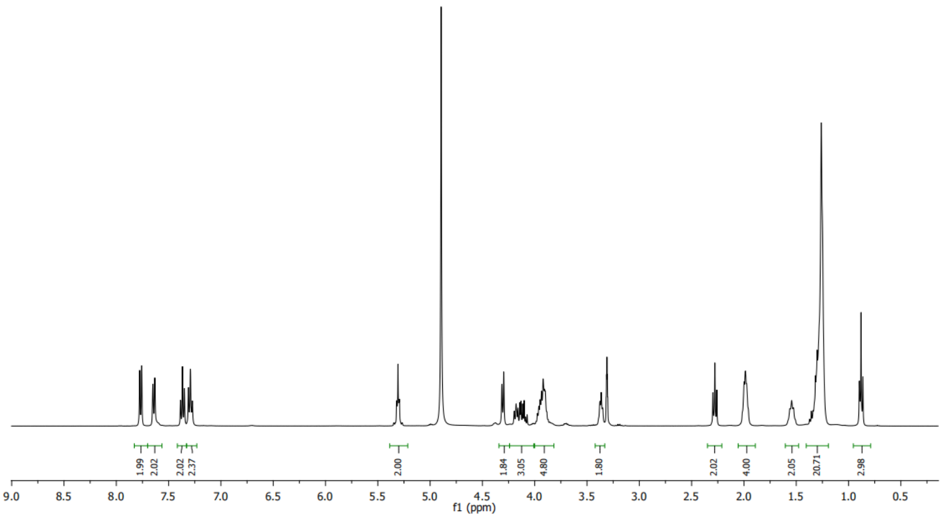

##### *[13]C spectrum of (7)*

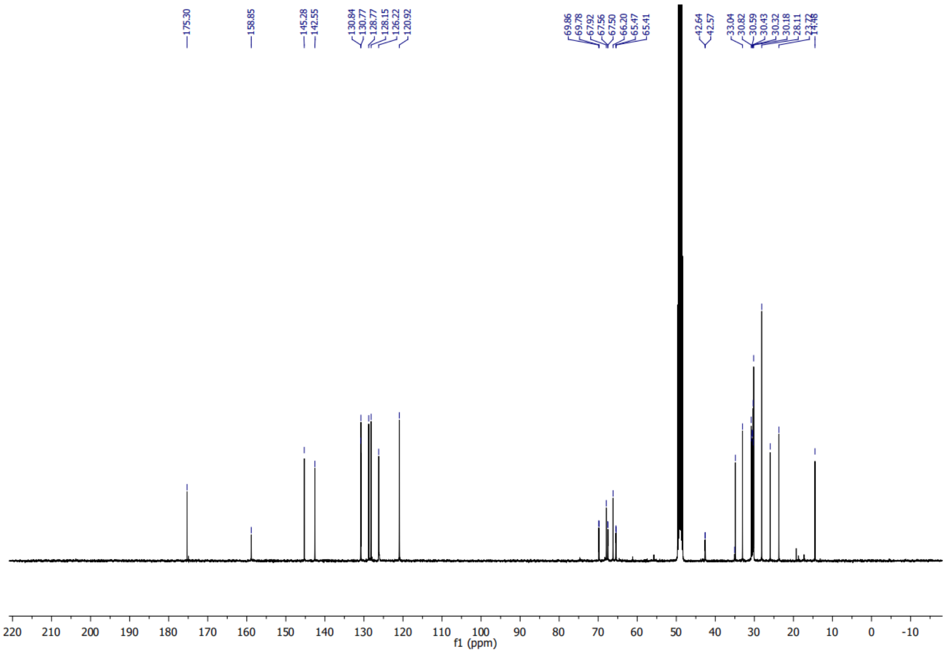

#### *Zoom of $^{13}$C spectrum of (7)*

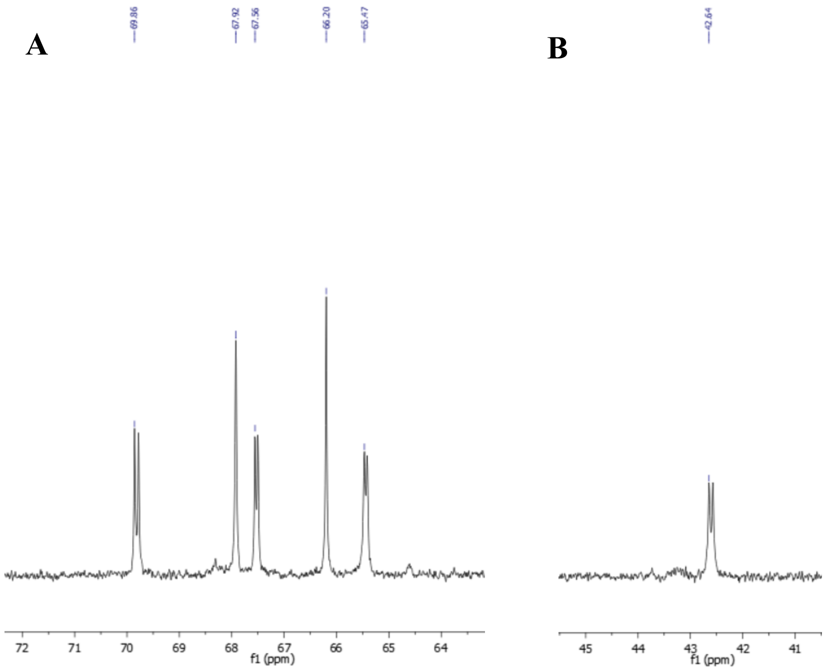

#### *$^{31}$P spectrum of (7)*

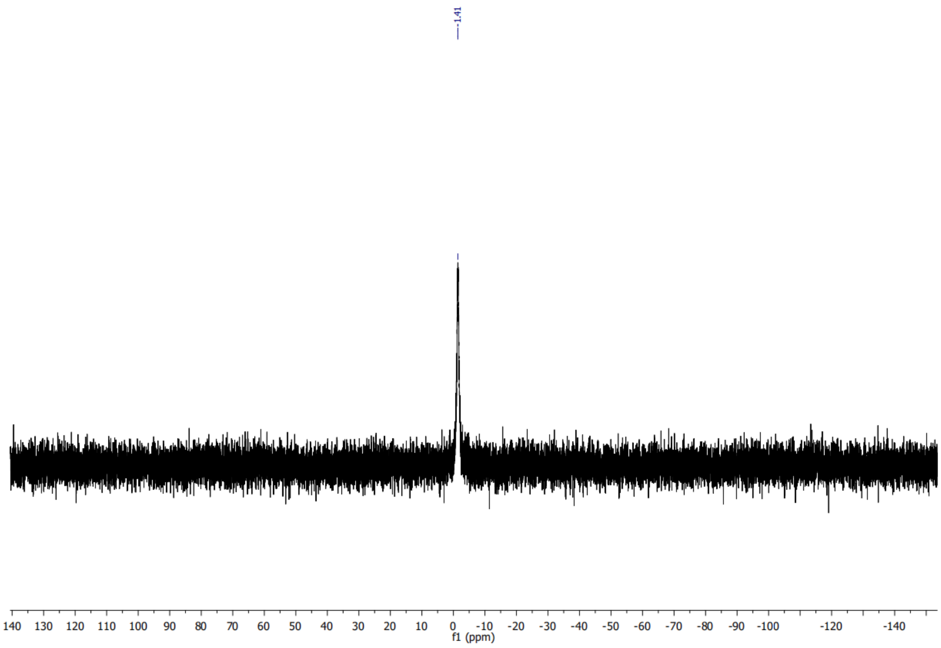

# PE(18:1/Y) (8)

#### *[1]H spectrum of (8)*

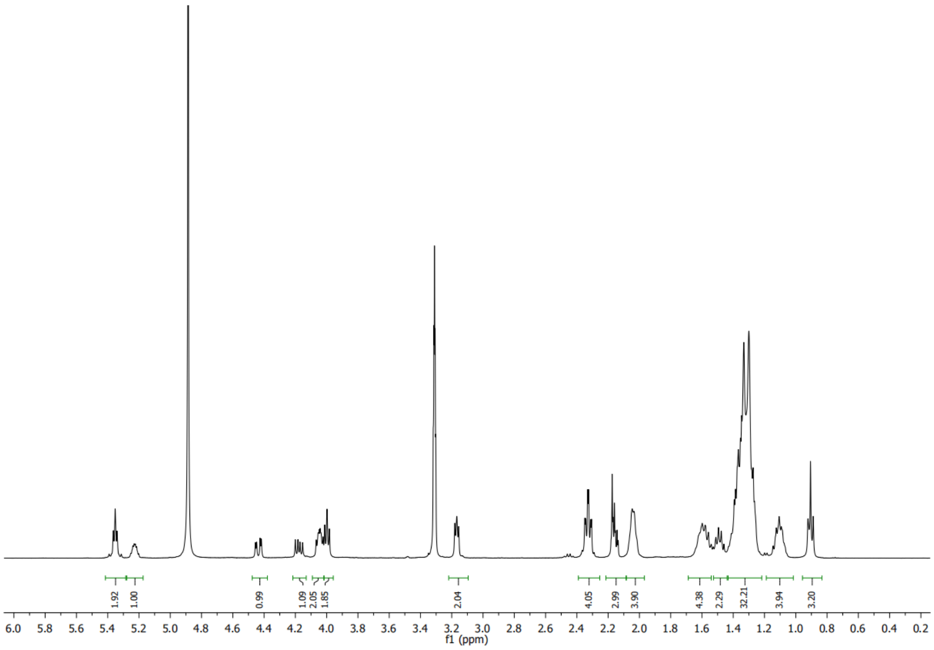

#### [13]C spectrum of (8)

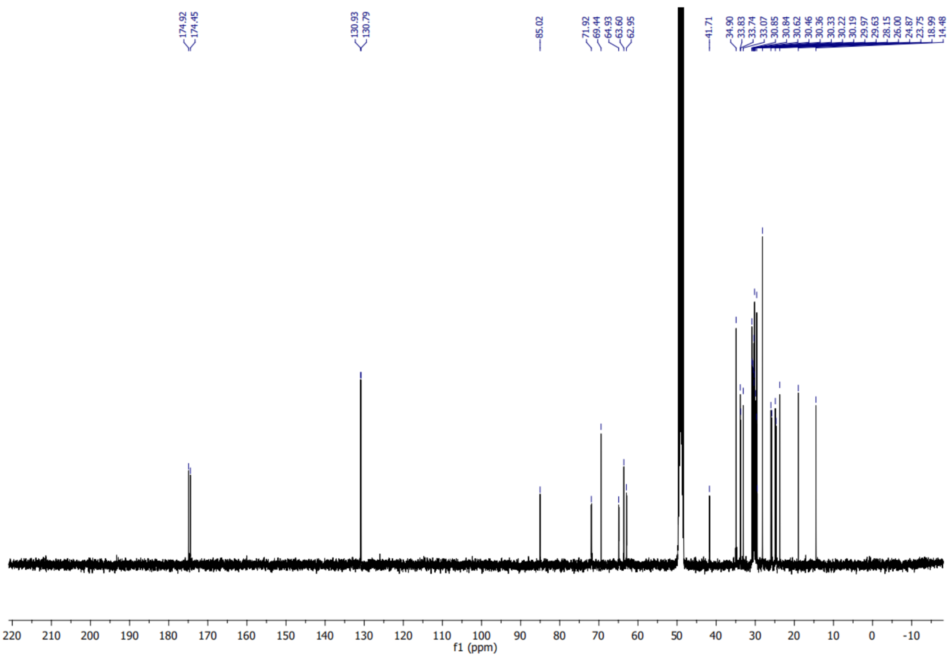

#### *Zoom $^{13}C$ spectrum of 8*

C

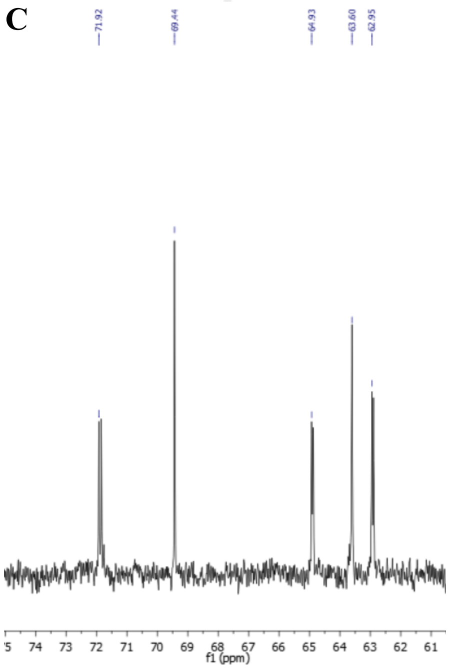

D

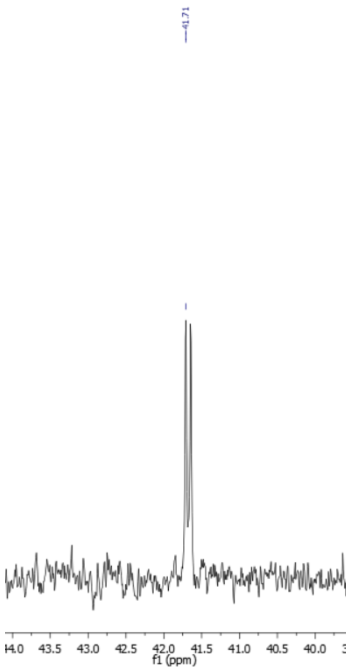

#### *$^{31}P$ spectrum of (8)*

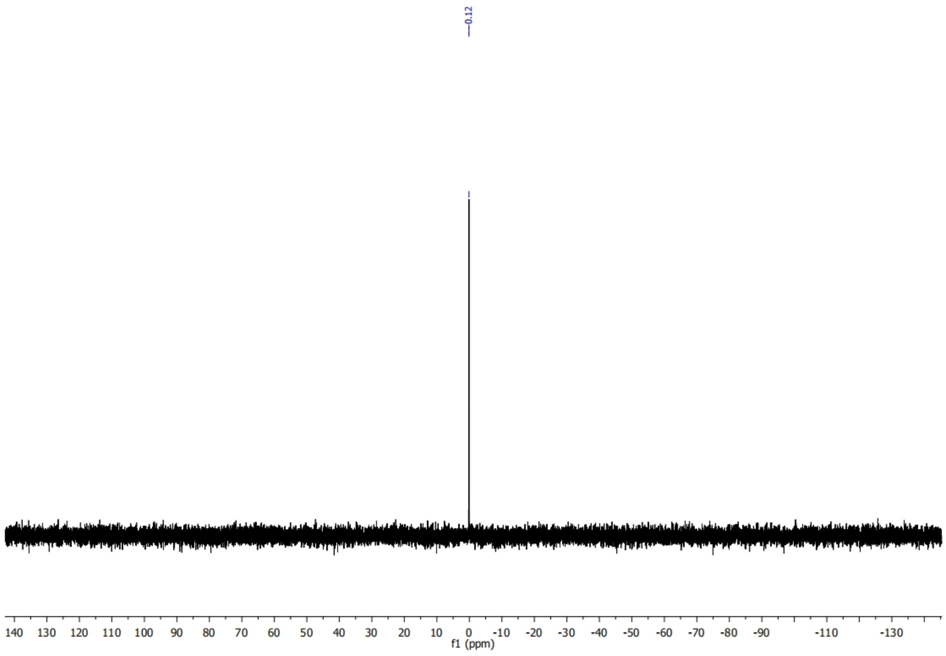

## PC(16:0/Y) (10a)

##### *[1]H Spectrum of (10a)*

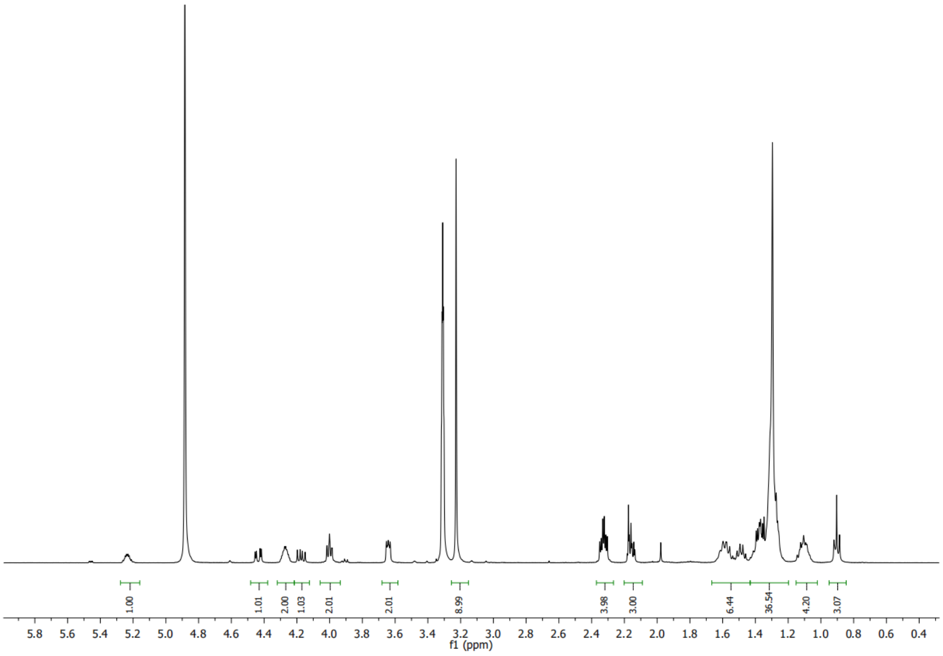

##### *[13]C spectrum of (10a)*

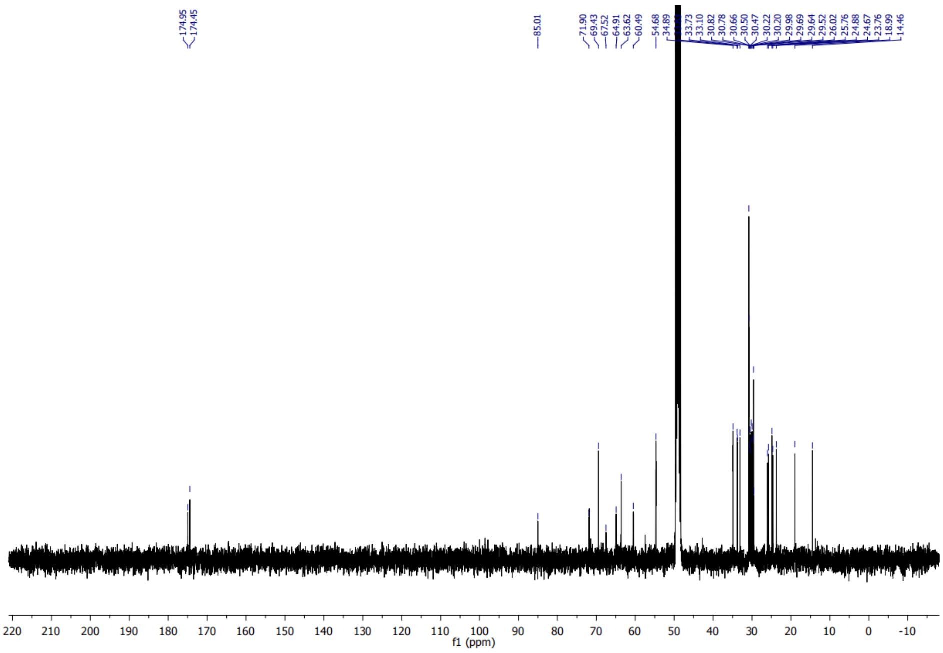

#### *Zoom of $^{13}$C spectrum of (10a)*

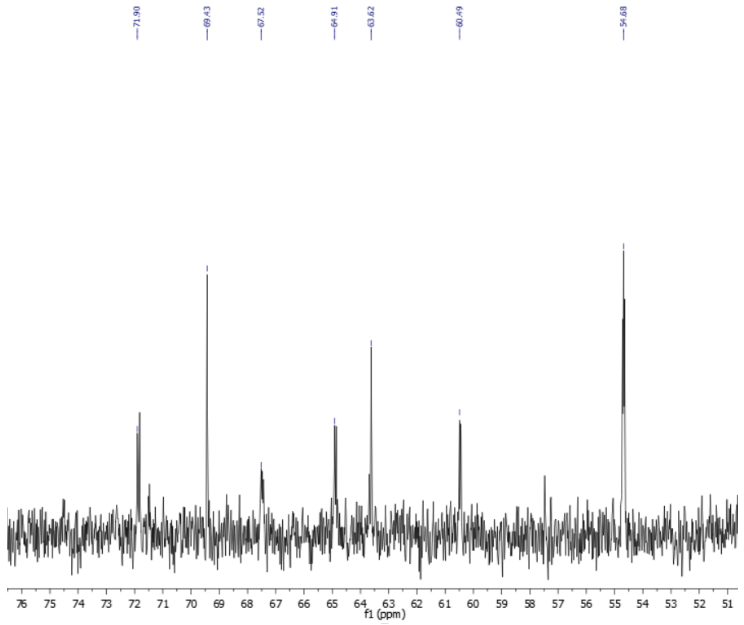

#### *$^{31}$P Spectrum of (10a)*

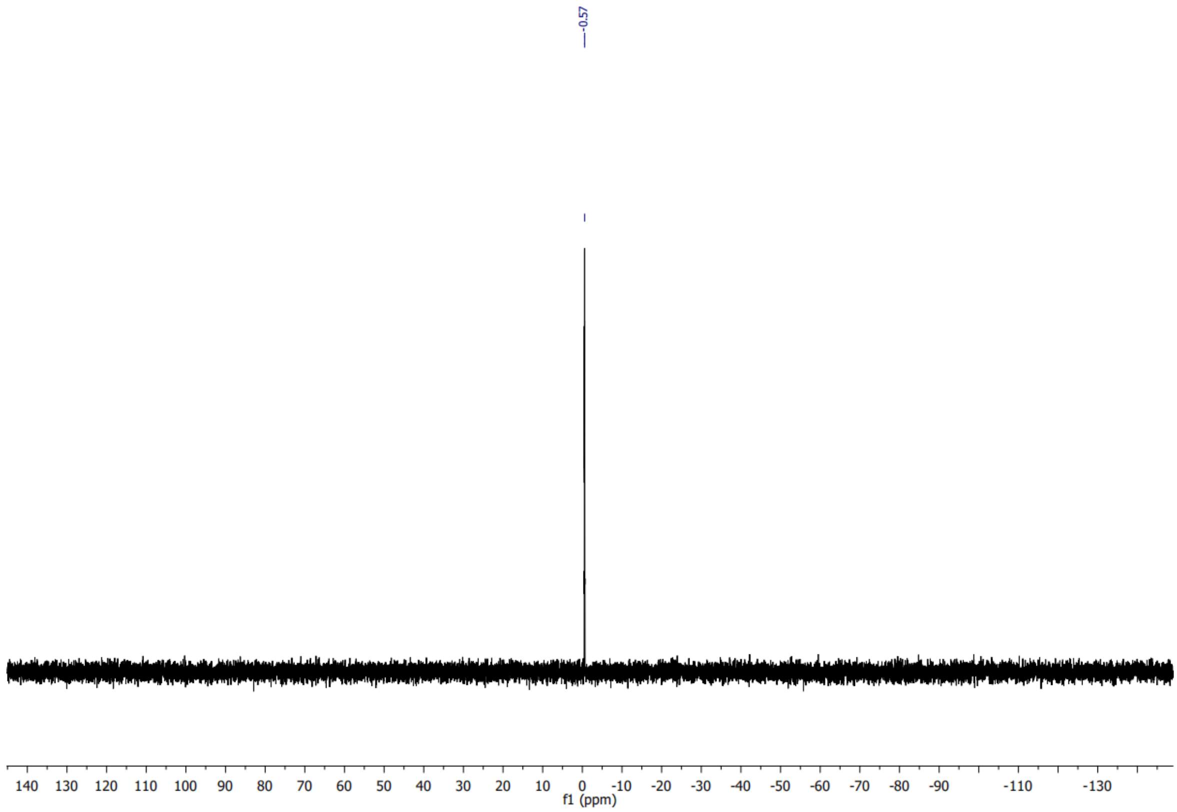

# PC(18:1/Y) (10b)

#### *[1]H spectrum of (10b)*

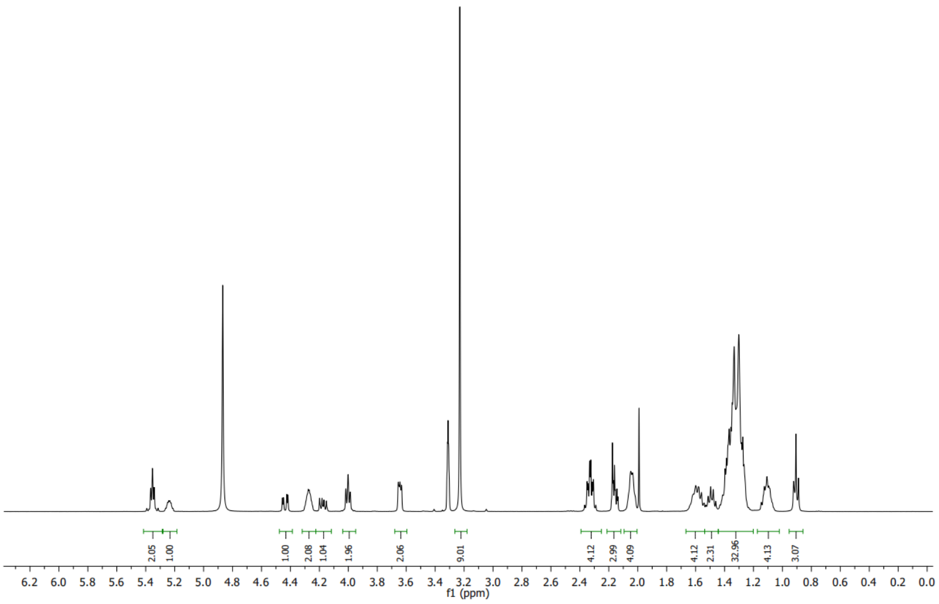

#### *[13]C Spectrum of (10b)*

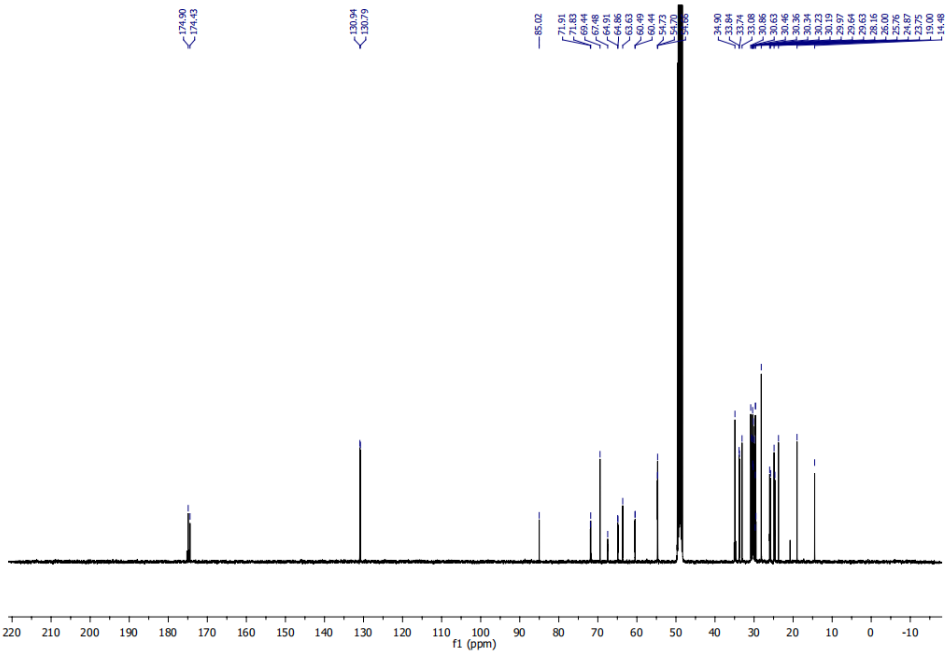

#### *Zoom $^{13}C$ spectrum (10b)*

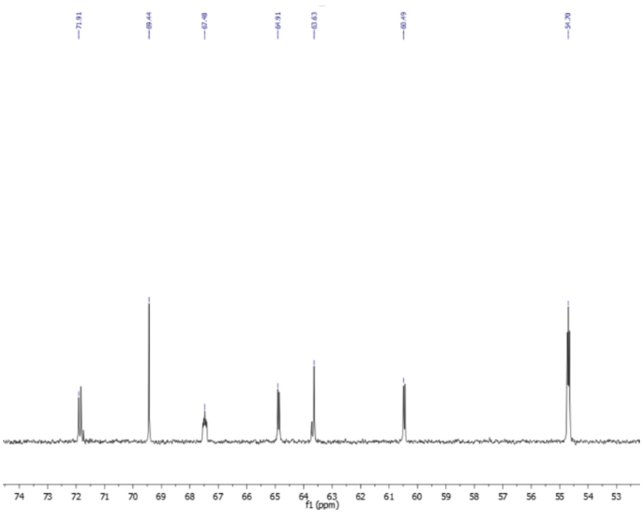

#### *$^{31}P$ spectrum of (10b)*

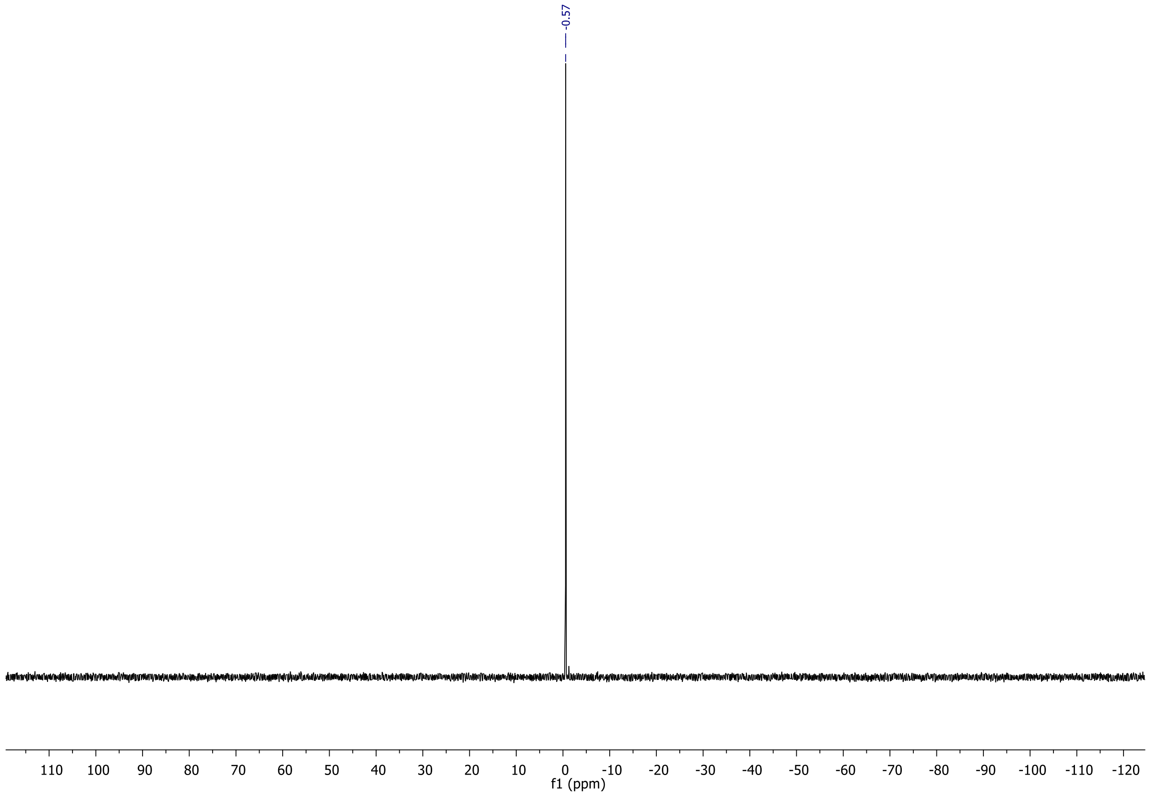

#### *sn1*-Dimethoxytrityl-*sn2*-BIF PC (13a)

##### *[1]H spectrum of (13a)*

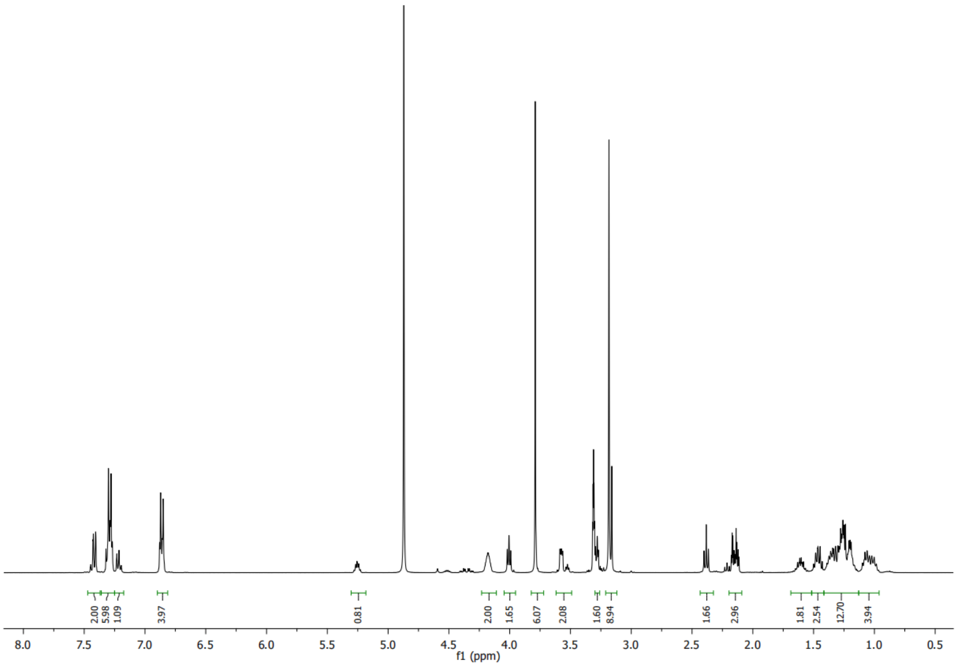

##### *[13]C spectrum of (13a)*

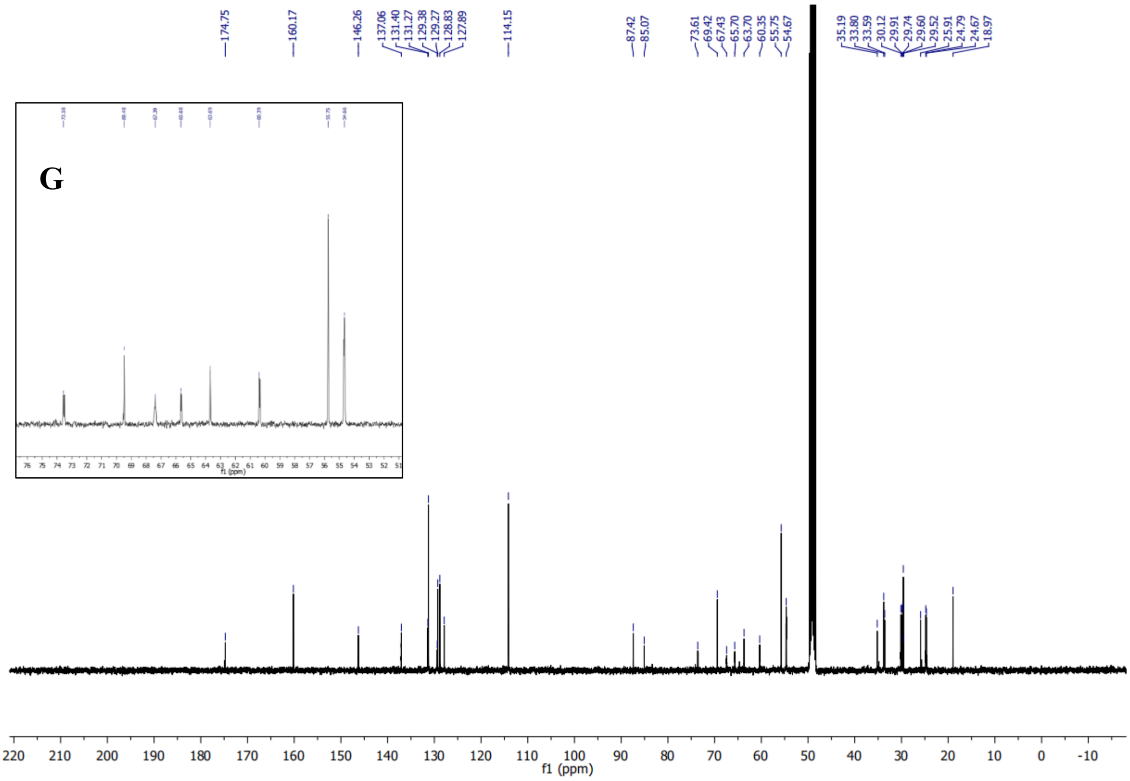

#### *[1]H spectrum (13c)*

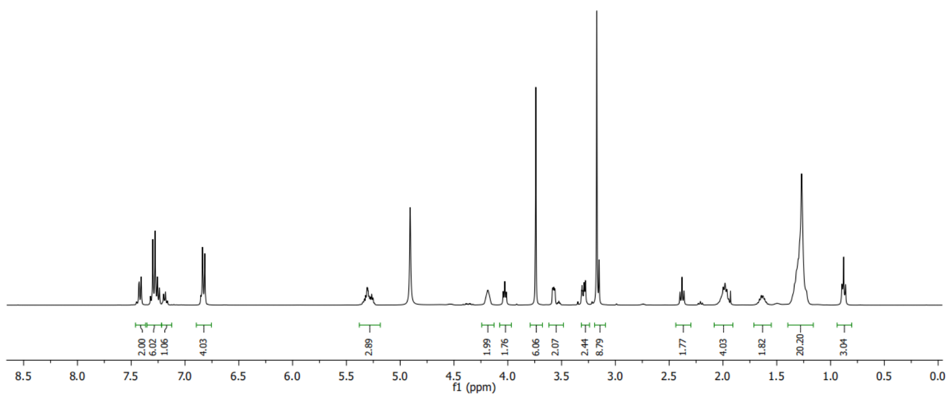

#### *[13]C Spectrum of (13c)*

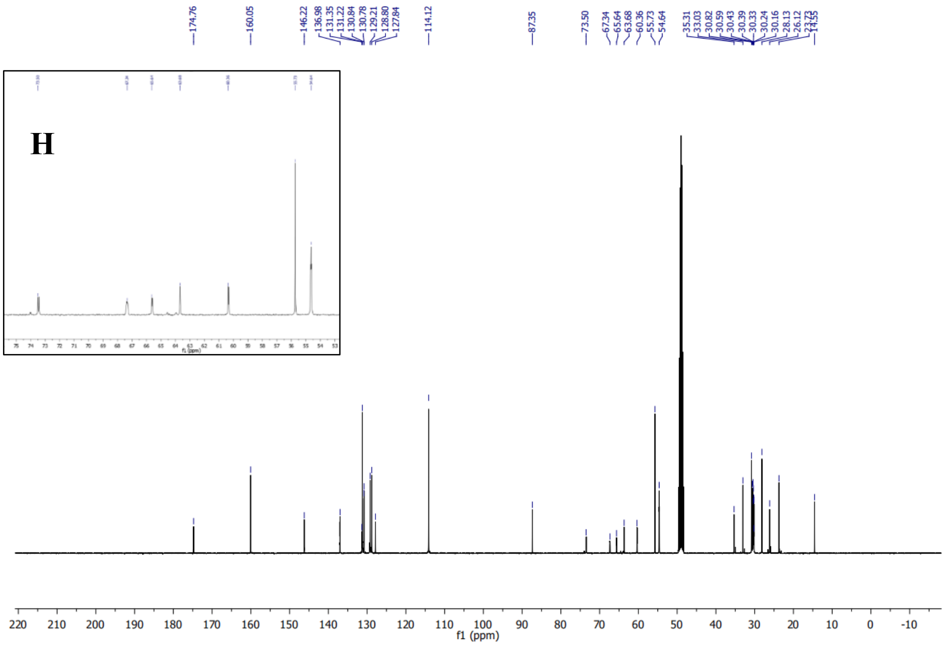

#### *sn1*-Dimethoxytrityl-*sn2*-arachidonyl PC (13d)

##### *[1]H spectrum of (13d)*

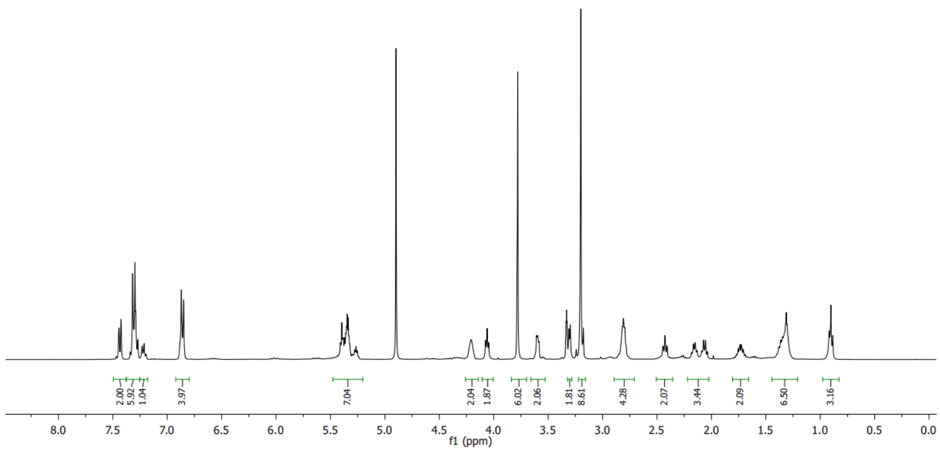

##### *[13]C Spectrum of (13d)*

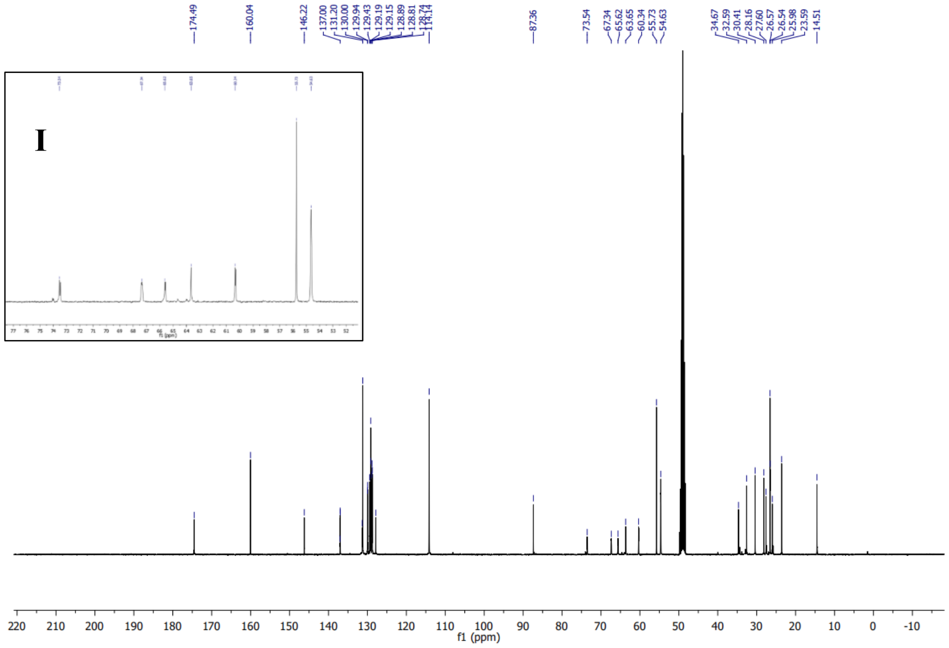

#### *sn2*-BIF-*sn*-lysophosphocholine (14a)

##### *¹H spectrum of (14a)*

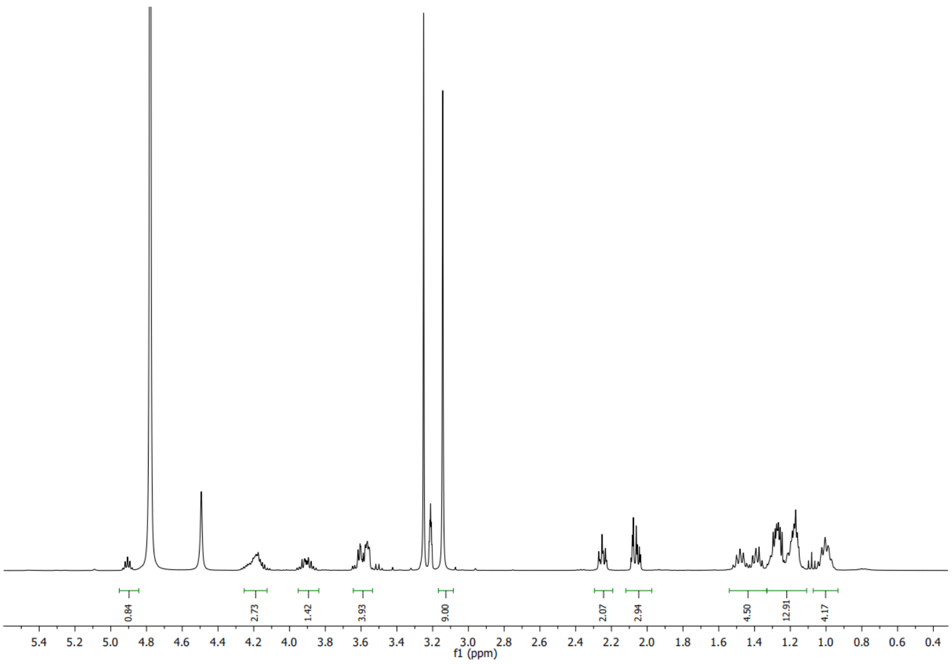

##### *¹³C spectrum of (14a)*

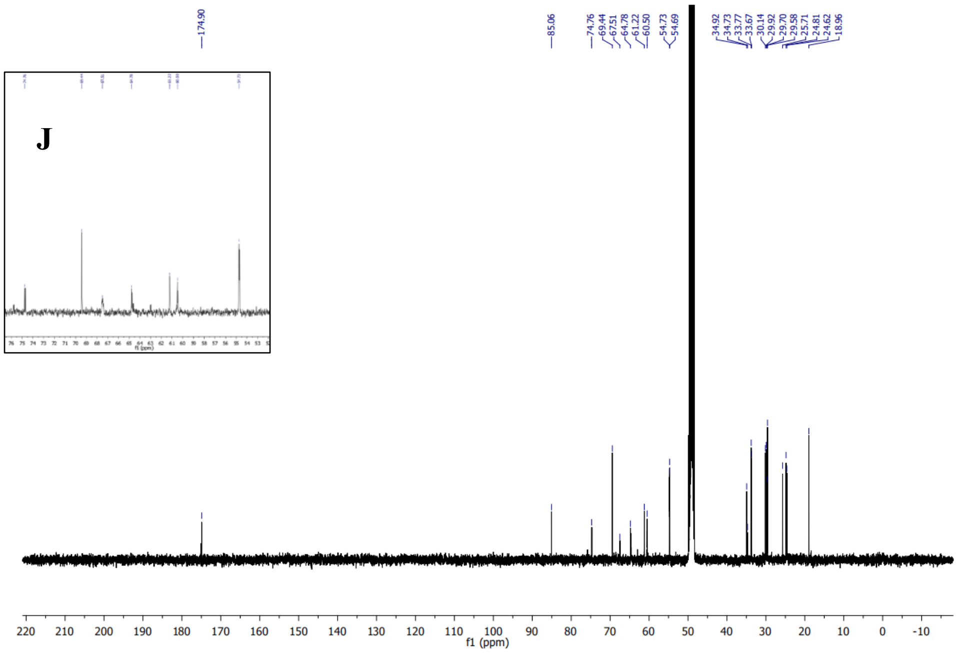

#### *sn2*-Oleoyl-*sn*-lysophosphocholine (14c)

##### *[1]H spectrum of (14c)*

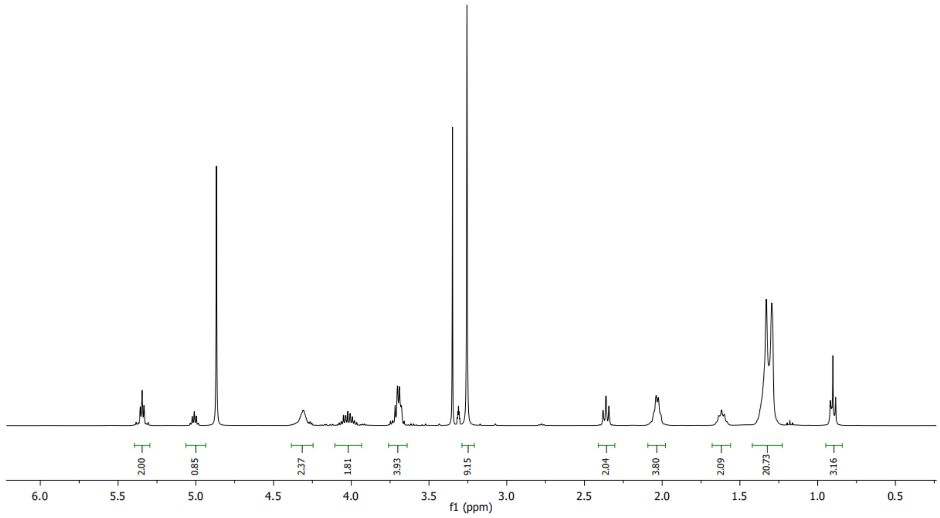

##### *[13]C spectrum of (14c)*

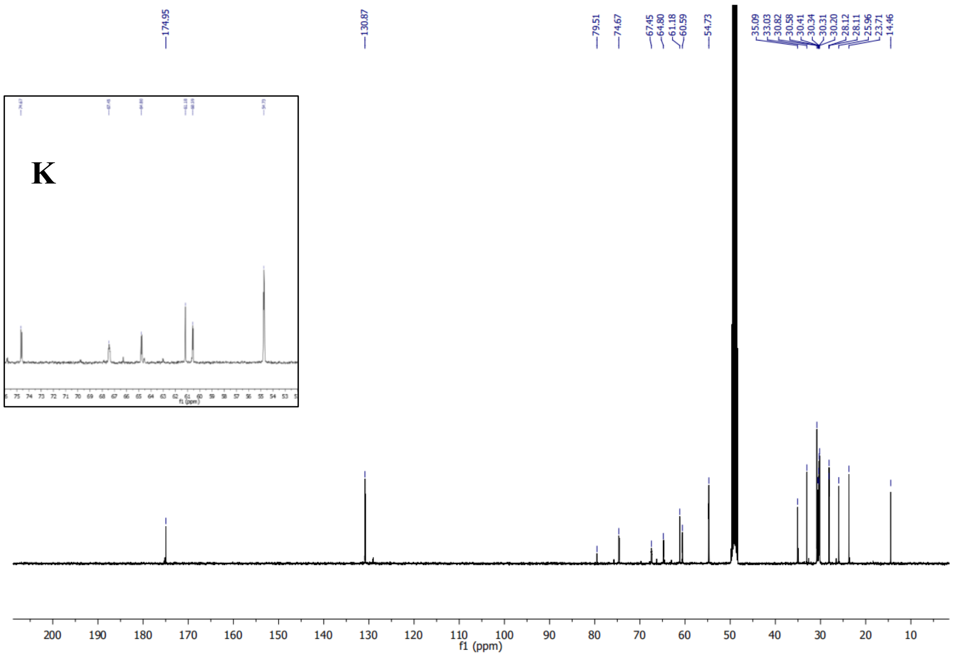

#### *sn2*-Arachidonyl-*sn*-lysophosphocholine (14d)

##### *[1]H Spectrum of (14d)*

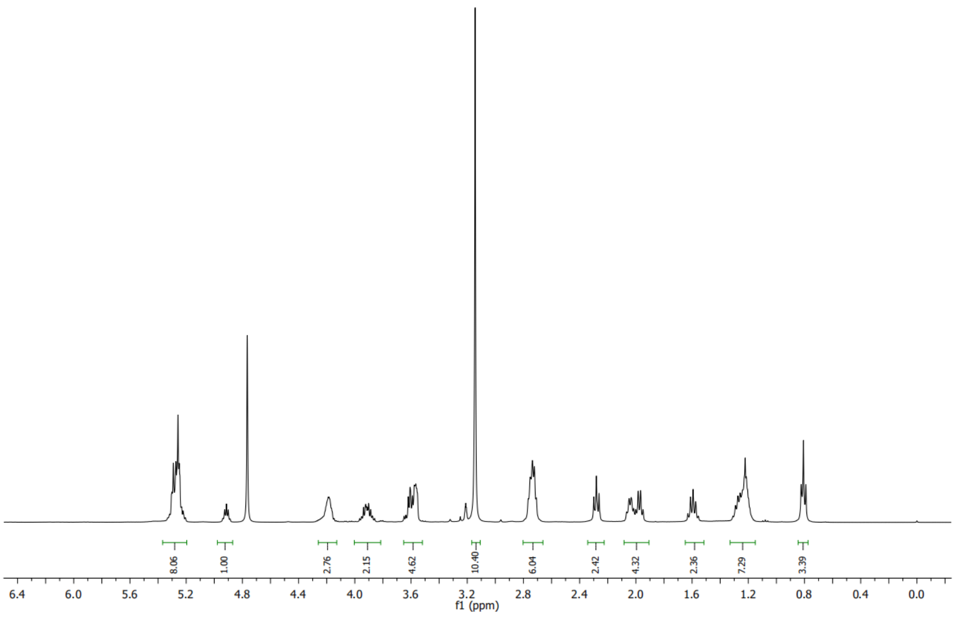

##### *[13]C spectrum of (14d)*

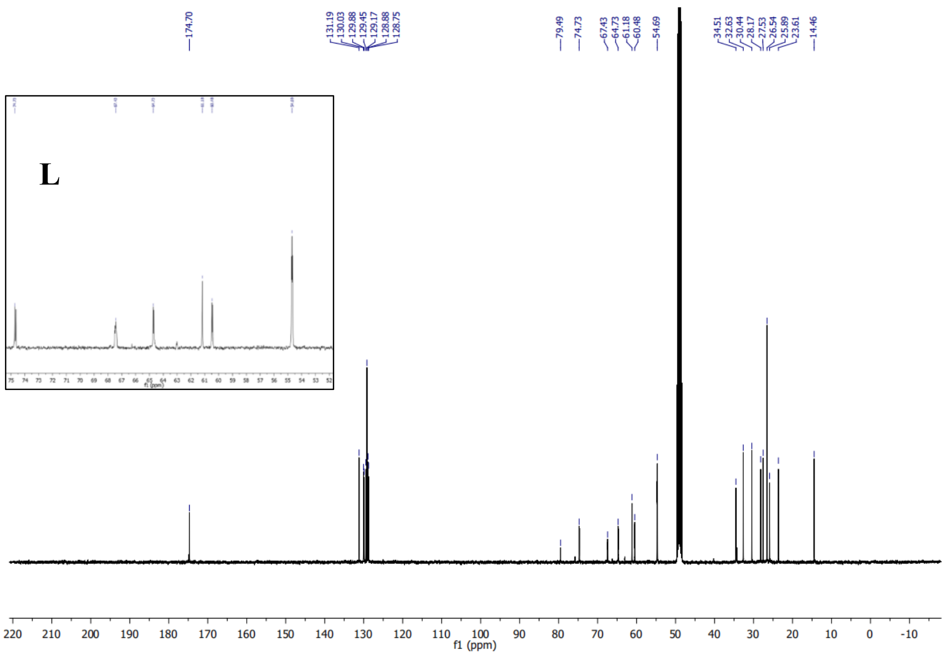

# PC(20:4/Y) (15a)

#### *[1]H spectrum of (15a)*

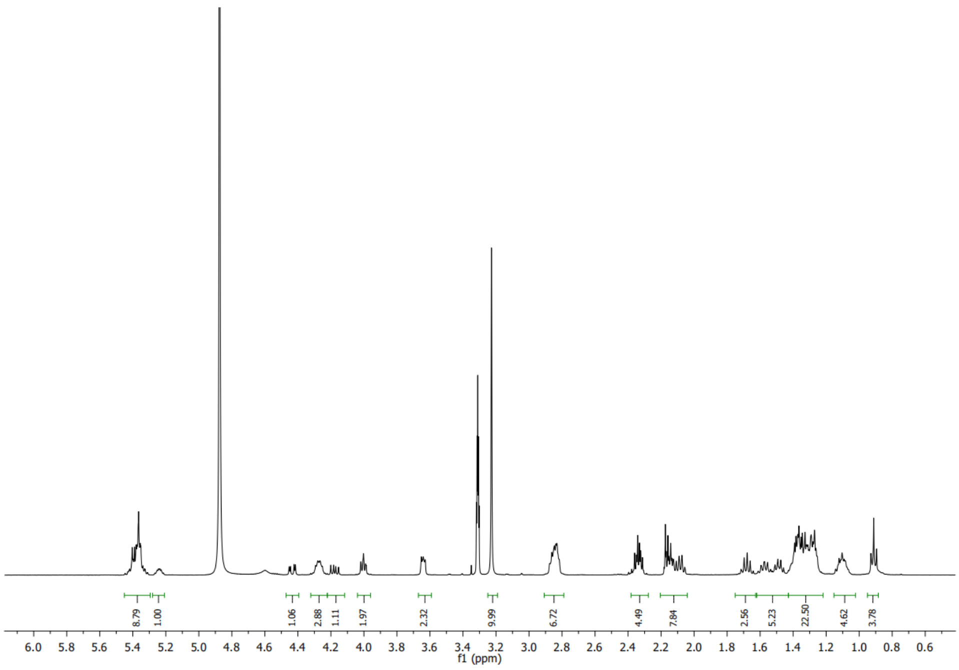

#### *[13]C spectrum of (15a)*

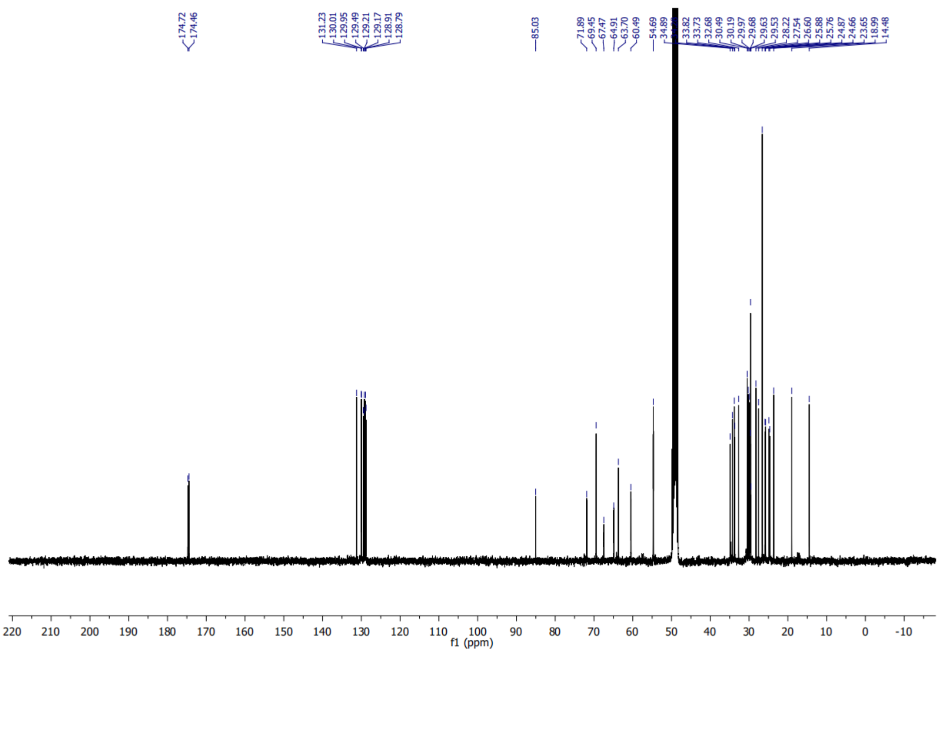

#### *Zoom of $^{13}$C spectrum of (15a)*

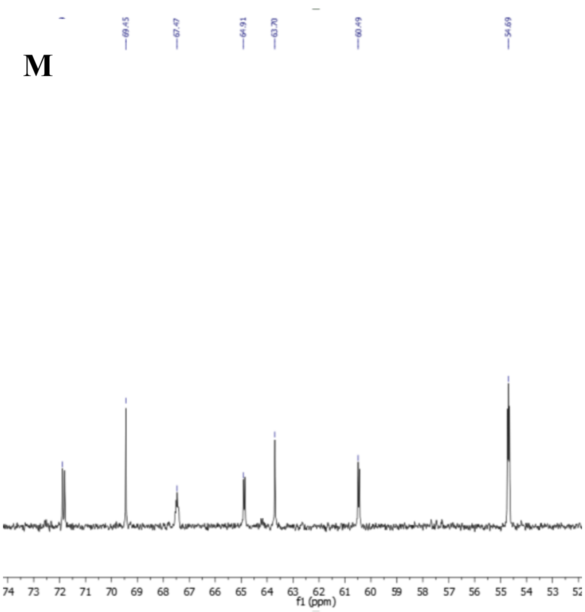

#### *$^{31}$P spectrum of (15a)*

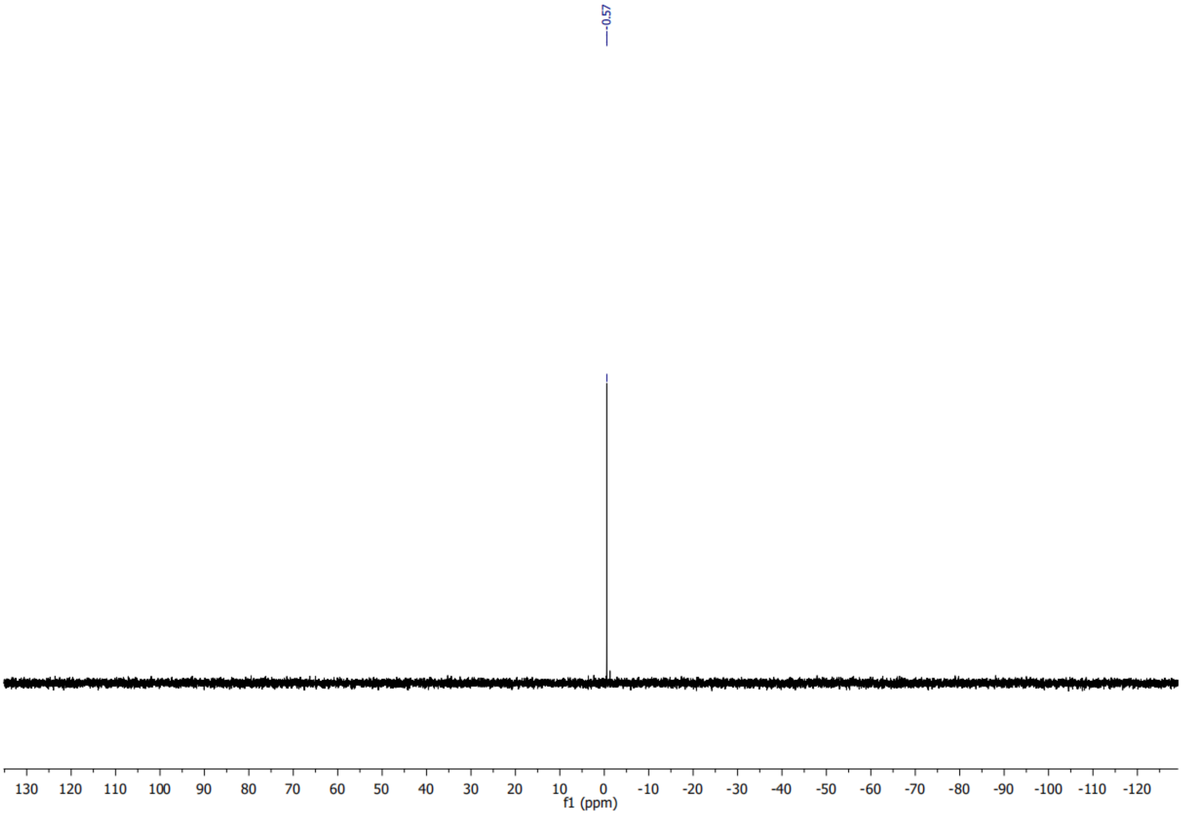

# PC(Y/16:0) (15b)

#### *[1]H spectrum of (15b)*

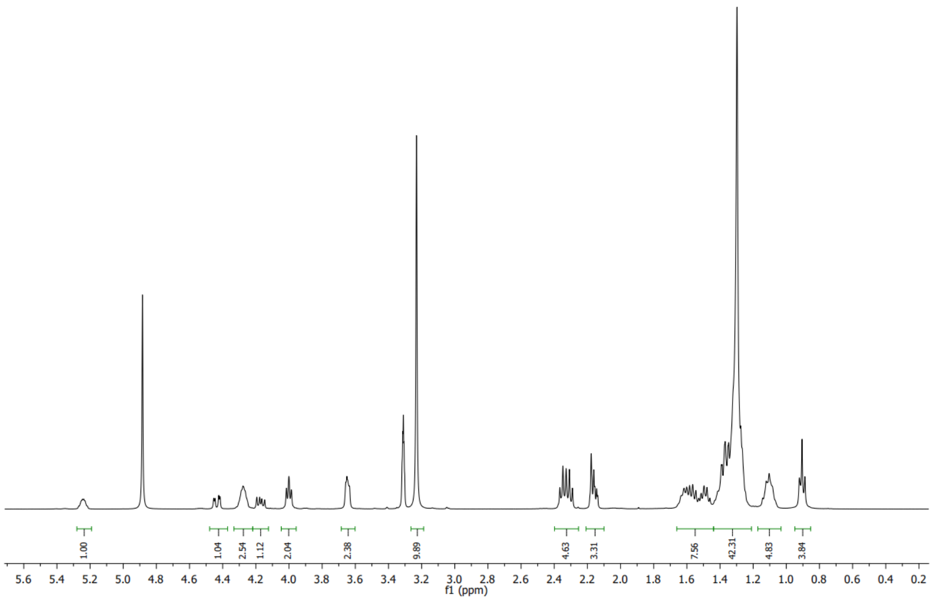

#### *[13]C spectrum of (15b)*

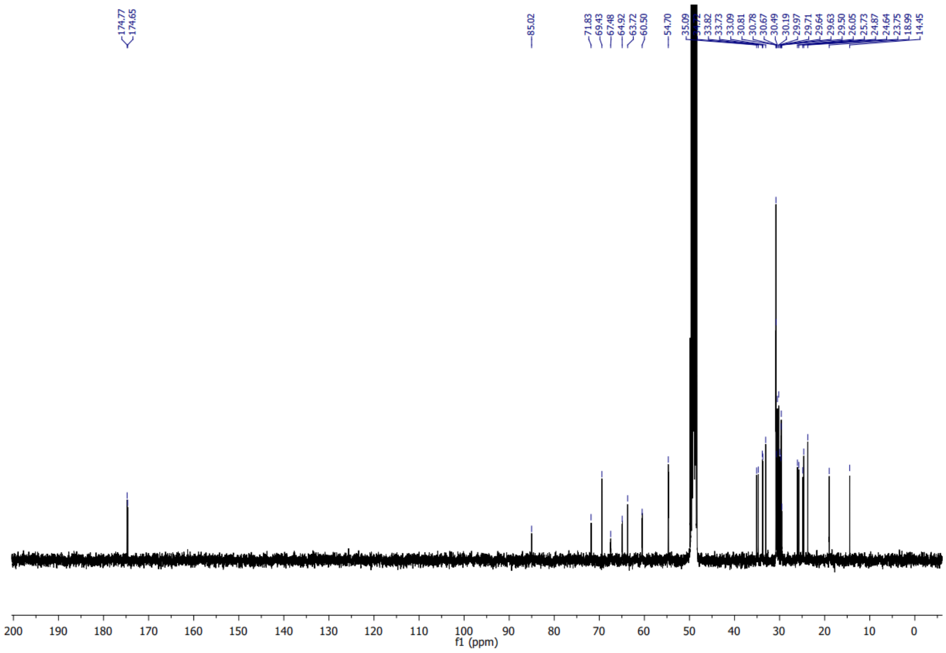

##### *Zoom of $^{13}C$ spectrum of (15b)*

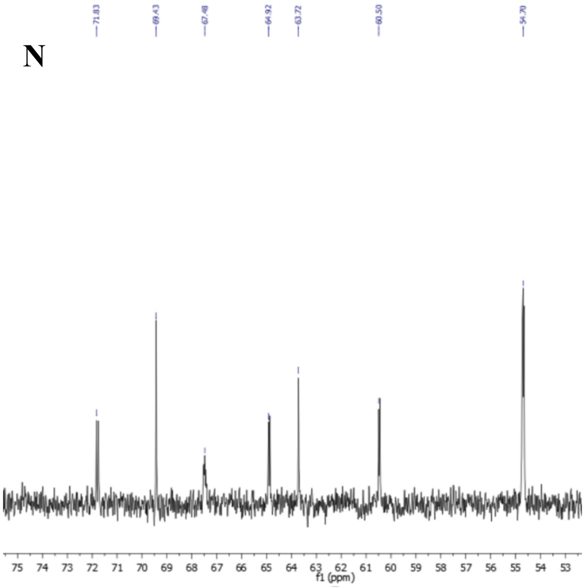

##### *$^{31}P$ Spectrum of (15b)*

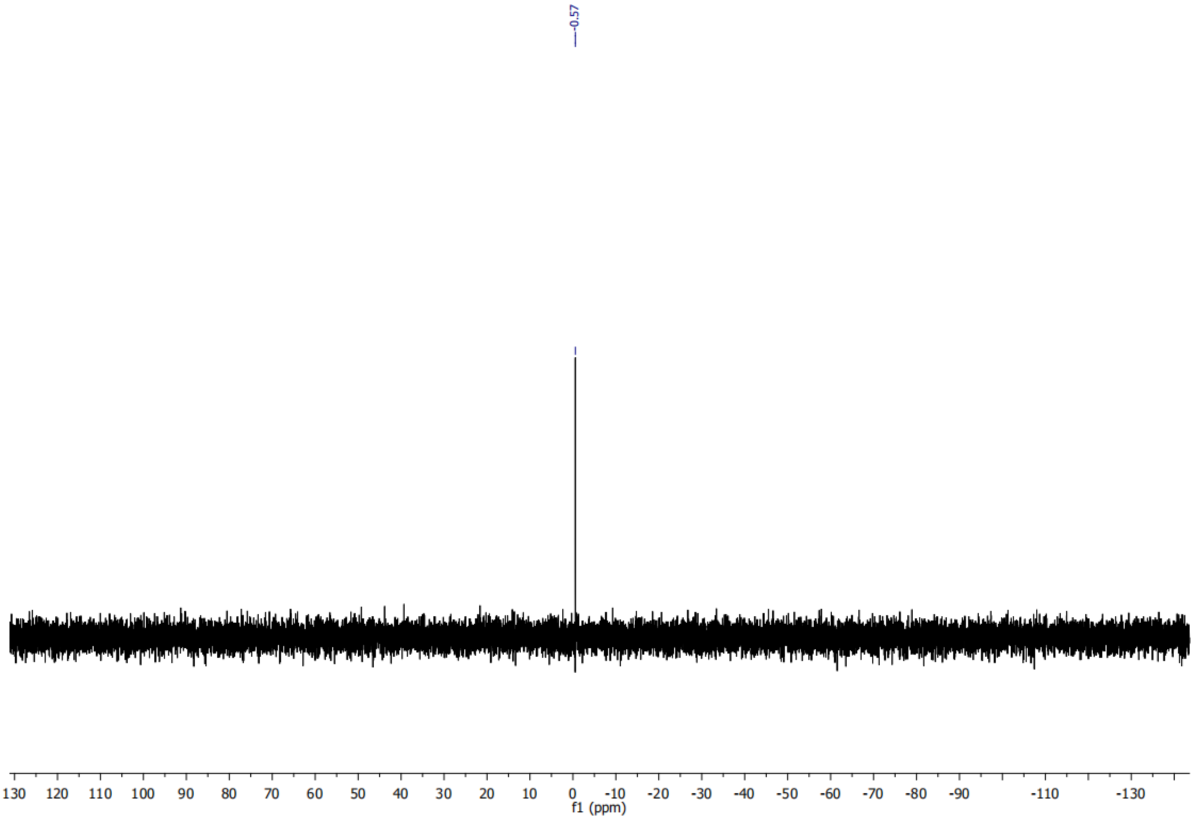

# PC(Y/18:1) (15c)

#### *[1]H spectrum of (15c)*

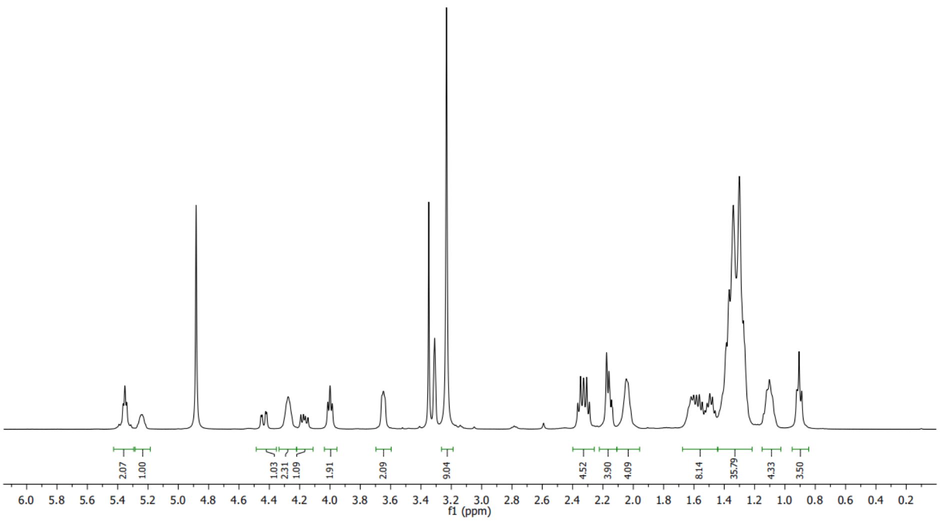

#### *[13]C spectrum of (15c)*

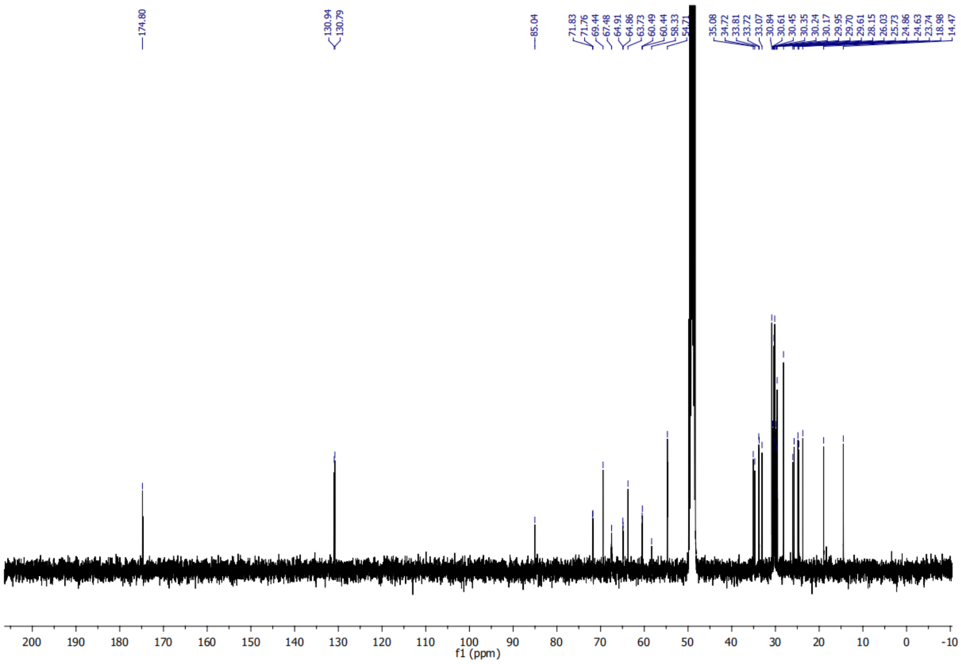

#### *Zoom of $^{13}C$ spectrum of (15c)*

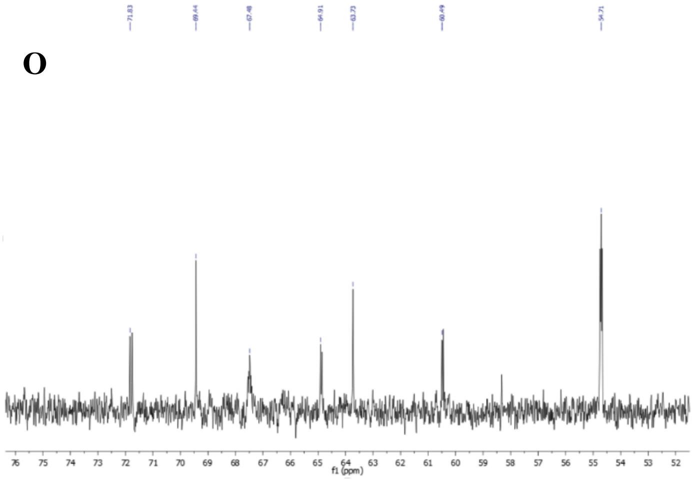

#### *$^{31}P$ spectrum of (15c)*

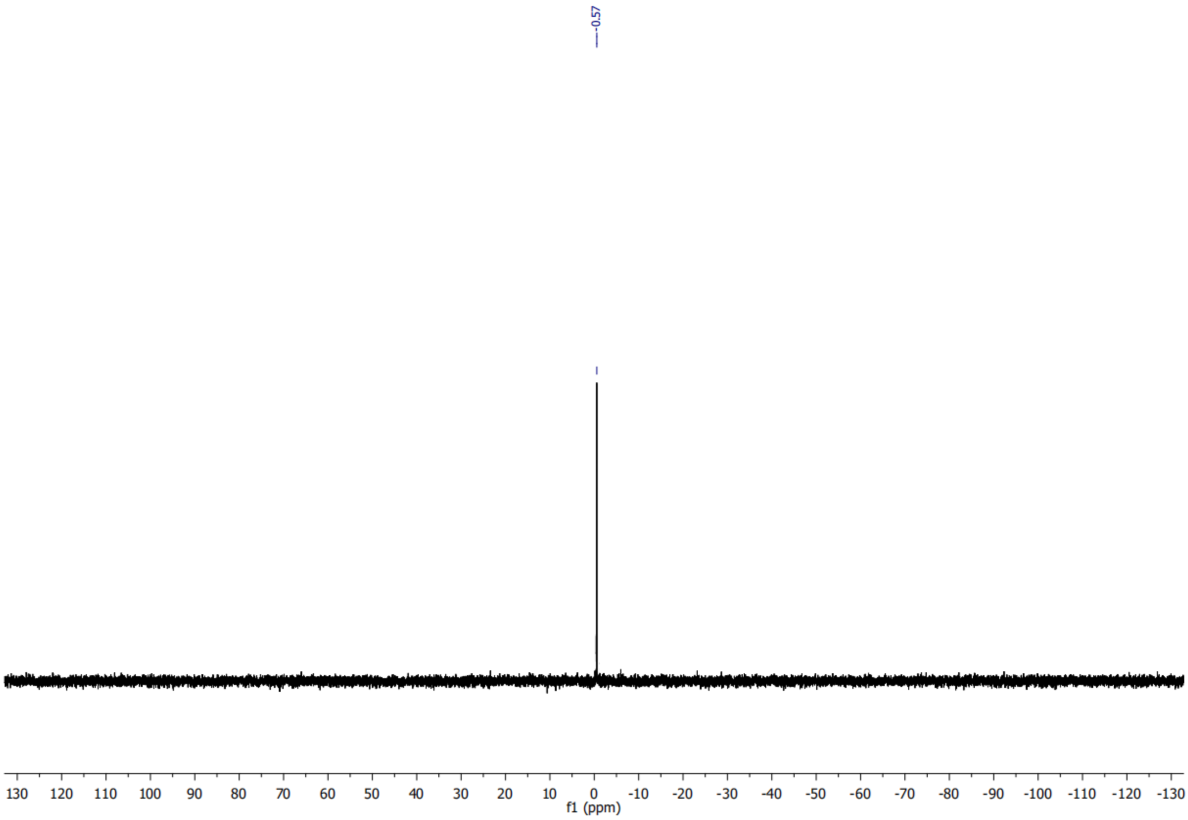

# PC(Y/20:4) (15d)

#### *¹H spectrum of (15d)*

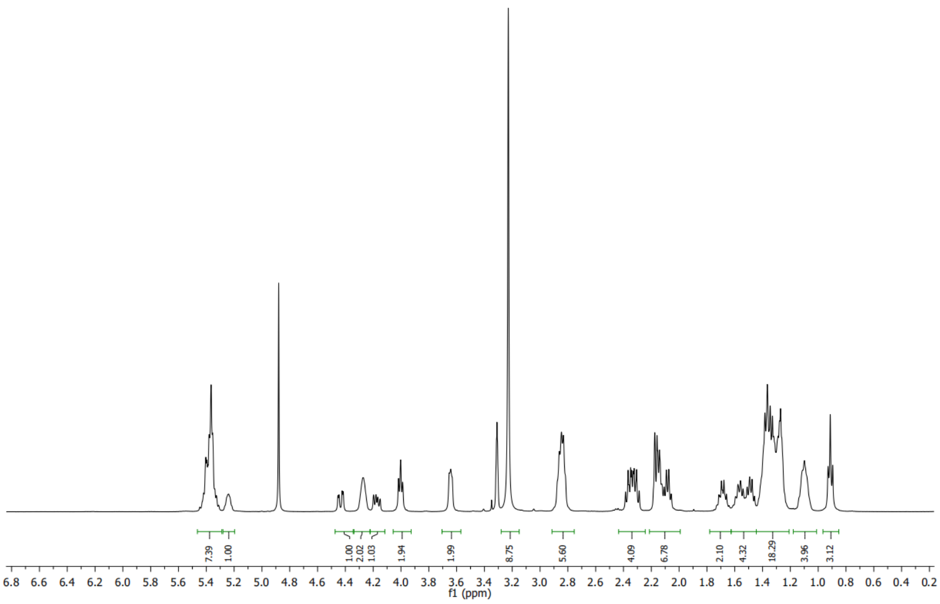

#### *¹³C spectrum of (15d)*

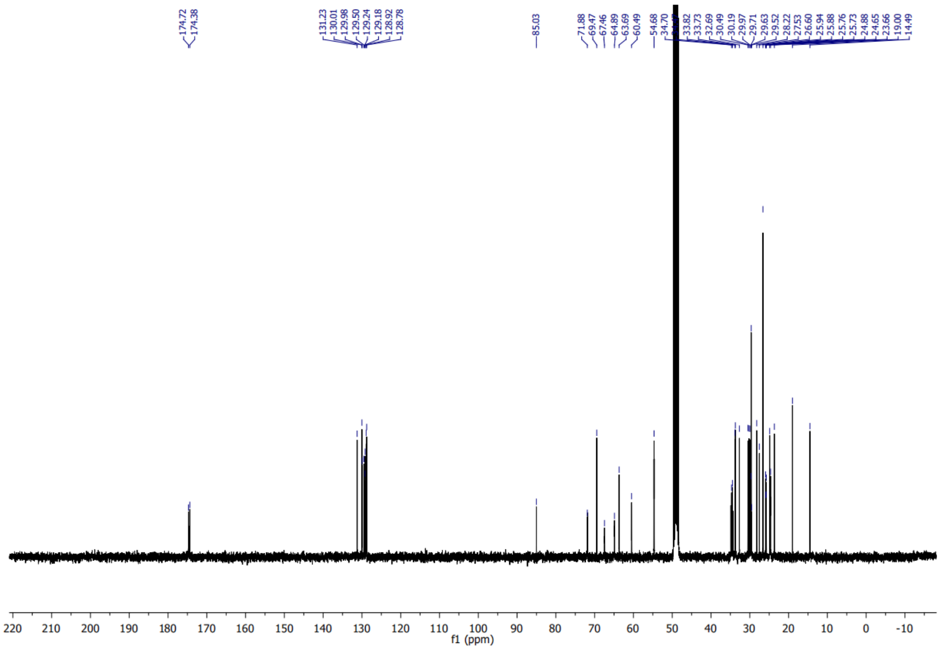

#### *Zoom of* $^{13}$*C spectrum of (15d)*

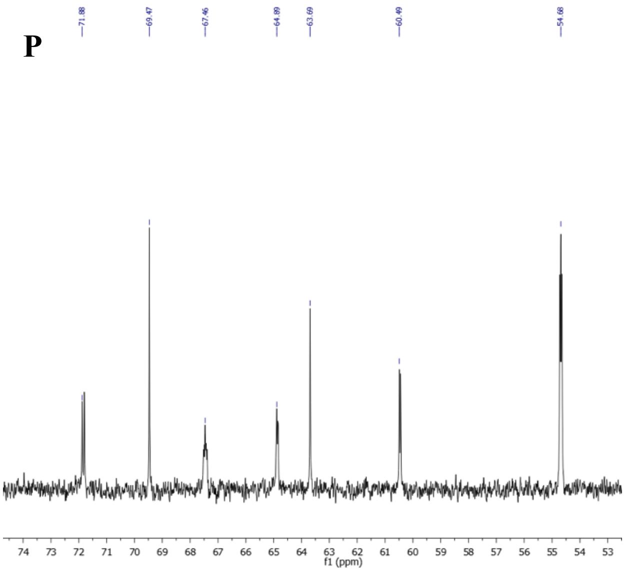

#### $^{31}$*P spectrum of (15d)*

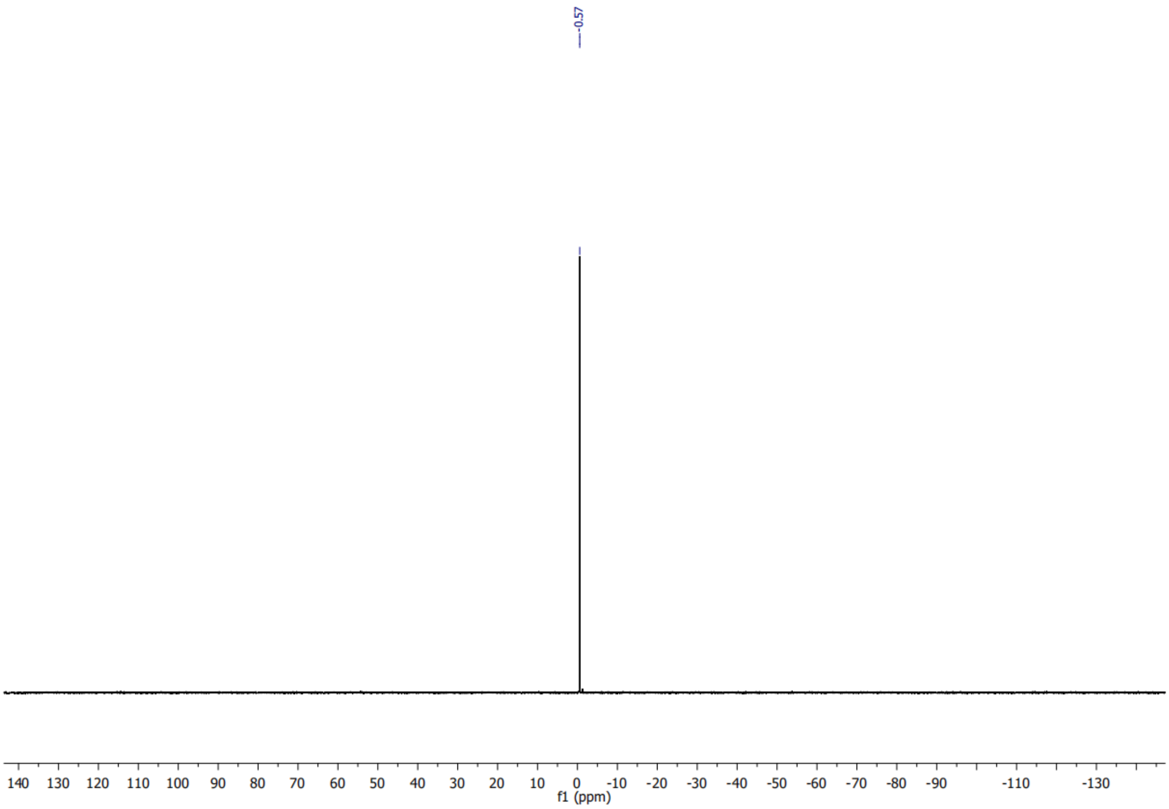

### Sphingomyelin/Y (16)

#### *[1]H Spectrum of (16)*

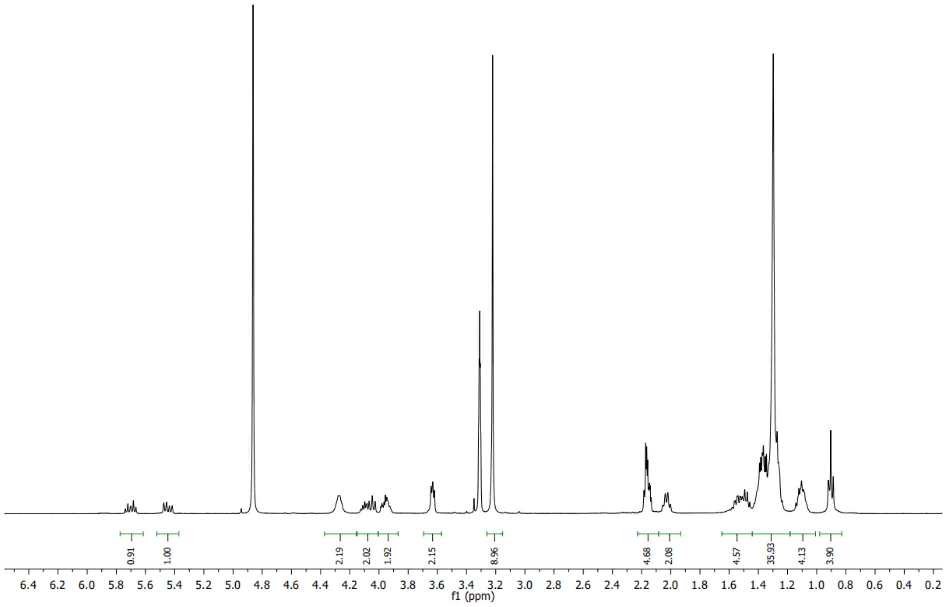

#### *[13]C spectrum of (16)*

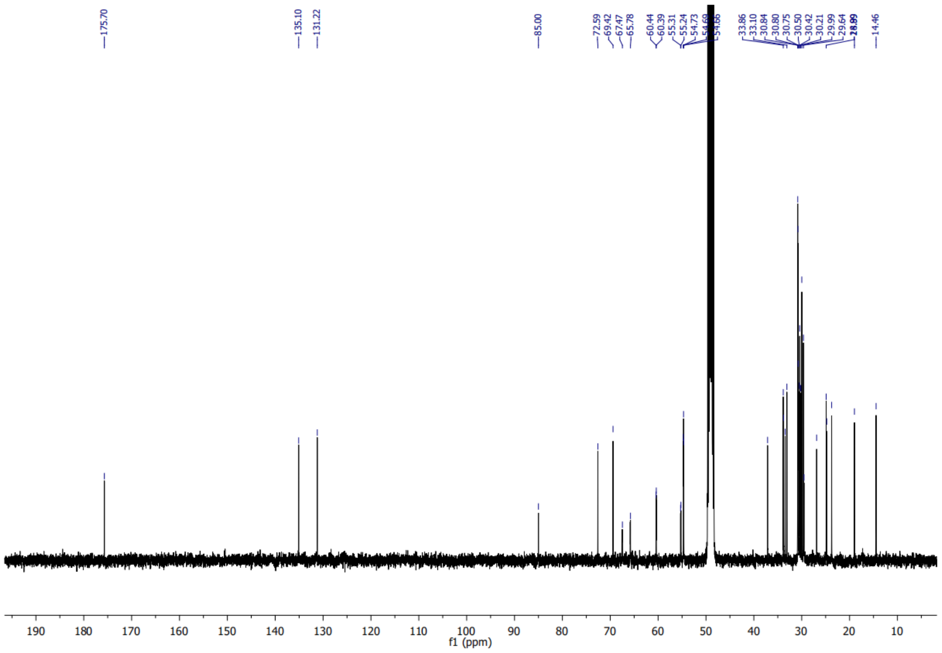

#### *Zoom of $^{13}C$ spectrum (16)*

Q

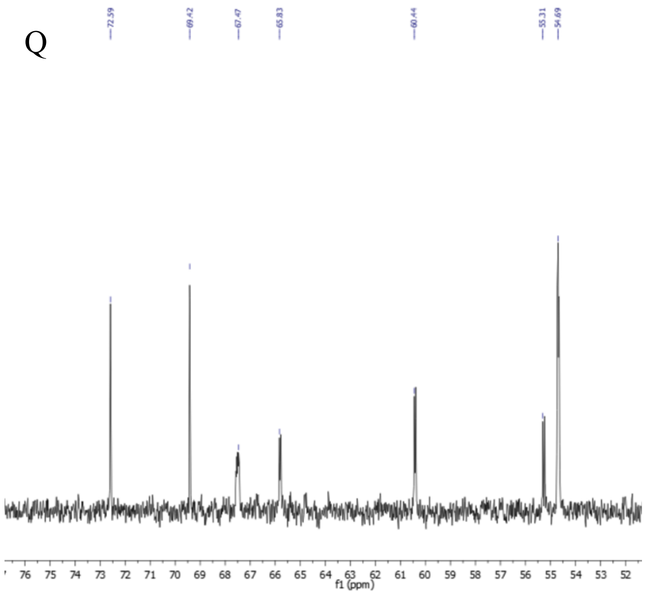

#### *$^{31}P$ spectrum of (16)*

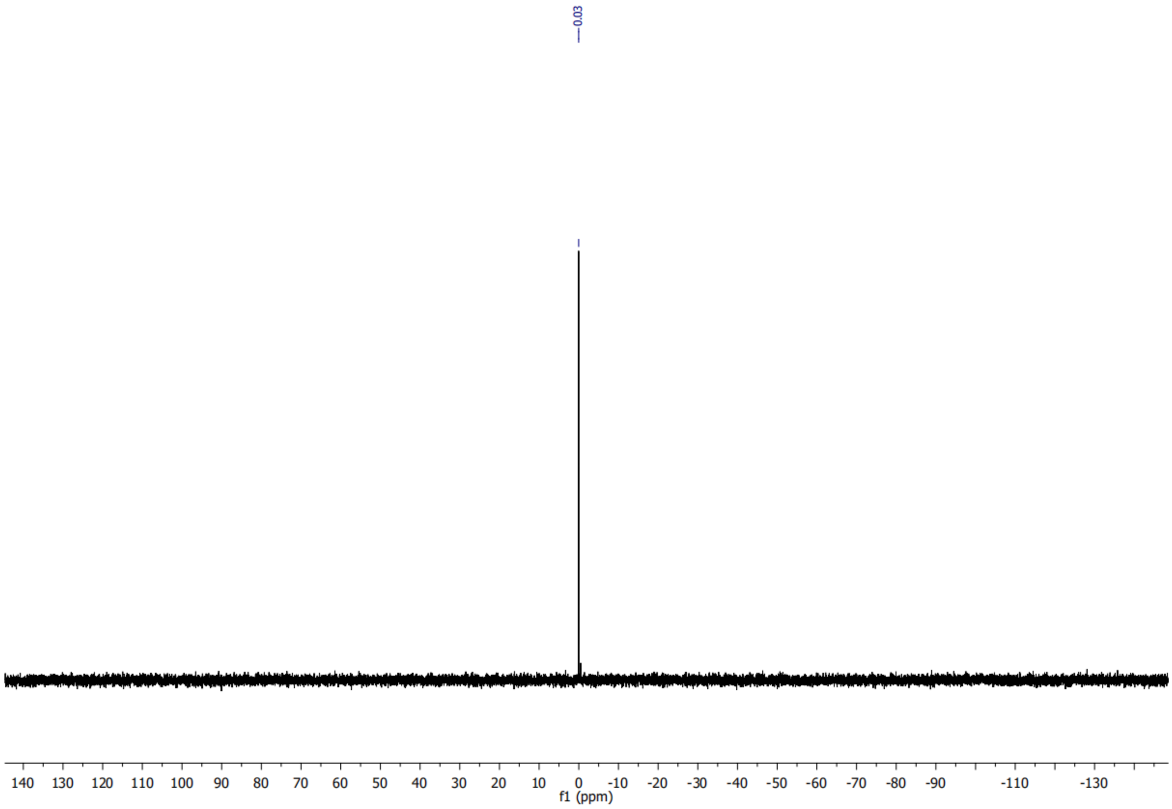

#### (R)-2-((6-(3-(non-8-yn-1-yl)-3H-diazirin-3-yl)hexanoyl)oxy)-3-((triethyl-silyl)oxy)propyl oleate (18)

##### *$^{1}$H Spectrum of (18)*

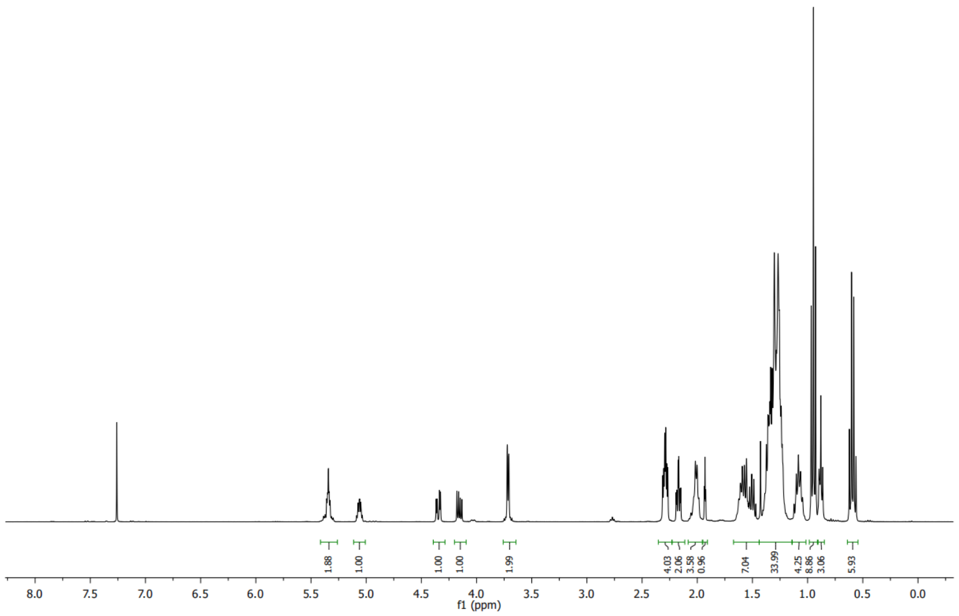

##### *$^{13}$C spectrum of (18)*

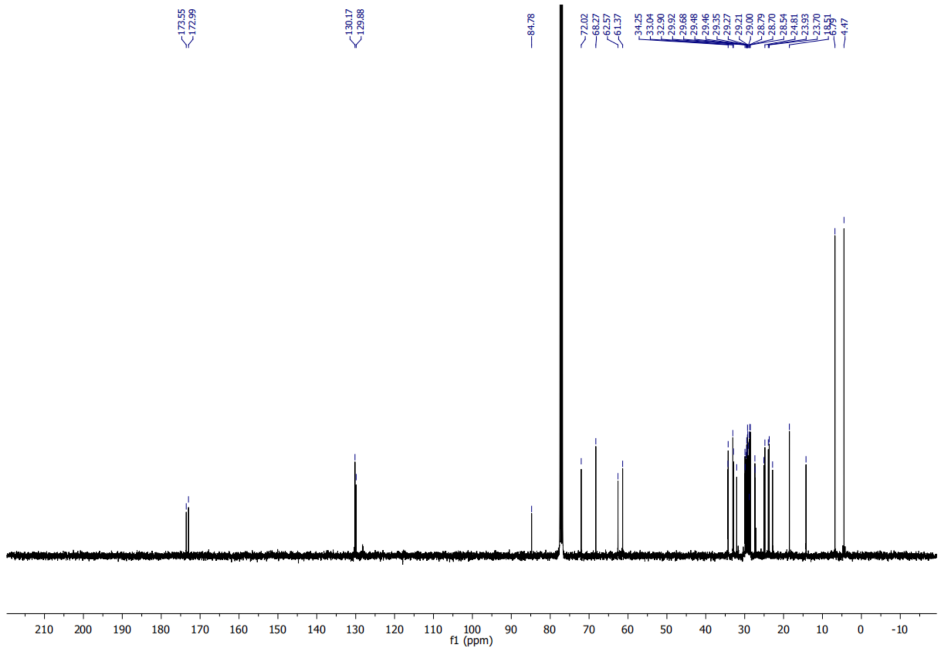

#### (S)-3-hydroxy-2-((6-(3-(non-8-yn-1-yl)-3H-diazirin-3-yl)hexanoyl)oxy) propyl oleate (19)

##### *$^{1}$H spectrum of (19)*

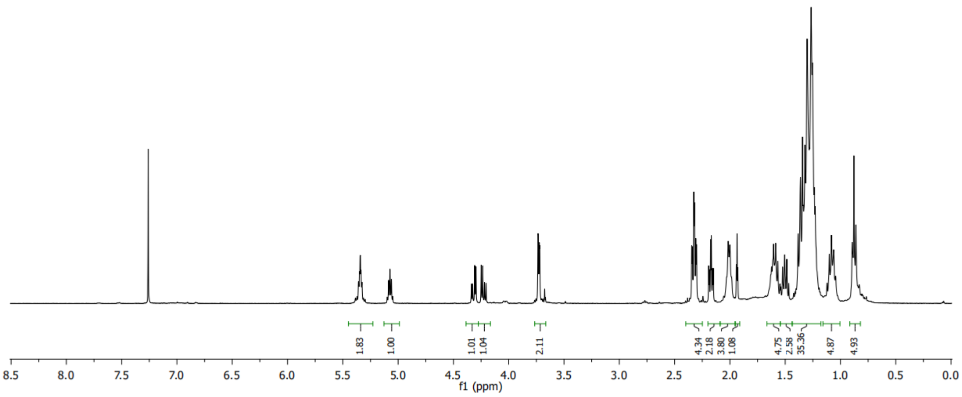

##### *$^{13}$C spectrum of (19)*

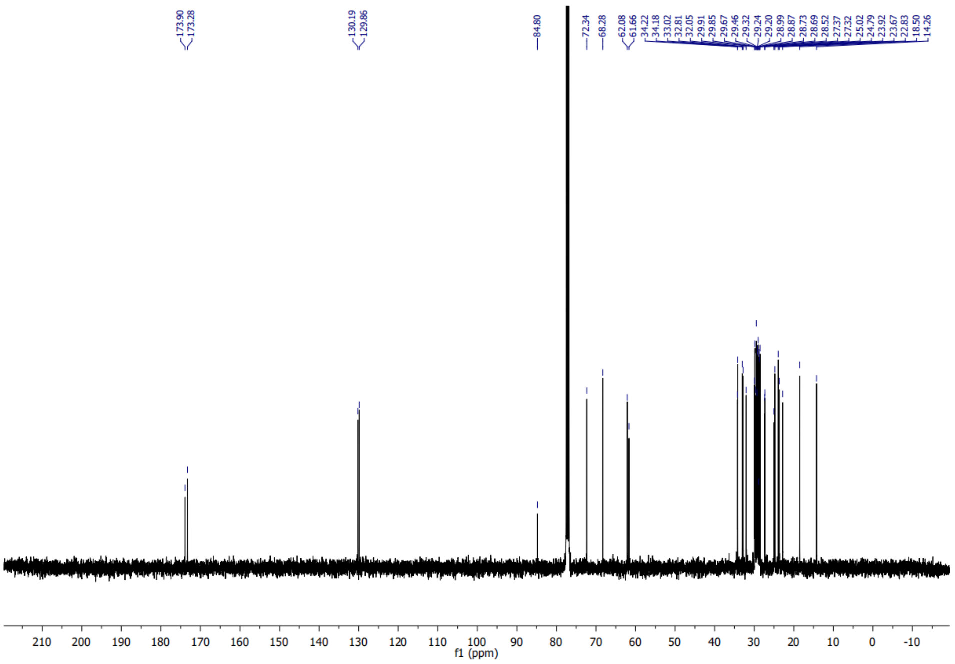

#### (R)-3-((di-tert-butoxyphosphoryl)oxy)-2-((6-(3-(non-8-yn-1-yl)-3H-diazirin-3-yl)hexanoyl)oxy)propyl oleate (21)

##### *$^{1}$H Spectrum of (21)*

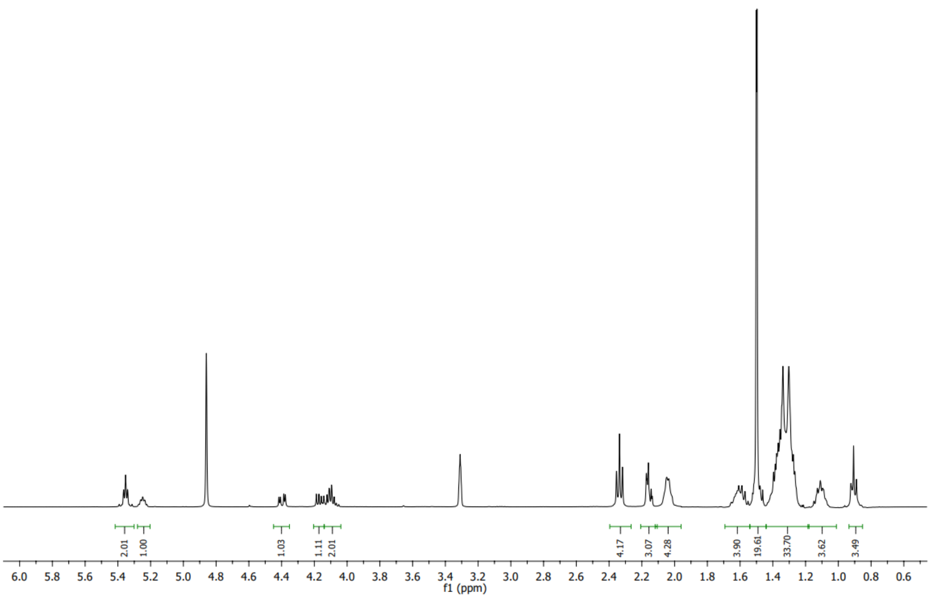

##### *$^{13}$C spectrum of (21)*

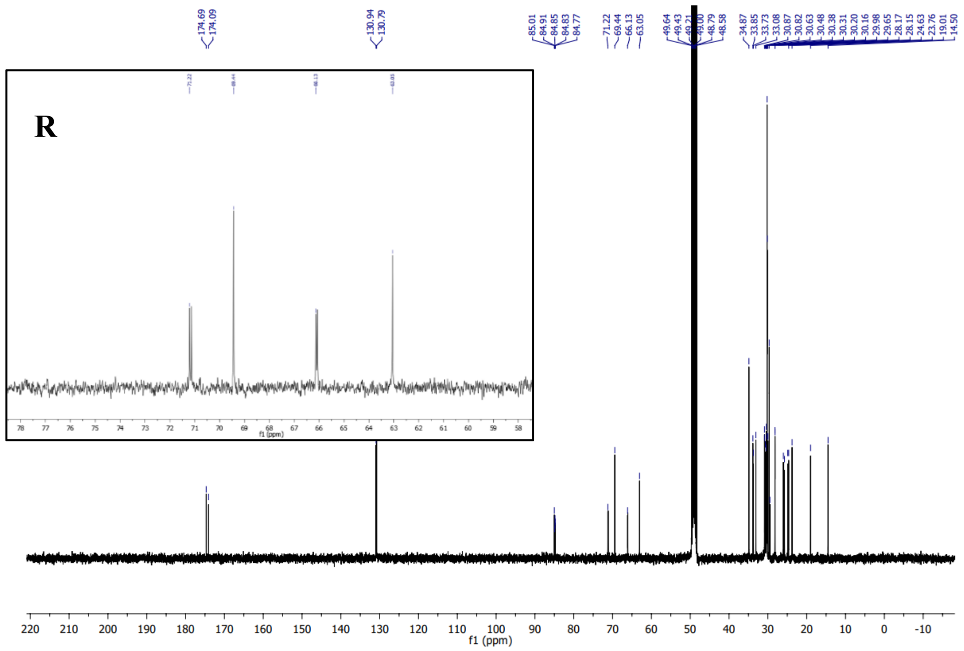

#### ***$^{31}$P spectrum of (21)***

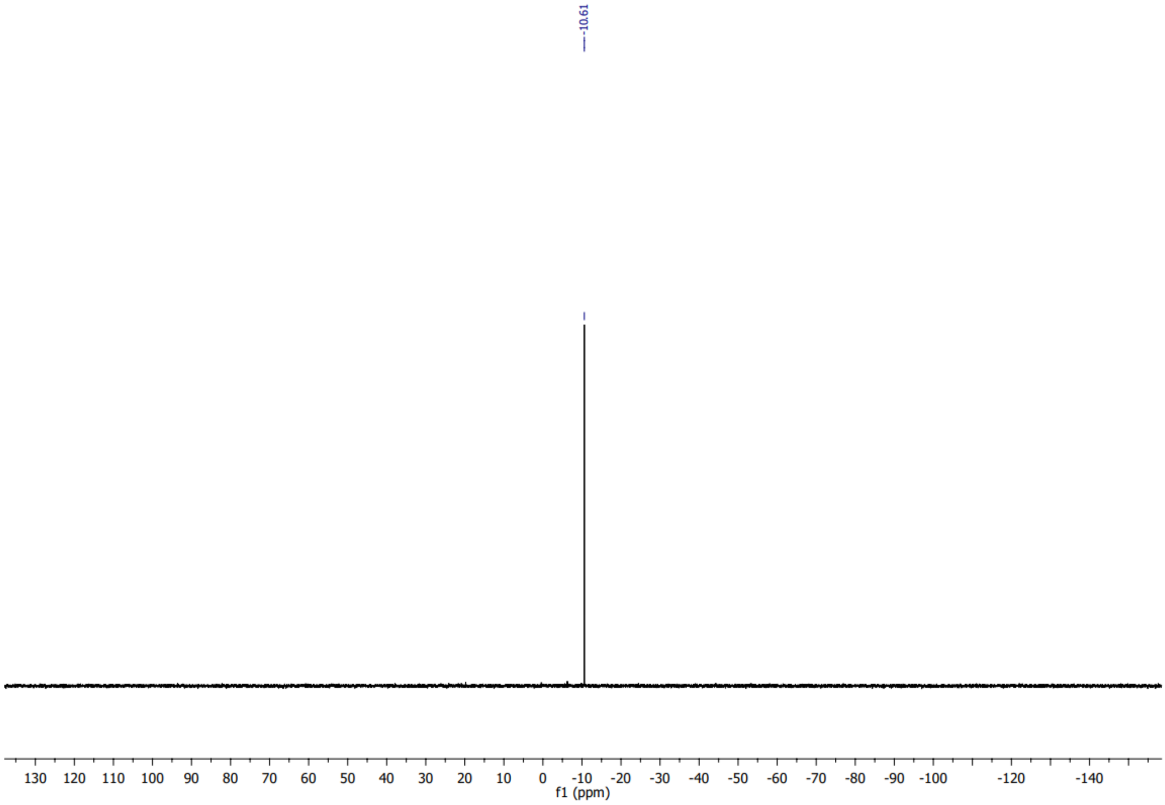

# PA(18:1/Y) (22)

#### *[1]H spectrum of (22)*

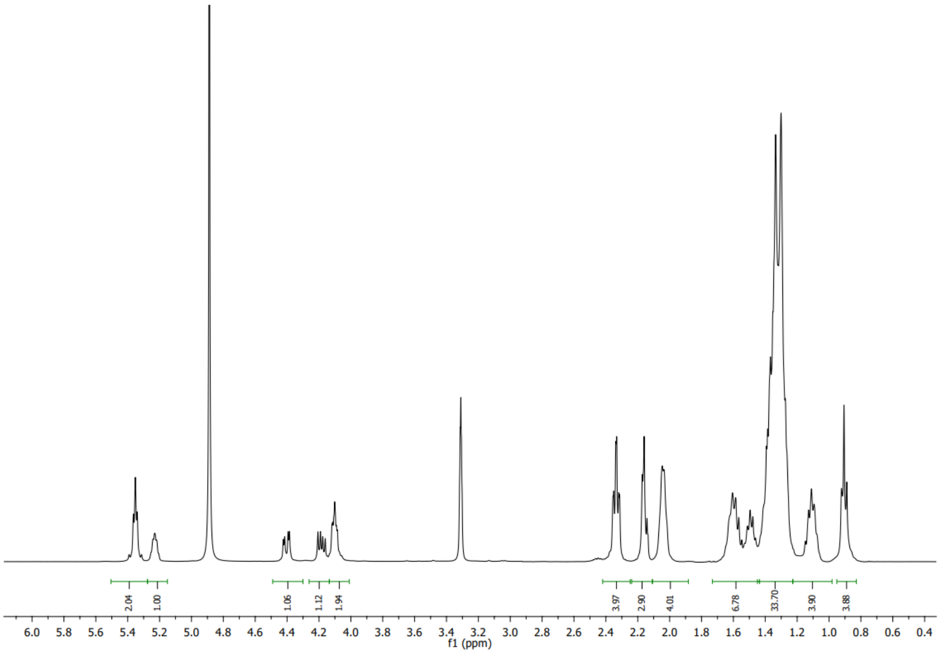

#### *[13]C spectrum of (22)*

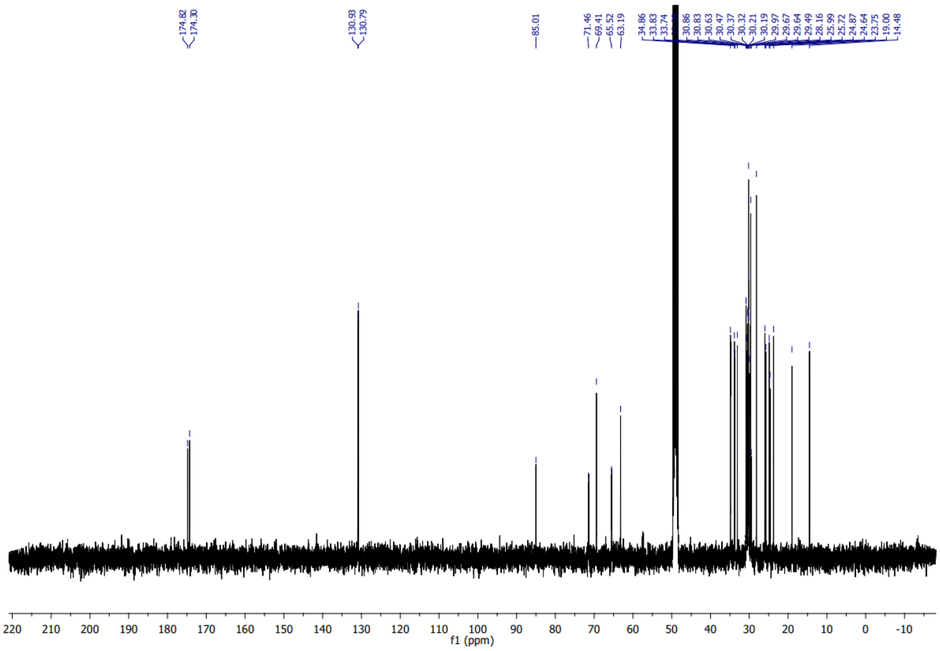

#### *Zoom of $^{13}C$ spectrum of (22)*

S

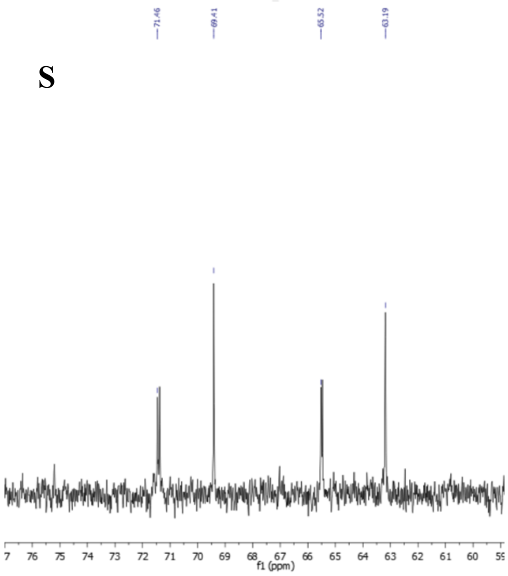

#### *$^{31}P$ spectrum of (22)*

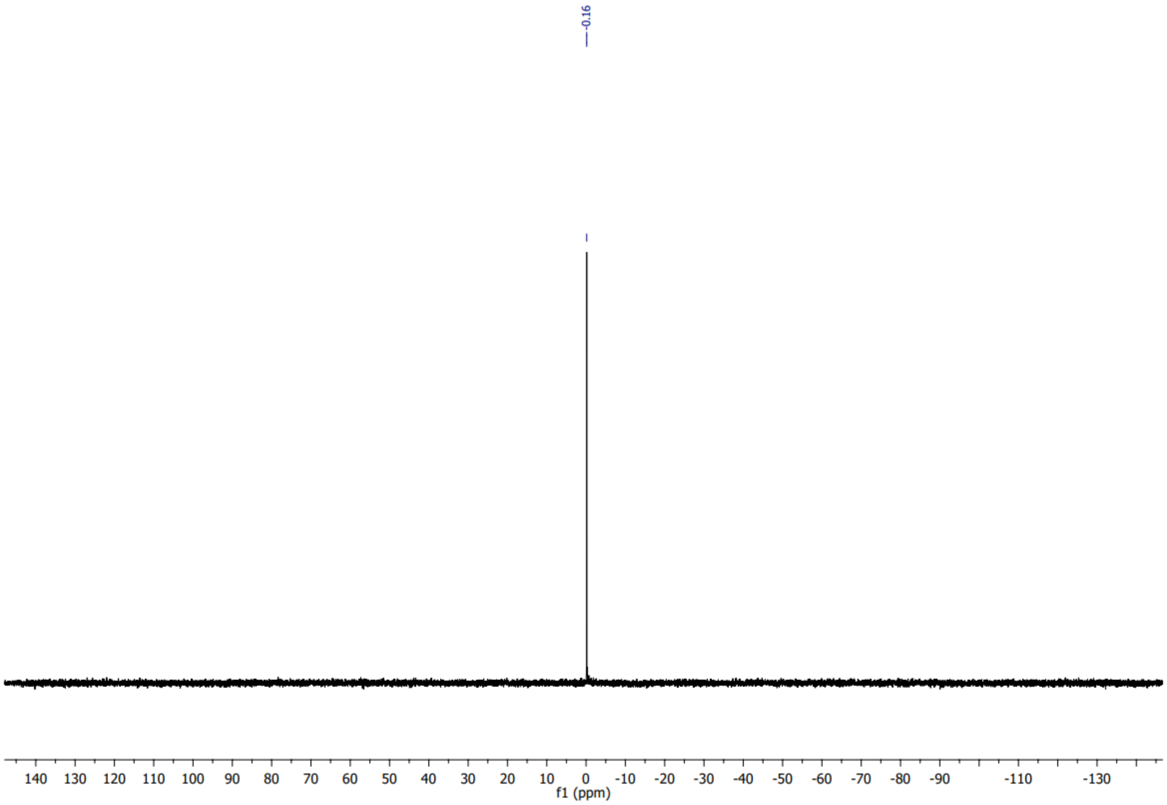

# PC(18:1/16:0[$^{13}C_{16}$]) (23)

#### *$^{1}$H spectrum of (23)*

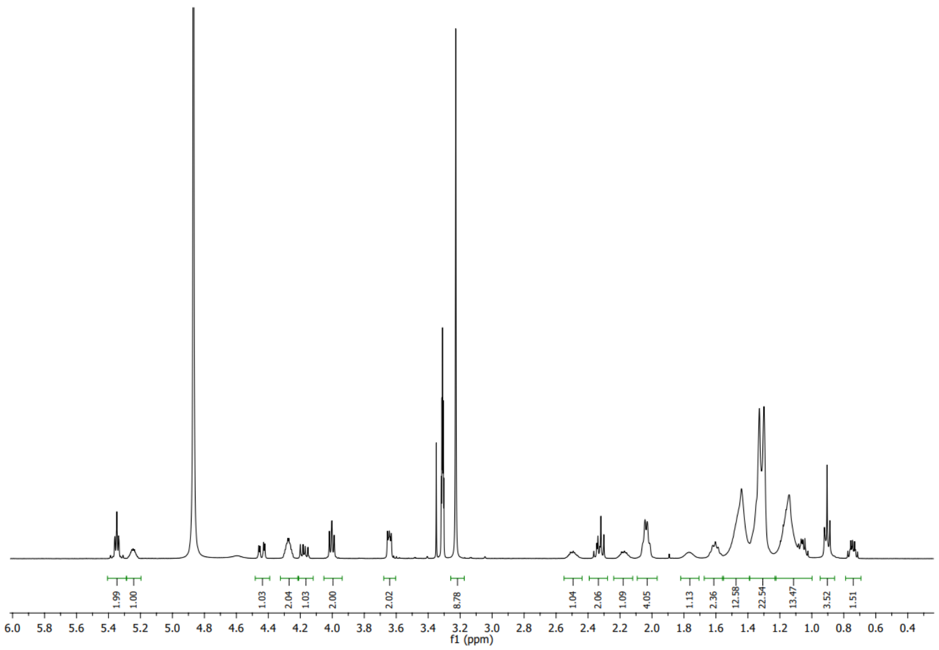

#### *$^{13}$C spectrum of (23)*

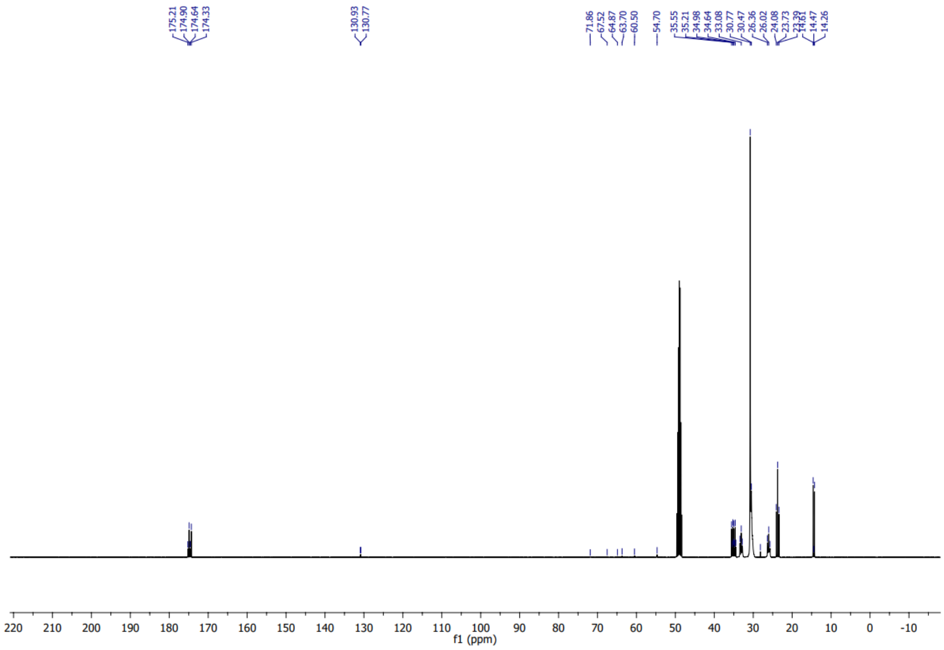

#### *Zoom of $^{13}$C spectrum of (23)*

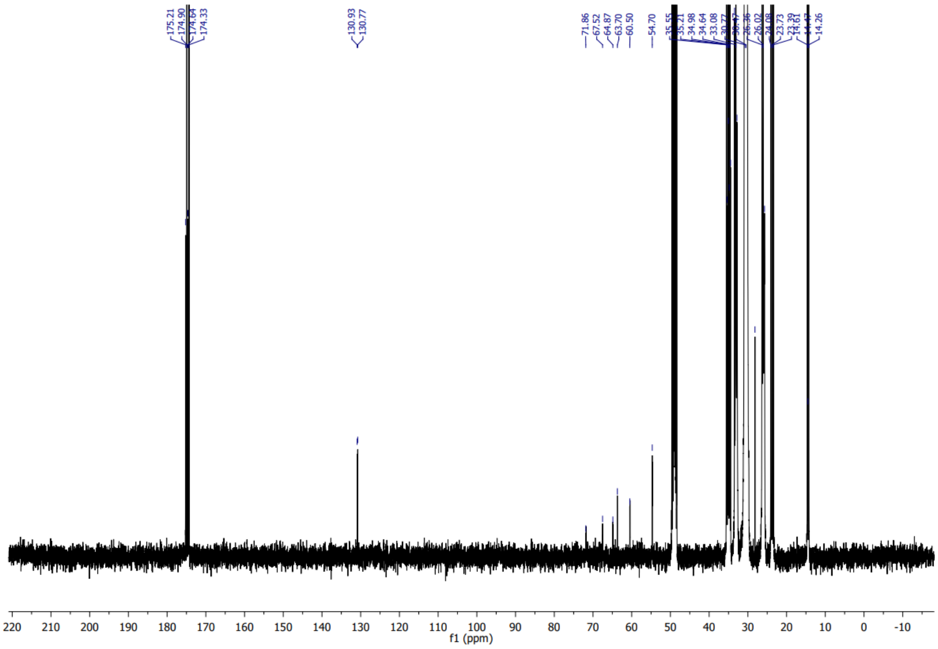

#### *$^{31}$P spectrum of (23)*

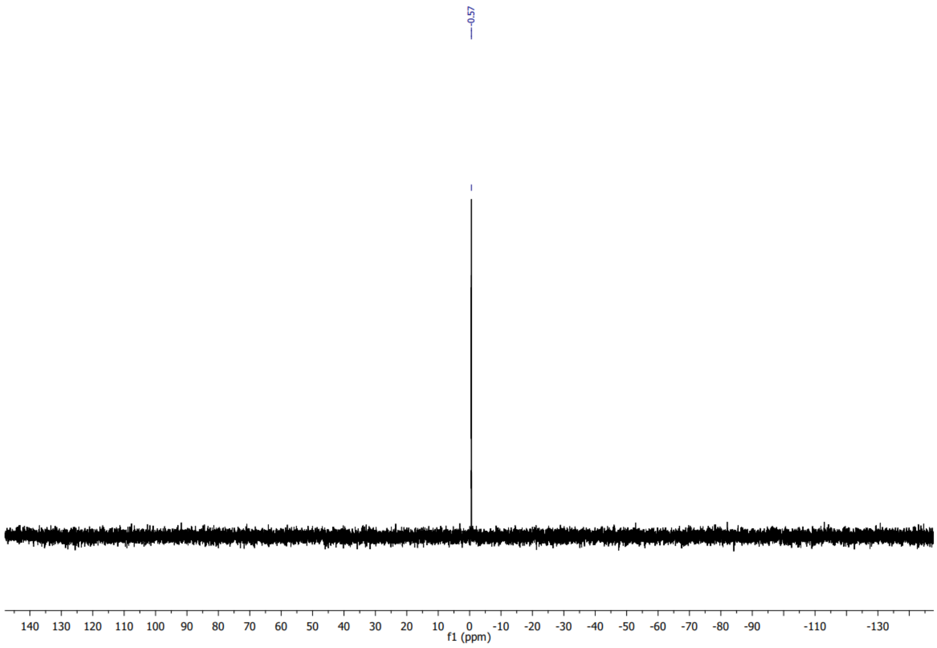

### References

1. Lorent, J. H. *et al.* Plasma membranes are asymmetric in lipid unsaturation, packing and protein shape. *Nat Chem Biol* **16**, 644–652 (2020).

2. Zhou, H., Huo, Y., Yang, N. & Wei, T. Phosphatidic acid: from biophysical properties to diverse functions. *The FEBS Journal* **291**, 1870–1885 (2024).

3. Bonsergent, E. *et al.* Quantitative characterization of extracellular vesicle uptake and content delivery within mammalian cells. *Nat Commun* **12**, 1864 (2021).

4. Rilla, K. Diverse plasma membrane protrusions act as platforms for extracellular vesicle shedding. *J of Extracellular Vesicle* **10**, e12148 (2021).

5. Hanson, P. I. & Cashikar, A. Multivesicular Body Morphogenesis. *Annu. Rev. Cell Dev. Biol.* **28**, 337–362 (2012).

6. Marushchak, D., Gretskaya, N., Mikhalyov, I. & Johansson, L. B.-Å. Self-aggregation – an intrinsic property of G $_{M1}$ in lipid bilayers. *Molecular Membrane Biology* **24**, 102–112 (2007).

7. Angelova, M. I. & Dimitrov, D. S. Liposome electroformation. *Faraday Discuss. Chem. Soc.* **81**, 303 (1986).

8. Vinklárek, I. S. *et al.* Experimental Evidence of the Existence of Interleaflet Coupled Nanodomains: An MC-FRET Study. *J. Phys. Chem. Lett.* **10**, 2024–2030 (2019).

9. Van Rossum, G. & Drake, F. L. *Python 3 Reference Manual*. (CreateSpace, Scotts Valley, CA, 2009).

10. Berg, S. *et al.* ilastik: interactive machine learning for (bio)image analysis. *Nat Methods* **16**, 1226–1232 (2019).

11. Van Der Walt, S. *et al.* scikit-image: image processing in Python. *PeerJ* **2**, e453 (2014).

12. Virtanen, P. *et al.* SciPy 1.0: fundamental algorithms for scientific computing in Python. *Nat Methods* **17**, 261–272 (2020).

13. Baretton, G. The OpenCV Library. *Dr. Dobb's Journal of Software Tools* (2000).

14. Pedregosa, F. *et al.* Scikit-learn: Machine Learning in Python. *Journal of Machine Learning Research* **12**, 2825–2830 (2011).

15. Matyash, V., Liebisch, G., Kurzchalia, T. V., Shevchenko, A. & Schwudke, D. Lipid extraction by methyl-tert-butyl ether for high-throughput lipidomics. *J Lipid Res* **49**, 1137–1146 (2008).

16. Schuhmann, K. *et al.* Bottom-Up Shotgun Lipidomics by Higher Energy Collisional Dissociation on LTQ Orbitrap Mass Spectrometers. *Anal. Chem.* **83**, 5480–5487 (2011).

17. Grgic, A. *et al.* Ultrahigh-Mass Resolution Mass Spectrometry Imaging with an Orbitrap Externally Coupled to a High-Performance Data Acquisition System. *Anal Chem* **96**, 794–801 (2024).

18. Schuhmann, K. *et al.* Monitoring Membrane Lipidome Turnover by Metabolic 15N Labeling and Shotgun Ultra-High-Resolution Orbitrap Fourier Transform Mass Spectrometry. *Anal Chem* **89**, 12857–12865 (2017).

19. Herzog, R. *et al.* LipidXplorer: a software for consensual cross-platform lipidomics. *PLoS One* **7**, e29851 (2012).

20. Wu, C. F. J. Jackknife, Bootstrap and Other Resampling Methods in Regression Analysis. *Ann. Statist.* **14**, (1986).

21. Drobot, B. *et al.* Cm $^{3+}$ /Eu $^{3+}$ induced structural, mechanistic and functional implications for calmodulin. *Phys. Chem. Chem. Phys.* **21**, 21213–21222 (2019).

22. Spinella, A., Caruso, T. & Coluccini, C. First total synthesis of natural aplyolides B and D, ichthyotoxic macrolides isolated from the skin of the marine mollusk Aplysia depilans. *Tetrahedron Letters* **43**, 1681–1683 (2002).

23. Schuhmann, K. *et al.* Quantitative Fragmentation Model for Bottom-Up Shotgun Lipidomics. *Anal. Chem.* **91**, 12085–12093 (2019).

24. Schuhmacher, M. *et al.* Live-cell lipid biochemistry reveals a role of diacylglycerol side-chain composition for cellular lipid dynamics and protein affinities. *Proc. Natl. Acad. Sci. U.S.A.* **117**, 7729–7738 (2020).
